# TGF-beta coordinates changes in Keratin gene expression during complex tissue regeneration

**DOI:** 10.1101/2025.06.02.657343

**Authors:** Dipak D Meshram, Yuchang Liu, Molly Worth, Changqing Zhang, Tom J. Carney, Henry H. Roehl

## Abstract

Zebrafish larval tail regeneration can be broadly broken down into three phases: wound repair, initiation of regeneration and redevelopment of the tail. We and others have shown that activation of the Hedgehog and TGF-beta signalling pathways is required to initiate tail regeneration. In this study we initially focus on these two pathways to better understand their role regeneration. We have found that Hedgehog signalling acts upstream of the TGF-beta pathway by activating transcription of the TGF-beta ligand *tgfb1a*. We have identified differentially expressed genes that are regulated after tail excision as well as those that are dependent upon TGF-beta signalling. From this analysis we found that Keratin genes fall into categories of co-expression: one set that is upregulated by TGF-beta signalling after injury (*krt_up_*) and a second set that is downregulated (*krt_down_*). To see whether these two sets show co-expression more broadly, we analysed single cell RNA sequencing data and found that *krt*s fall into the same categories during zebrafish development. We have mapped the expression of a representative member of *krt_up_ (krt97*) and *krt_down_* (*krt5*) sets during development and regeneration and show that *krt97* is found in epithelia that is moving and *krt5* is expressed in epithelia that is stationary. Based upon our results we propose that the *krt_up_* set allows for collective epithelial migration while the *krt_down_* set inhibits mobility. These results reveal one mechanism by which TGF-beta may initiate regeneration and could help to explain how changes in cytoskeletal composition influence cell movement during development and repair.

## Introduction

Although mammals are able to heal wounds and regrow many tissues such as skin and muscle, the regeneration of complex structures is limited. Aquatic vertebrates on the other hand, are able to regenerate large portions of organs including limb, tail, spinal cord, retina and heart(Poss and Tanaka 2024). When comparable damage occurs in mammals, tissue usually fails to regrow and scarring occurs. One possible explanation for the poor regenerative potential observed in mammals may be a failure to recruit competent cells to the site of injury in sufficient numbers to restore the missing tissue. Thus, the characterisation of signalling pathways that are able to induce regenerative cell migration in model organisms may aid in the development of novel regenerative medicine strategies.

Zebrafish larval tail excision offers a rapid and simple model to understand the basic principles of regeneration(Fig. 1a)(Rojas-Muñoz et al. 2009; Romero et al. 2018). Excision is typically performed between 48 and 72 hours post fertilisation (hpf) and involves removal of some of the larval fin (finfold), muscle, neural tube, notochord and other tissue. Following excision, early wound signals (ATP, Calcium and ROS) are released and the wound rapidly closes (Fig. 1b)(Niethammer 2016). The body axis shortens and a ball of notochord cells forms at the stump (called the notochord bead)(Fig. 1c-d). The notochord bead expresses Hedgehog (HH) ligands which induce migration of regenerative cells to the stump (Fig. 1d-e). This results in the formation of the blastema and wound epithelium (Fig. 1f). The blastema and wound epithelium express signalling pathways which orchestrate the redevelopment of the tail (Fig. 1g). Regeneration of the tail completes by around 72 hours post excision (hpe).

**Figure 1.**
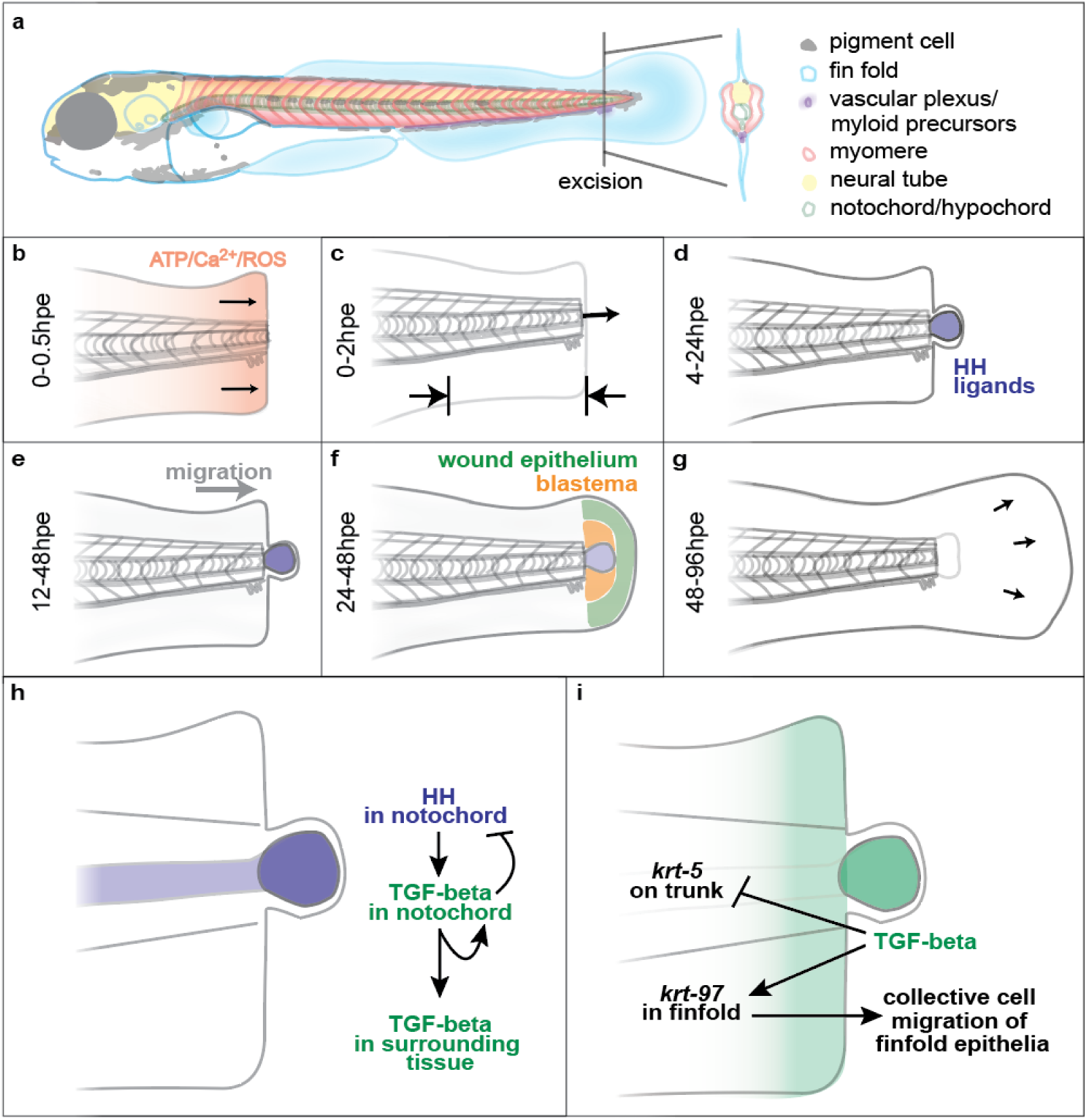
Signalling events during tail regeneration. **(a)** Larval tail excision removes the end of the zebrafish tail and includes different cell types. **(b)** Within 0.5 hours post excision (hpe) early wound signals are released and the wound is closed. **(c)** The body axis shortens and the notochord bead is extruded. **(d)** The notochord bead becomes a HH signalling centre that activates regeneration. **(e)** Regenerative cells migrate towards the stump. **(f)** The wound epithelium and blastema form. **(g)** The tail reforms. **(h and i)** Models presented in this paper.

The TGF-beta pathway plays crucial roles in wound healing in mammalian systems and is involved with wound closure, fibroblast activation, inflammation and fibrosis(Massagué and Sheppard 2023). It has also been shown to act during the regeneration of different zebrafish tissues including heart, spinal cord, lateral line and adult skin(Richardson et al. 2016; Bensimon-Brito et al. 2020; Keatinge et al. 2021; Hsu et al. 2024). Recent analysis has shown that TGF-beta signalling acts during larval tail regeneration and is linked closely with pro-regenerative metabolic reprogramming(Sinclair et al. 2021). How the TGF-beta pathway relates to the existing signalling network for tail regeneration (Fig. 1a-g) is not clear.

Keratins (Krts) are intermediate filament proteins that are found mainly in the cytoskeleton of epithelial cells and are associated with desmosomes and hemidesmosomes. They give strength to many structures including skin, hair and nails. In addition to their structural roles they have effects on diverse cellular processes such as cell signalling, metabolism and mechanotransduction(Redmond and Coulombe 2021). Studies in zebrafish have primarily used *krts* as markers of epithelia and transgenic fish carrying fluorescent proteins driven by *krt* promoters are widely used(Love, Huber, and Anders 2014; Gong et al. 2002; Lee, Asharani, and Carney 2014).

In this study we have propose that the TGF-beta pathway plays a central role during tail regeneration. We find that it is activated by notochord-derived HH signalling through the upregulation of transcription of the ligand *tgfb1a* in the notochord bead (Fig. 1h). Notochordal TGF-beta signalling then inhibits HH signalling by repressing transcription of *indian hedgehog a* (*ihha*) in the notochord and further activates *tgfb1a* in the notochord and surrounding tissue (Fig. 1i). To better understand the downstream effects of TGF-beta signalling we use bulk RNA-sequencing (RNA-seq) to identify differentially expressed genes (DEGs) after tail excision and after pharmacological inhibition of TGF-beta signalling. We identify members of the *krt* gene family as targets of the TGF-beta pathway after excision, and find that they fall into two categories of co-expression (synexpression): one set that is upregulated by TGF-beta signalling after injury (*krt_up_*) and a second set that is downregulated by TGF-beta signalling after injury (*krt_down_*). Next we focus on two *krt* genes, *krt97* and *krt5* as representative members of *krt_up_* and *krt_down_* to examine their expression during development and regeneration. We find that whereas *krt97* is expressed in epithelia that are actively moving, *krt5* is expressed in epithelium that is stationary. Using developmental single cell RNA sequencing (scRNA-seq) datasets we show that genes in *krt_up_* and *krt_down_* tend not to be co-expressed suggesting that these two groups have distinct functions during development. Collectively these data suggest a model where TGF-beta allows for collective cell movement in the epithelia by modulating expression of different members of the *krt* gene family.

## Results

### TGF-beta signalling acts downstream of HH signalling

Pharmacological inhibition of the HH and TGF-beta signalling pathways have striking similarities: In both cases the wound closes and the notochord bead forms, but there is no recruitment of regenerative cells to the stump(Sinclair et al. 2021; Romero et al. 2018). We have previously shown that HH ligands *ihhb* and *shha* are expressed in the notochord both before and after tail excision. To begin to understand the relationship between the HH and TGF-beta pathways we used standard RNA in situ methods(Thisse and Thisse 2014) to identify *tgfb1a* as a candidate ligand gene that is expressed in the notochord bead immediately after wounding (Fig. 2a). Expression is initially just in the notochord and then spreads to the surrounding tissue. To determine whether the two pathways regulate each other we treated fish with either a HH pathway inhibitor (cyclopamine) or a TGF-beta pathway inhibitor (SB431542). We found that cyclopamine treatment blocks *tgfb1a* transcriptional activation (Fig. 2b) and SB431542 treatment increases *ihha* expression (Fig. 2c). These two findings indicate that TGF-beta signalling is likely to act downstream of the HH pathway and forms a negative feedback loop to modulate HH activity. In addition we found that TGF-beta activates its own transcription especially in the tissue surrounding the notochord bead (Fig. 2d). We have previously found that while the standard whole-mount RNA in situ methods detect expression in the notochord bead, it does not detect genes expressed in the intact notochord(Romero et al. 2018). To investigate whether *tgfb1a* is expressed in notochord cells prior to injury, we switched to the hybridization chain reaction fluorescent (HCR) in situ method which utilises much smaller probes(Choi, Schwarzkopf, and Pierce 2020). Whereas, *ihhb* is expressed through-out the notochord before and after injury, *tgfb1a* is only activated in the notochord bead upon injury (Fig. 2e). Together these findings suggest a model whereby the formation of the notochord bead causes the induction of *tgfb1a* expression by HH signalling.

**Figure 2.**
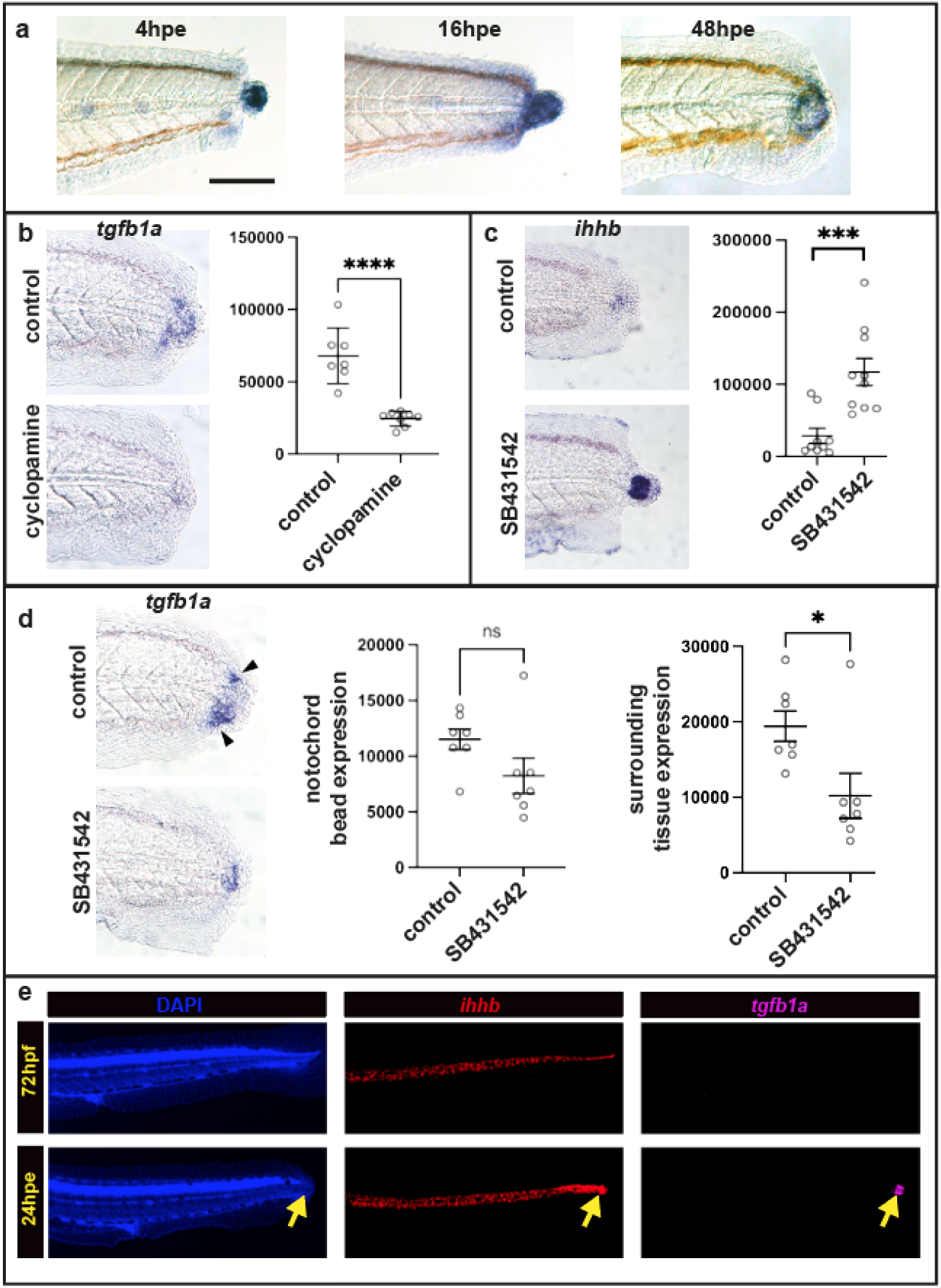
*tgfb1a* expression is activated by Hedgehog signalling in the notochord bead. **(a)** Expression of *tgfb1a* at three time points after tail excision. Expression is initially confined to the notochord, but spreads to surrounding tissue over time. Scale bar 100um **(b)** Cyclopamine treatment from −2 to 24hpe results in a significant reduction in *tgfb1a* expression. **(c)** SB431542 treatment from −2 to 24hpe results in a significant increase in *ihhb* expression. **(d)** SB431542 treatment from 4 to 24hpe results in a significant decrease in *tgfb1a* expression in the tissue that surrounds the notochord bead (arrow heads). **(e)** HCR in situs of an uninjured fish at 72hpf (top row) and injured fish at 24hpe (bottom row). In uninjured fish *ihhb* is expressed throughout the notochord prior to excision and *tgfb1a* is absent. After excision *tgfb1a* expression is activated only in the notochord bead (yellow arrows). Units for the graphs in b-d are thresholded pixels (see methods for details). Asterisks indicate significance (* = 0.05; ** = 0.01; *** = 0.001; **** = 0.0001) results using the unpaired t test and ns is not shown to be significant.SB431542 concentration was 50uM and cyclopamine concentration was 20uM.

### Transcriptional analysis of TGF-beta signalling during regeneration

To begin to understand the role of TGF-beta signalling in regenerative cell recruitment, we performed bulk RNA-seq to identify DEGs after tail excision (Fig. 3a). To focus on genes that are regulated at the start of regeneration, fish were processed at 18 hours post excision (hpe). RNA was collected using four treatments: DMSO (vehicle control) unoperated, DMSO operated, SB431542 unoperated and SB431542 operated (Fig. 3b). Three replicates were done for each treatment and the data was processed using the DESeq2 package in R(Choi, Schwarzkopf, and Pierce 2020; Love, Huber, and Anders 2014). One conclusion that can be drawn from this analysis is that DEGs that are found in the SB431542 operated versus DMSO operated comparisons tend to also be DEGs in the SB431542 unoperated versus DMSO unoperated set (Fig. 3c). This suggests that genes that are targets of TGF-beta signalling during regeneration are also responding to TGF-beta activity in uninjured fish. This is unexpected as treatment with SB431542 in unoperated fish confers no visible changes. There are however some DEGs that are regulated by SB431542 in operated fish that do not show significant regulation by SB431542 in unoperated fish. These genes are potential wound/regeneration specific targets of the TGF-beta pathway (see Supplemental Fig. 1).

**Figure 3.**
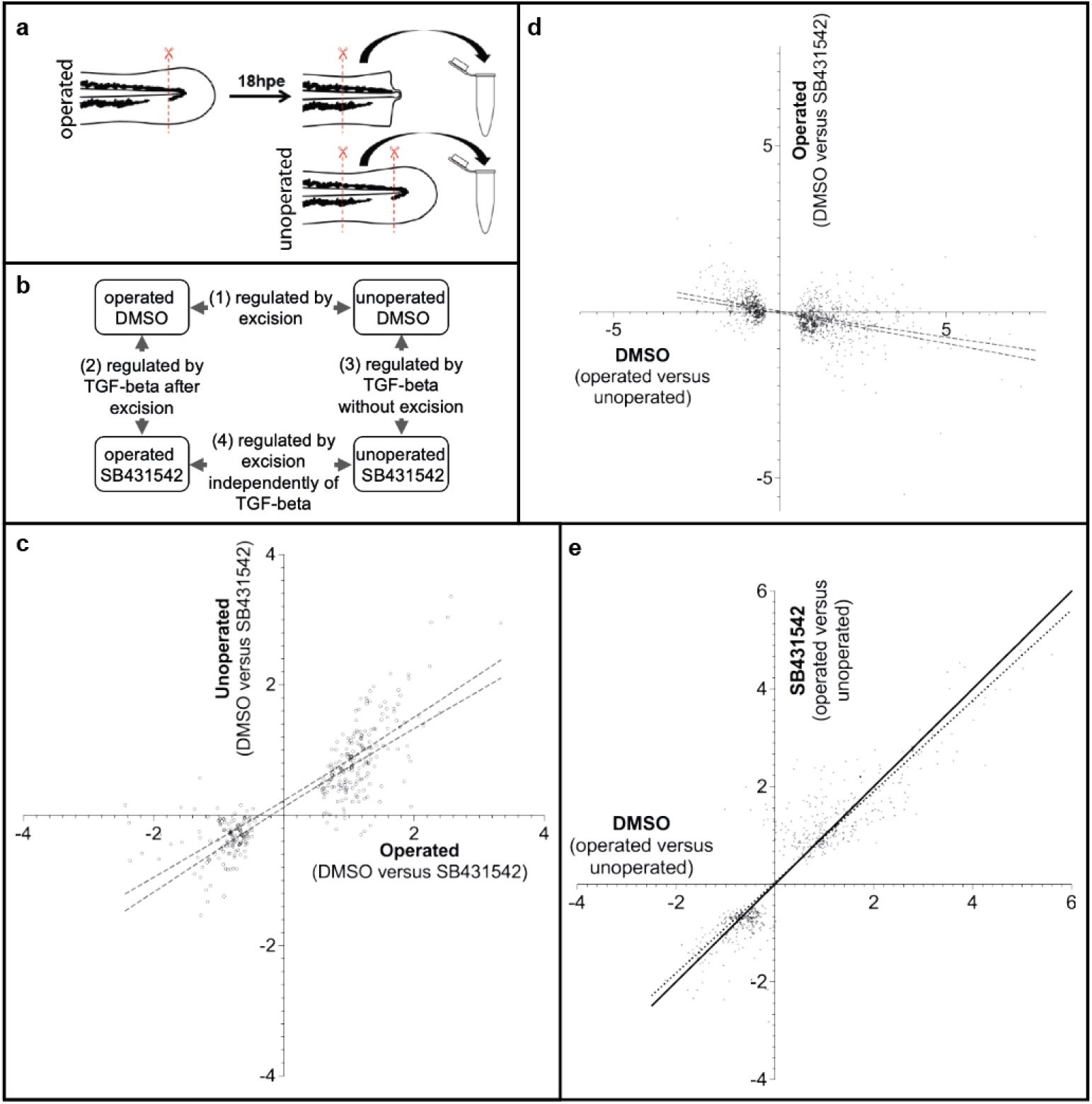
Transcriptome changes due to injury and TGF-beta activation. **(a)** Preparation of samples. Operated samples have approximately 200um of posterior tail tissue removed and are incubated until 18hpe. A second 200um tissue is then removed and processed for sequencing. The unoperated samples take a similar region of the tail for processing. **(b)** Four-way comparisons are made between the samples. **(c)** Comparison of potential TGF-beta pathway targets after injury to potential TGF-beta pathway targets without injury. The X-axis is SB431542 operated compared to DMSO operated (2 in panel b) and the Y-axis is SB431542 unoperated compared to DMSO unoperated (3 in panel b). Values are in Log2. Only DEGs that have an adjusted P-value (Padj) value <0.05 in the SB431542 operated compared to DMSO operated set are plotted. DEGs in the upper right quadrant are genes that are potentially suppressed by TGF-beta signalling in both operated and unoperated samples. DEGs in the lower left quadrant are those that are potentially activated by TGF-beta signalling in both operated and unoperated samples. The lines indicate the regression with 95% confidence. The regression slope is 0.62 indicating that regulation is stronger in the injured dataset. **(d)** Comparison of potential DEGs after injury to potential TGF-beta pathway targets after injury. The X-axis is DMSO operated compared to unoperated (1 in panel b) and the Y-axis is SB431542 operated compared to DMSO operated (2 in panel b). Values are in Log2. Only DEGs that have a Padj value <0.05 in the operated compared to unoperated set are plotted. DEGs in the upper left quadrant are genes that are potentially suppressed by TGF-beta signalling following injury. DEGs in the lower right quadrant are those that are potentially activated by TGF-beta signalling after injury. The lines indicate the regression with 95% confidence. **(e)** Comparison of potential targets after injury to potential TGF-beta pathway targets after injury. The X-axis is DMSO operated compared to unoperated (1 in panel b) and the Y-axis is SB431542 operated compared to SB431542 unoperated (4 in panel b). Values are in Log2. Only DEGs that have a Padj value <0.05 in the SB431542 operated compared to SB431542 unoperated set are plotted. The solid line represents a slope of one, and the dotted line represents the linear regression.

A second conclusion from the RNA-seq comparisons is that genes that are regulated during regeneration are usually influenced by SB431542 treatment (Fig. 3d). For example, DEGs that are upregulated following injury are likely to also be downregulated when injured fish are treated with SB431542. This supports the model that TGF-beta is pro-regenerative and is required during the onset of regeneration to recruit cells. However, there are some DEGs that do not fit this model (see upper right and lower left quadrants in Fig.3d). For example DEGs in the upper right quadrant of Fig 3d are upregulated following injury and are upregulated when injured fish are treated with SB431542. It is not immediately clear why this is, but one possible explanation is that there is negative feedback affecting these genes and that testing at earlier time points would find that they are initially downregulated by TGF-beta. The most significant DEGs that are identified by this comparison are presented in Supplemental Fig. 2.

Further evidence that TGF-beta has a pro-regenerative role comes from comparison of DEGs after injury in DMSO to DEGs after injury in fish treated with SB431542 (Fig. 3e). In this graph, DEGs that are unaffected by the presence of SB431542 after injury fall on a slope of 1. The regression slope is closer to 0.9 indicating that there is a bias towards DEGs that are injury specific and are responding to TGF-beta. For example, genes that fall below the slope of 1 line in the upper right quadrant are predicted to be activated by TGF-beta whilst those that fall above the slope of 1 line in the lower left quadrant are predicted to be repressed by TGF-beta during regeneration. A subset of these genes are presented in Supplemental Fig. 3.

To find genes that are injury-specific but not TGF-beta targets we can take DEGs that fall close to the slope of 1 in Fig 3e. We can also identify this type of gene by selecting DEGs that lie close to the X-axis in Fig 3d. By applying both criteria we identified a subset of DEGs that are potential TGF-beta independent regeneration genes (Supplemental Fig. 4).

### Members of the Keratin family show TGF-beta-dependent responses to injury

To learn more about coordinated gene expression during regeneration we searched for families of genes in the RNA-seq data (Supplemental Fig. 5). Of these, the Krt family stands out because most members are regulated by TGF-beta and these members are either activated during regeneration (*krt_up_*) or inhibited (*krt_down_*) (Fig. 4a). We grouped the *krts* as being in these synexpression groups based upon being significant (P < 0.05) in at least one of the comparisons. There are nine *krts* that do not fall into these groupings. Two of these, *krt18a.2* and *krt95*, were not identified in our analysis. Of the remaining seven, six are expressed at low levels at the time of our assay which perhaps affects our detection, and the seventh (*krt222*) is only distantly related to other members of the *krt* family.

**Figure 4.**
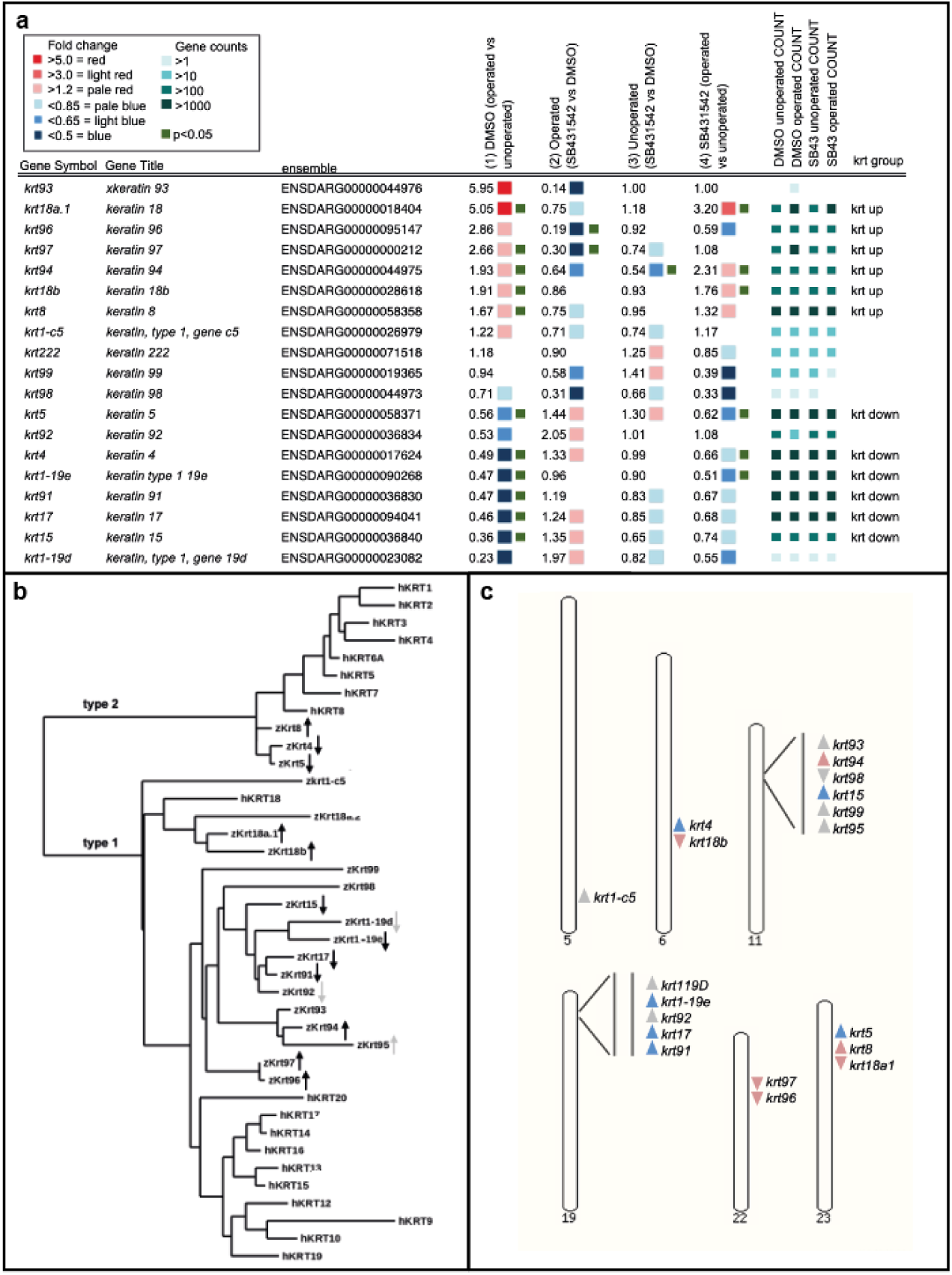
Zebrafish *krt* gene regulation and relatedness. **(a)** A table with zebrafish *krt* genes, their ensemble identifiers, the fold level of change in all four comparisons, the adjusted counts for each sample and their allocation into *krt_up_* or *krt_down_* groups. **(b)** Molecular phylogeny dendrogram with zebrafish (zKrt) and human (hKRT) representatives of the Keratin family. The arrows indicate upregulation or downregulation as shown in panel a. Black arrows indicate significance (P < 0.05) and grey arrows indicate a trend that was not significant. Phylogram tree and alignments were done with “one click” settings www.phylogeny.fr (Dereeper et al. 2008) **(c)** The genomic locations of the zebrafish *krts* with an arrowhead showing their chromosomal orientation and colour showing upregulation (pink) or downregulation (blue) by TGF-beta. The genome assembly for this analysis was GRCz11 (www.ensemble.org).

The zebrafish *krt* family is smaller than the mammalian family and is less well studied(Krushna Padhi, Akimenko, and Ekker 2006; Ho et al. 2022). To begin to understand the significance of these two groupings, we first performed phylogenetic analysis to see whether members of Krt_up_ or Krt_down_ are more closely related to each other based upon their predicted protein sequence (Fig. 4 b). We have included members of the human Krt (hKRT) family in this analysis for comparison. There are three type II Krts with one Krt_up_ and two Krt_down_ members. These fall next to the human type II Krts in one cluster. The type I branch has greater variation with one of the human Krts (hKRT18) showing more similarity to zebrafish genes than to other human Krts. Within the type I branch there is some correlation between expression and relatedness. Krt_up_ and Krt_down_ appear to separate between Krt15 and Krt94. Intriguingly, the members that show regulation but a P value that is greater than 0.5 do still show a correlation in their relatedness. For example, although *krt1-c5* does not show significance in pairwise comparisons, it does show the expected changes in expression for Krt_up_ genes (Fig 4a) and falls into the Krt_up_ phylogenetic branch. Likewise, *krt1-19d* has the expected changes in expression for Krt_down_ and falls into the Krt_down_ branch. The correlation in the type I group between expression and relatedness may indicate that there are structural differences between these Krt_up_ and Krt_down_ members and that the two groups were formed by one evolutionarily event.

To see whether this correlation extends to their chromosomal organisation, we looked at the genomic organisation of *krts* (Fig. 4c). *krts* are found clustered on six chromosomes. The cluster on chromosome 19 contains up to three members of *krt_down_* and the cluster on chromosome 22 has two members of *krt_up_*. Three of the other chromosomes are mixed synexpression genes and chromosome 5 has a single *krt* gene. Overall the genomic organisation does not show a strong correlation with the type of synexpression.

### *krts* have distinct domains of expression during regeneration and development

To understand the relevance of *krt* regulation at the tissue level we decided to do HCR in situ analysis. For this we have focused on *krt5* as a representative of *krt_down_* and *krt97* as a representative of *krt_up_*. We first tested the response of these at three stages, uninjured (96hpf), 24hpe(96hpf) and 48hpe(120hpf) (Fig. 5). In fish at these stages we found that *krt97* and *krt5* are expressed in distinct domains. *krt97* is primarily in epithelial cells of the finfold in a crescent shape around the end of the tail (Fig. 5a). This region of expression is removed following tail excision. *krt5* on the other hand is found in the epithelia that sits on the trunk. At 24hpe epithelial cells have gathered on the end of the stump to form the wound epithelium. At this time, *krt97* expression is upregulated in wound epithelium and *krt5* expression is reduced in the trunk epithelia nearest to the stump. By 48hpe the fin fold is reforming and krt97 is expressed in the new area of growth. On the other hand *krt5* is strongly downregulated near to the regrowing tail. Inhibition of TGF-beta with SB431542 during regeneration results in a strong increase in *krt5* and a reduction in *krt97* consistent with the RNA-seq data. To further confirm our results we probed scRNA-seq data for fish at 24hpe and 48hpe(Sinclair et al. 2021). We see that *krt97^+^* and *krt5^+^* cells are in distinct domains with *krt97^+^* cells increasing after excision and *krt5^+^* cells decreasing. Pseudo bulk RNA-seq analysis of this scRNA-seq data suggests an increase of 1.5 fold for *krt97* at 24hpe and a decrease of *krt5* expression to 0.5 at 24hpe and 0.38 at 48hpe. Together these data suggest that *krts* respond to injury in a TGF-beta dependent fashion and are expressed in distinct compartments of the skin.

**Figure 5.**
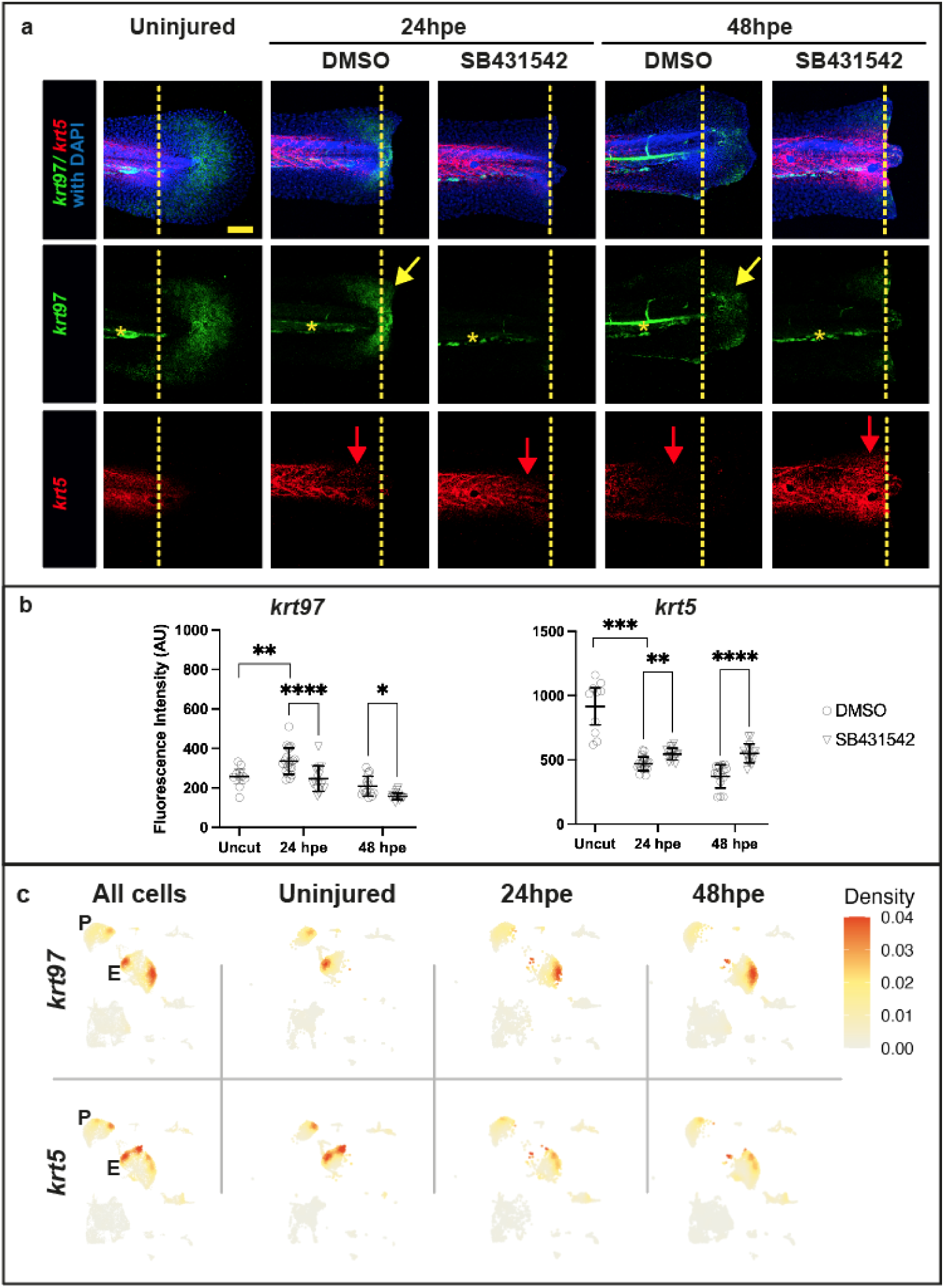
*krt97* and *krt5* are inversely regulated during tail regeneration. **(a)** In situ analysis of *krt97* and *krt5* in uninjured and injured fish treated with SB431542. Larvae were treated with DMSO or 50 M SB431542 for 2 hours prior to tail excision and kept in the chemical solution until 24 or 48 hpe. Yellow arrows indicate the re-expression of *krt97* in regenerating epithelia. Red arrows indicate the reduction of *krt5* expression on the trunk of control injured fish and the increase in expression in injured fish treated with SB431542. Yellow dotted lines indicate the excision plane. Yellow stars indicate non-specific fluorescence in the vasculature. Scale bar is 50um **(b)** Quantification of fluorescent in situs as shown in (a). Asterisks indicate significance as in Fig.2 resulting from the unpaired t test. **(c)** Expression of *krt97* and *krt5* in scRNA-seq cell clusters representing periderm (P) and epithelia (E) during regeneration.

Given that regeneration of the fin fold is likely to recapitulate aspects of fin fold development, we decided to look at tail developmental stages to compare development to regeneration. The fin fold emerges between 20hpf and 48hpf (Figure 6a). At 20hpf it can be seen as a ridge of epidermal cells that sits on the midline of the left and right sides of the tail. At this stage *krt5* is in scattered cells throughout the tail epidermis and at 24hpf and 32hpf *krt5* is expressed throughout the tail epidermis. Between 36hpf and 48hpf the fin fold goes through morphogenesis and *krt5* expression is limited to the trunk of the tail. At 20hpf *krt97* expression is in a layer of cells at the base of the epidermal ridge in a crescent of cells around the end of the tail. This expression expands and as the finfold expands it becomes more restricted to the caudal end of the finfold. Intriguingly *krt97* is not expressed in the most distal layer of cells of the finfold. To summarise, *krt5* and *krt97* are expressed in different areas of the tail epidermis with *krt97* being expressed in the fin fold as it undergoes morphogenesis.

**Figure 6.**
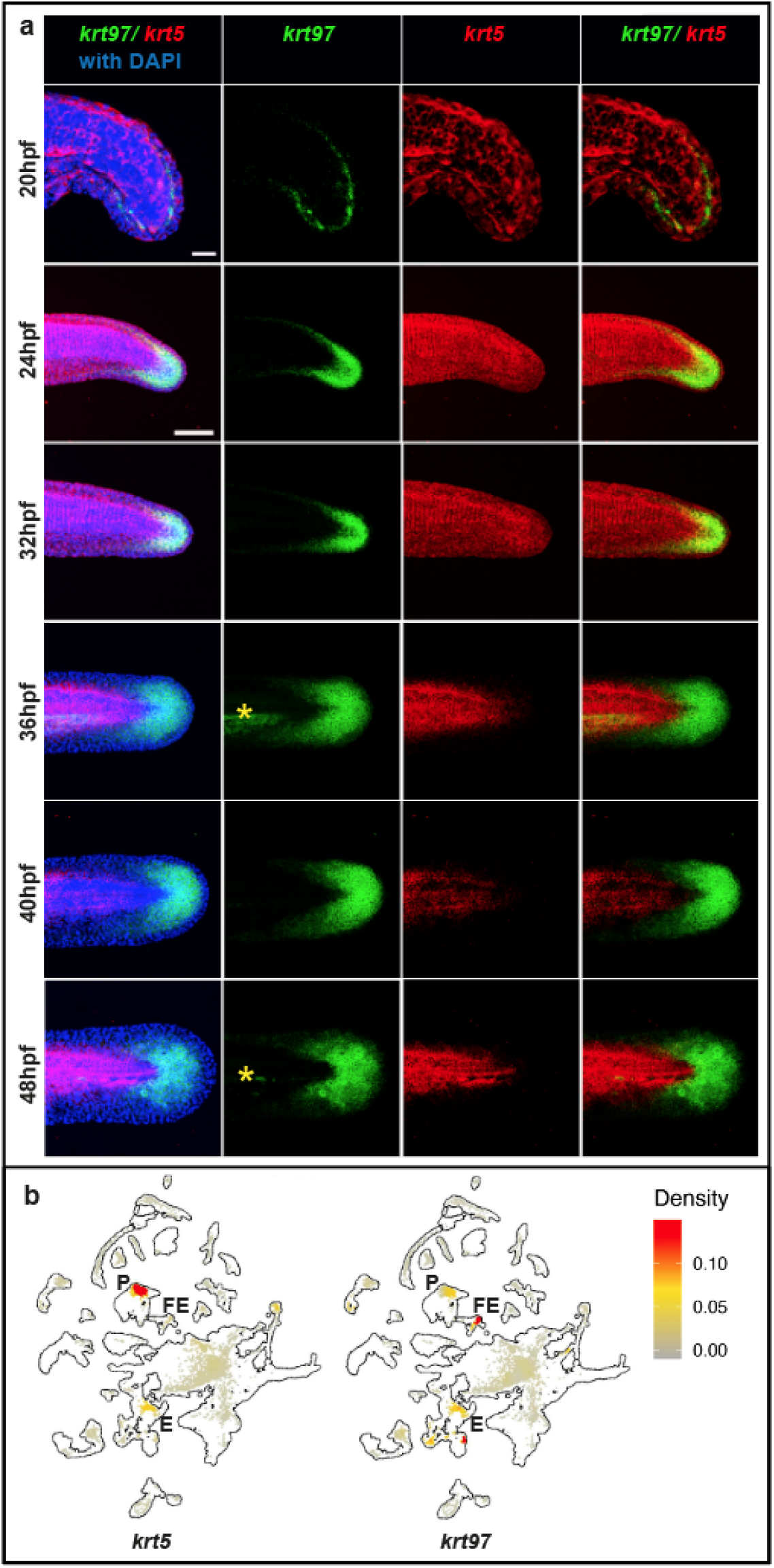
Dynamic expression of *krt97* and *krt5* during finfold development. **(a)** HCR in situ analysis of *krt97* and *krt5* between 20hpf and 48hpf. Yellow stars indicate non-specific fluorescence in the vasculature. Scale bar for 20hpf is 50um and 100um for 24hpf to 48hpf. **(b)** Expression of *krt97* and *krt5* in scRNA-seq cell clusters. The outline shows all cells in the dataset (3hpf to 120hpf) Cells in the 36-46hpf data set are shown in grey, yellow or red according to their level of expression. Clusters representing periderm (P), fin epithelium (FE) and Epithelium (E) are labelled. Data taken from Daniocell scRNA-seq dataset(Farrell et al. 2018; Sur et al. 2023).

To further understand the compartmentalisation of *krt5* and *krt97* expression we analysed the Daniocell scRNA-seq dataset which contains cells from 3hpf to 120hpf(Farrell et al. 2018; Sur et al. 2023). We found that *krt97* and *krt5* show separate domains of expression primarily between 30hpf and 58hpf and at other time points the expression domains are not distinct (data not shown). Figure 6b shows pooled cells from 36-46hpf and *krt5* is expressed in a cluster annotated as periderm and *krt97* is in fin epithelium and epithelium clusters.

Treatment with the inhibitor SB431542 during finfold development does not confer a morphological phenotype suggesting that TGF-beta signalling does not strongly affect finfold development (data not shown). However, considering our result that most genes which are regulated by TGF-beta during regeneration also show some regulation in uninjured fish, we decided to see whether SB431542 affects *krt* gene expression during development. We found that treatment between 16.5hpf to 24hpf and 40hpf to 48hpf results in a strong reduction in *krt97* expression (Fig. 7). On the other hand, *krt5* did not show a change in expression. This suggests that the regulation of *krts* during development may be related to that seen during regeneration, at least for *krt97*.

**Figure 7.**
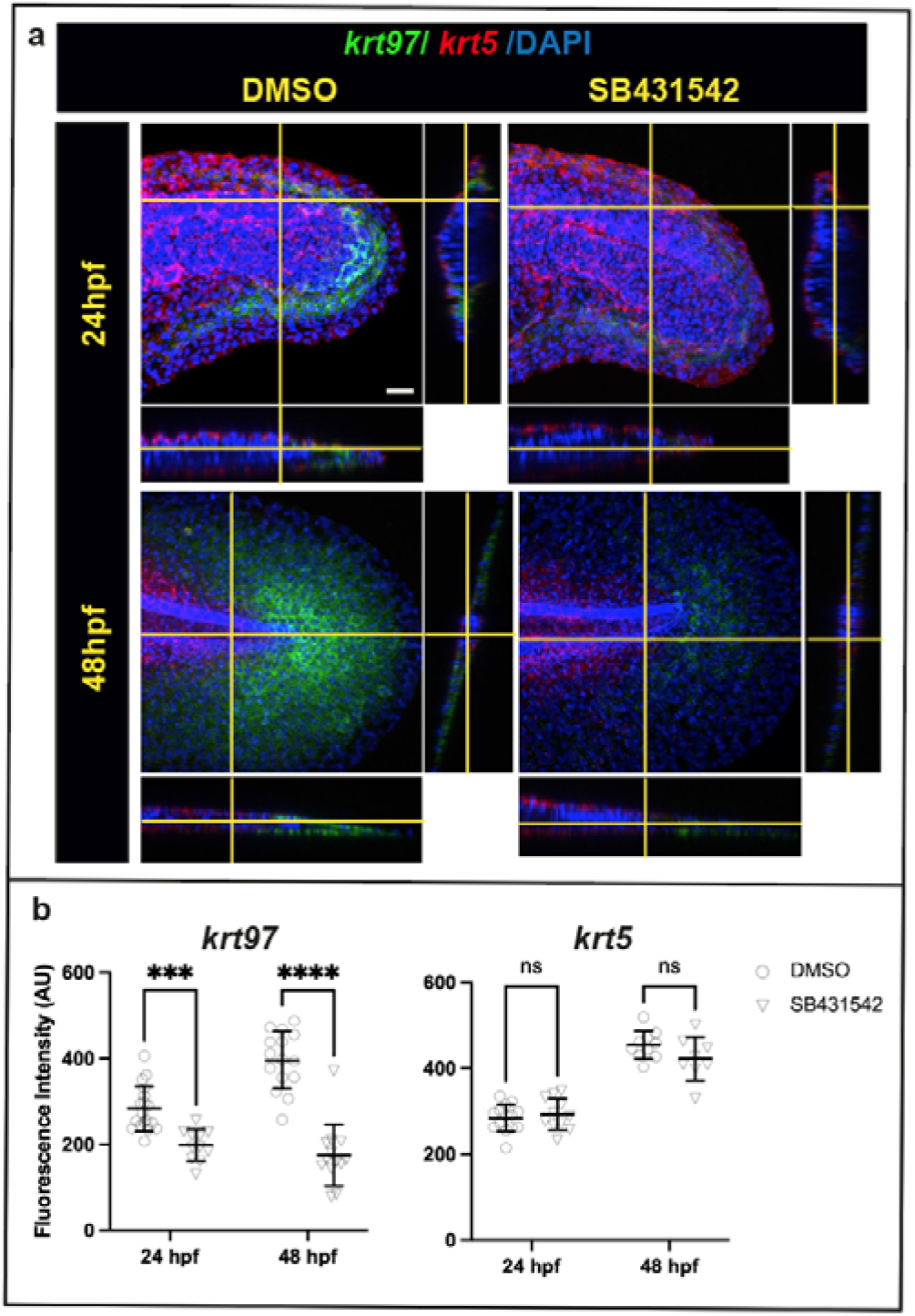
Inhibition of TGF-beta signalling affects krt97 expression during development. **(a)** HCR in situ staining for *krt97* and *krt5* with and without SB43154. Orthogonal views are shown to the right and below each image. The planes used for the orthogonal views are indicated by the yellow lines. Scale bar is 20um. **(b)** Quantification of fluorescence in (a). Asterisks indicate significance as in Fig.2 resulting from the unpaired t test. For the 24hpf timepoint, embryos at the 15 somites stages (16.5 hpf) were treated with DMSO or 100 uM SB431542 continuously. For the 48 hpf stage, larvae at 40 hpf were treated with DMSO or 100 uM SB431542.

This compartmentalised expression of *krt5* and *krt97* during finfold development and regeneration raises the possibility that other members of *krt_up_* and *krt_down_*show segregated expression patterns during development. We have tested other keratins and find that there is limited overlap between *krt_up_* and *krt_down_*gene expression (Supplemental Figure 6 and data not shown). To see whether this is the case more broadly we pooled periderm, fin epithelium and epithelium clusters from the Daniocell scRNA-seq dataset and assessed the degree of co-expression between *krt_up_* and *krt_down_* using Pearson’s correlation coefficient analysis (Fig. 8). This data shows that there is a strong degree of coexpression within each group. For example, *krt1-19e* expression is correlated to other members of *krt_down_* especially in the epidermis with *krt5* and *krt91* showing the highest association with *krt1-19e*. *krt1-19e* shows a negative correlation to members of *krt_up_* particularly with *krt18a.1*, *krt97* and *krt8* in the periderm. This data shows the co-regulation of *krts* seen during regeneration and with manipulation of TGF-beta signalling is also found more broadly in epithelia during developmental stages.

**Figure 8.**
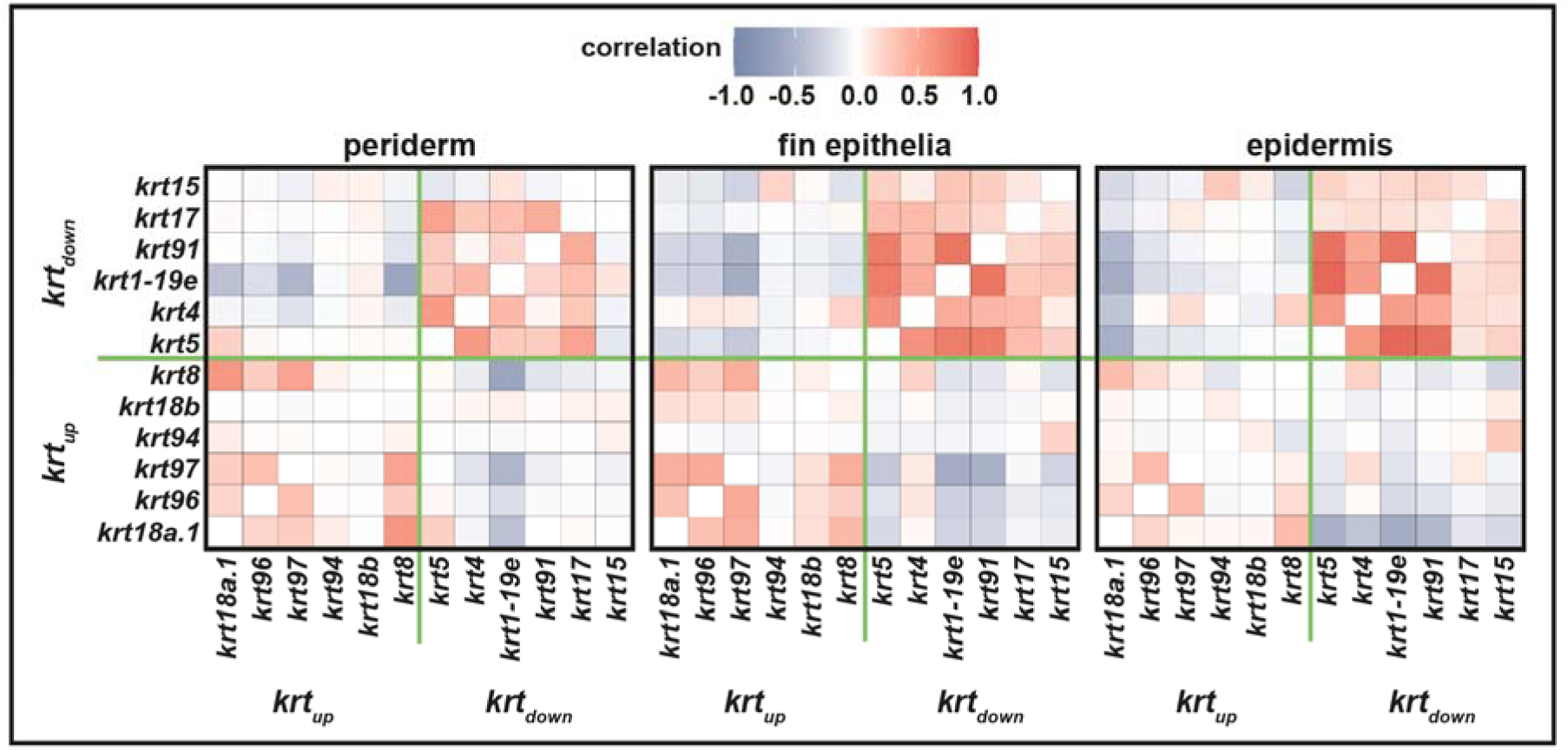
Coexpression of *krts* in skin clusters. scRNA-seq data from the Daniocell analysis (Farrell et al. 2018; Sur et al. 2023) was subsetted based upon periderm, fin epithelium and epidermis identities using all available timepoints. Each of these subsets was then tested for correlation (using Pearson Correlation in Seurat) for the *krt_down_* and *krt_up_* genes. Red colour indicates a positive correlation and blue colour indicates a negative correlation.

The expression *krt97* during finfold regeneration and development correlates to regions that are undergoing morphogenesis. These epithelia need to engage in collective cell migration and are likely to alter their cytoskeleton to allow this to occur. On the other hand, *krt5* is reduced during finfold morphogenesis and regeneration suggesting that it is inhibitory to epithelial movement. To investigate this hypothesis, we decided to analyse epithelial cell movement during development and regeneration to see whether different epithelial regions have different mobility. Our predictions were that regions that express *krt97* (epithelia of the fin fold) move more quickly than regions that express *krt5* (epithelia on the trunk). To test cell movement during regeneration we used photo-conversion of a nuclear localised dendra transgene (Fig 9a, b, g and Supplemental Fig. 7). We found that cells in the proximal finfold move the furthest averaging approximately 70um per day relative to cells on the trunk. In unoperated animals there was very little movement. To assess cell movement during development, we made timelapse recordings of a nuclear-localised fluorescent reporter and measured movement relative to the underlying somites (Movie 1). As during regeneration we found that during development cells on the finfold move faster than cells on the trunk (by approximately 15um/hour) (Fig. 9c-f, h). These results show that epithelia move at dramatically different rates during regeneration and development, and that the differences in movement may correlate to differences in *krt* expression.

**Figure 9.**
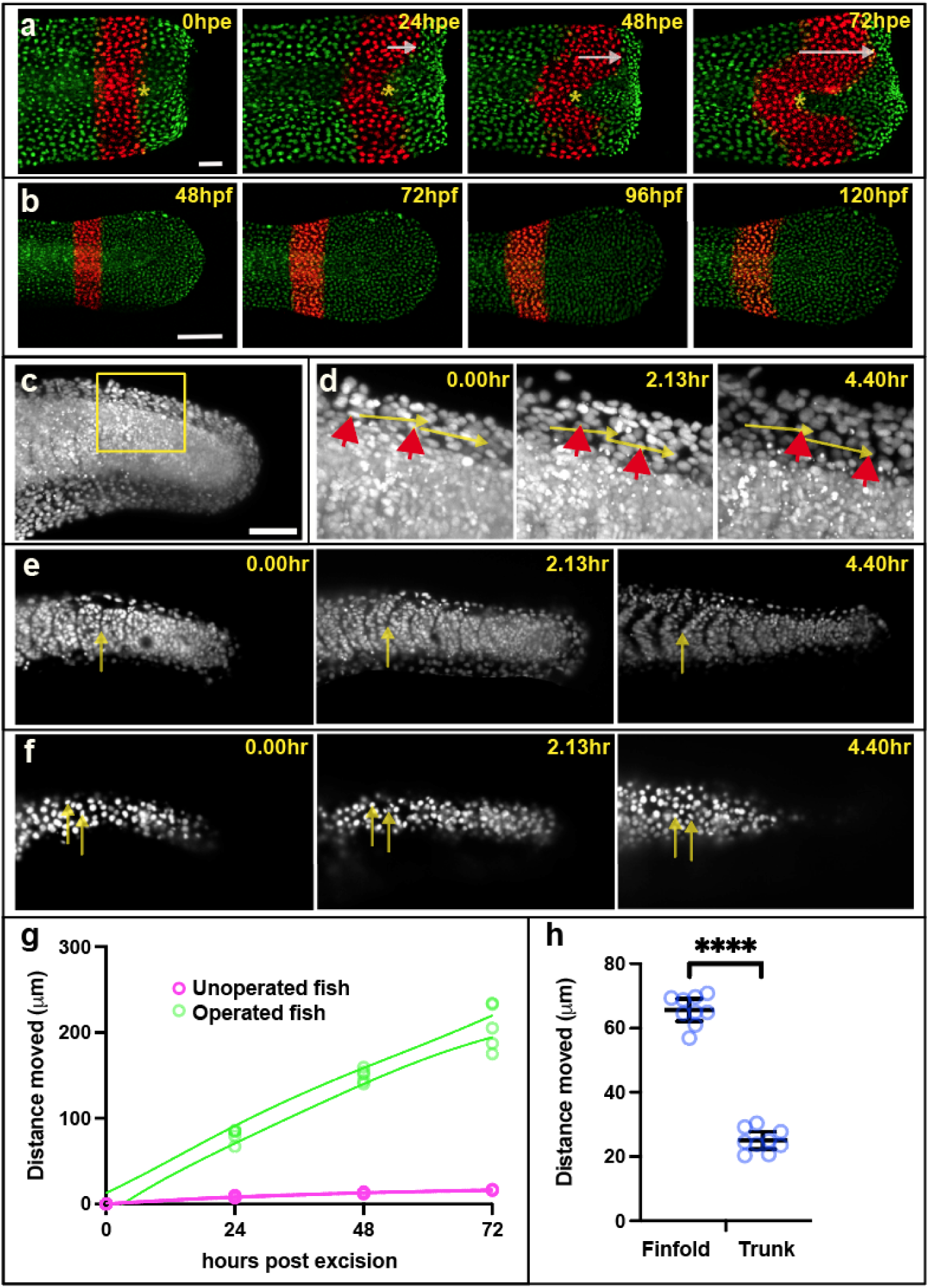
Collective migration of epithelial cells is faster in the finfold than on the trunk. **(a)** Cell nuclei were followed over a period of three days after tail excision using Dendra photoconversion (red nuclei). The asterisks mark the border of converted nuclei on the trunk and the arrows indicate the distance travelled by nuclei on the fin epithelia relative to the trunk. Scale bar = 50um **(b)** Unoperated control fish show very little movement over these four days. Scale bar = 100um **(c)** Overview image (z-projection) showing nuclei labelled red fluorescence *(actb2:h2b-mCherry)* at 24hpf. Scale bar = 100um. **(d)** Close-up of the box in (c) showing three time points during a time-lapse recording. The red arrowheads point to two finfold epithelium nuclei at the different time points. The yellow arrows indicate the distance travelled. **(e)** Sections showing somite boundary positions at three timepoints with arrows marking the same position. **(f)** Sections following epithelium nuclei on the trunk at three timepoints with arrows marking their positions. **(g)** Quantification of movement of finfold cells after excision as in panels (a) and (b). Lines indicate non-linear fit 95% confidence interval. Individual unpaired t test for each timepoint are < 0.0001. **(h)** Quantification of movement of finfold cells during development as in panels (c-f). Note that the midline somite boundary is used to standardise the epithelial cell movements. Asterisks indicate significance as in Fig.2 resulting from the unpaired t test.

## Discussion

In this study we focus on how regenerative cells are recruited to the site of injury after tail excision. We present a model whereby HH signalling activates regenerative cell recruitment by transcriptionally activating *tgfb1a* expression in the notochord (Fig. 1h). The TGF-beta pathway then induces more expression of *tgfb1a* whilst inhibiting expression of *ihhb* in the notochord. After establishing that TGF-beta signalling acts downstream of the HH pathway, we then focus on the identification of genes regulated by TGF-beta during regeneration to investigate how cell recruitment takes place. These data reveal sets of genes that are important in the regeneration and indicate that TGF-beta plays a central role in this process. We propose that members of the *krt* gene family as potential effectors of regenerative TGF-beta signalling and that TGF-beta signalling enables collective cell migration by modulating the expression of different *krts* (Fig. 1i).

TGF-beta has been linked to wound healing by mammalian studies(Lichtman, Otero-Vinas, and Falanga 2016). The initial activation of the pathway during wounding is by serum protease activity which releases TGF-beta1 from platelets. The activated pathway then has many roles including keratinocyte recruitment. Our results suggest that during tail regeneration the role in keratinocyte recruitment is similar, but the initial activation of the TGF-beta pathway involves transcriptional activation of *tgfb1a*. Another similarity is that there is also some evidence for TGF-beta regulation of mammalian *Krts* during wounding. For example, one study has found that *Krt15* is repressed during wound healing and is also repressed by TGF-beta signalling suggesting that it would be in the *krt_off_* synexpression group(Werner and Munz 2000). Furthermore, links between keratin and TGFbeta have been characterised for different tumour types and keratins are used as biomarkers for cancer prognosis(Li et al. 2024; Moll, Divo, and Langbein 2008).

Mammalian studies that primarily use scratch assays have suggested that Krt’s usually have negative roles on cell movement, but can also increase migration (Sanghvi-Shah and Weber 2017; Chung, Rotty, and Coulombe 2013); (Beil et al. 2003; Yoon and Leube 2019). For example, Keratin 6 and 16 are activated after wounding and are thought to play a role in retraction of intermediate filaments from the cell periphery(Paladini et al. 1996) and cause a reduction in cell adhesion(Wawersik and Coulombe 2000); (Wang et al. 2018); (Wawersik and Coulombe 2000). In addition, Krts may affect cell migration by altering cellular elasticity(Beil et al. 2003) and myosin activation(Nanes et al. 2024). In Xenopus keratins are involved with adhesion and cell contraction during gastrulation(Sonavane et al. 2017). Understanding the precise molecular role of zebrafish Krts during collective epithelial cell migration will take further investigation.

Regulated expression of *krts* during regeneration has been observed in adult zebrafish tail regeneration(Martorana et al. 2001), xenopus larval tail regeneration(Tazawa, Shimizu-Nishikawa, and Yoshizato 2006) and newt limbs(Corcoran and Ferretti 1997). This suggests that dynamic expression of *krts* is conserved during organ regeneration. Our finding that *krts* have similar regional expression domains during development and regeneration may indicate that embryonic expression patterns are re-established during regeneration. The significance of regionalised expression domains of *krts* during development has yet to be explored.

**Movie 1 Description.** The movie shows the animal imaged in Figure 9c-f. Panel A shows the maximum intensity projection with yellow arrows following two finfold epithelium nuclei. Panel B shows sections through the somites with a somite boundary position marked with an arrow. Panel C shows superficial sections of the trunk epithelium with arrows following two nuclei.

## Methods

### General methods

Experimental procedures and fish maintenance were performed using standard methods49. All methods were approved by the University of Sheffield and performed in accordance with the relevant guidelines and regulations. This study is reported in accordance with ARRIVE guidelines. All animal husbandry and experimentation was carried out under the supervision and approval of the Home Office (UK) and the University of Sheffield Ethics Board. Adult zebrafish were maintained with a 14 h light/10 h dark cycle at 28 °C according to standard protocols and were mated using pair mating in individual cross tanks. Fish were anesthetised in 40 μg/ml Tricaine (3-amino benzoic acidethylester) in E3. For more information on how individual experiments were performed, please refer to the figure legends and the sections below.

### Bulk RNA-seq analysis

Zebrafish embryos were raised until 72hpf. A scalpel was used to remove the end of the tail from anaesthetised larvae using the pigment gap as a reference (Romero et al). The four conditions were: Unoperated DMSO, Operated DMSO, Unoperated SB431542 and Operated SB431542. There were 3 replicates for each condition and these each contained 100 tails pieces. Each replicate was processed in 1ml of TRI Reagent (Merck) using a plastic pestle to dissociate the tissue. Following the TRI Reagent procedure the RNA was further purified by LiCL precipitation. The RNA quality was checked by Qubit (ThermoFisher) and TapeStation (Agilent) protocols. The libraries were made using the Lexogen QuantSeq 3’ FWD library kit and using NextSeq500 (Illumina) to generate approximately 5 million 75SE reads for each library. The resulting sequence data was demultiplexed using the BlueBee tool (Illumina), aligned to the ensemble zebrafish reference genome and analysed in DESeq2(Love, Huber, and Anders 2014). Plots were generated using Prism 10 (Graphpad). Data sets available at GEO Series GSE296469.

### RNA in situ hybridization

Standard RNA in situ hybridisation was done using the Thisse protocol (Thisse and Thisse). HCR in situ hybridization was performed using zebrafish whole mount protocol(Choi, Schwarzkopf, and Pierce 2020) and reagents purchased from Molecular Instruments (Los Angeles, California). To report in situ expression patterns, a representative animal is shown in the figure panel. For quantification, graphs were generated and analysed in Prism 10 software. Error bars indicate the 95% confidence interval and the centre bar represents the mean. Individual circles represent individual animals tested. The figure legends indicate the type of statistical test applied.

### PhotoConversion

*ubi:h2b-dendra*(Gurskaya et al. 2006) transgenic fish were used for photoconversion experiments. Larvae at 48hpf were placed in a holding chamber in a glass bottom dish. Rectangular ROI was used to draw the region on the tail and photoconverted exposing to 405nm with 100% laser power for 2 min. Unoperated and Operated fish were monitored over a period of time, and images were captured at 543nm and 488nm wavelength using Nikon A1 confocal microscope. For quantification cells at the centre of the trunk were used as a reference point, and the maximum distance travelled by fin fold cells was measured. The analysis was done on six operated animals and three unoperated animals.

### Time-lapse

*actb2:h2b-mCherry* transgenic fish were used for these experiments. Zebrafish Larvae were anaesthetised and imaged by light sheet microscopy (Bruker TrueLive3D) for 280 minutes. For quantification, three fish were analysed, (two started at 22hpf and one at 24hpf). Three epithelial cell nuclei were measured in the proximal fin fold and three epithelial cell nuclei in the centre of the trunk. The boundary between two somites was used as a reference point to standardise movement of the epithelial cells.

### scRNA-seq analysis

Data sets GSE145497 and GSE223922 were downloaded from the NCBI/Geo server. Analysis was done using Seurat version 5.1.0, R (version 4.4.1 2024-06-14) and R Studio (version 2024.04.2+764). A complete list of installed packages and versions is available in Supplemental Fig. 8.

## Acknowledgements

We would like to thank: Gideon Hughes for technical assistance during collection of the tissue for RNA-seq analysis and for providing the *ubi:h2b-dendra* line; Dimitris Rovanias for initial experiments with TGF-beta expression. Fish lines were maintained by the University of Sheffield Biological Services Aquarium under Home Office project licence 3627554. Imaging services were provided by the Wolfson Light Microscopy Facility at the University of Sheffield (Biotechnology and Biological Sciences Research Council Grant BB/W019450/1). Funding provided to HHR by the Biotechnology and Biological Sciences Research Council (Grant BB/S015086/1) and the Leverhulme Trust (Grant RPG-2021-073). For the purpose of open access, the author has applied a Creative Commons Attribution (CC BY) licence to any Author Accepted Manuscript version arising.

## Author Contributions Statement

Experiments were conceived and designed by T.J.C. and H.H.R. and were performed and analysed by D.D.M., Y.L., M.W. H.H.R. and C.Z. The manuscript was written by T.J.C and H.R. All authors reviewed the manuscript.

## Data availability

Data sets from our RNAseq experiment are available from GEO Series GSE296469.

