## Supplemental Figures for "TGF-beta coordinates changes in Keratin gene expression during complex tissue regeneration"

### Supplemental Data

### Supplemental Figures 1-8

Fold change
>5.0 = red
>3.0 = light red
>1.2 = pale red
<0.85 = pale blue
<0.65 = light blue
<0.5 = blue

Gene counts
>1
>10
>100
>1000

p<0.05 p<0.1

| Gene Symbol | Gene Title | ensemble | operated vs unoperated DMSO | operated vs unoperated SB43 | SB43 vs DMSO operated | SB43 vs DMSO unoperated | DMSO unoperated COUNT | DMSO operated COUNT | SB43 unoperated COUNT | SB43 operated COUNT |
|---|---|---|---|---|---|---|---|---|---|---|
| lama3 | laminin subunit alpha-3 | ENSDARG00000022615 | 7.41 | 1.23 | 0.18 | 1.11 | | | | |
| manf | mesencephalic astrocyte-derived neurotrophic | ENSDARG00000063177 | 3.22 | 1.87 | 0.59 | 1.02 | | | | |
| tram1 | translocation associated membrane protein 1 | ENSDARG00000019137 | 2.99 | 1.36 | 0.49 | 1.08 | | | | |
| efemp2a | EGF containing fibulin-like extracellular matrix | ENSDARG00000094324 | 2.92 | 1.98 | 0.61 | 0.90 | | | | |
| krt96 | keratin 96 | ENSDARG00000095147 | 2.86 | 0.59 | 0.19 | 0.92 | | | | |
| si:dkey | protein coding gene | ENSDARG00000090552 | 2.83 | 1.82 | 0.67 | 1.05 | | | | |
| hpcal4 | hippocalcin like 4 | ENSDARG00000070491 | 2.75 | 0.85 | 0.33 | 1.09 | | | | |
| slc39a7 | solute carrier family 39 (zinc transporter), | ENSDARG00000104451 | 2.46 | 1.53 | 0.59 | 0.95 | | | | |
| si:dkey | protein coding gene | ENSDARG00000096678 | 2.13 | 0.77 | 0.39 | 1.07 | | | | |
| BX511082.1 | protein coding gene | ENSDARG00000113678 | 1.82 | 0.99 | 0.52 | 0.97 | | | | |
| serpinh1a | serpin peptidase inhibitor, clade H (heat shock | ENSDARG00000075954 | 1.79 | 1.00 | 0.51 | 0.91 | | | | |
| arhgdig | Rho GDP dissociation inhibitor (GDI) gamma | ENSDARG00000004034 | 1.75 | 1.17 | 0.72 | 1.07 | | | | |
| cpm | carboxypeptidase M | ENSDARG00000011769 | 1.66 | 1.01 | 0.56 | 0.91 | | | | |
| sox4b | SRY (sex determining region Y)-box 4b | ENSDARG00000098834 | 1.56 | 0.82 | 0.51 | 0.96 | | | | |
| sec23a | Sec23 homolog A, COPII coat complex | ENSDARG00000104230 | 1.56 | 1.09 | 0.65 | 0.94 | | | | |
| virma | si:ch211-79l20.4 | ENSDARG00000075824 | 1.50 | 0.92 | 0.57 | 0.94 | | | | |
| dlx5a | distal-less homeobox 5a | ENSDARG00000042296 | 1.47 | 0.81 | 0.53 | 0.96 | | | | |
| bhlha9 | basic helix-loop-helix family, member a9 | ENSDARG00000103981 | 1.47 | 0.77 | 0.50 | 0.95 | | | | |
| psat1 | phosphoserine aminotransferase 1 | ENSDARG00000016733 | 1.41 | 0.75 | 0.50 | 0.94 | | | | |
| cct5 | chaperonin containing TCP1, subunit 5 | ENSDARG00000045399 | 1.40 | 0.97 | 0.63 | 0.91 | | | | |
| acot7 | | ENSDARG00000018653 | 1.39 | 0.77 | 0.51 | 0.92 | | | | |
| psmd6 | proteasome 26S subunit, non-ATPase 6 | ENSDARG00000070674 | 1.37 | 0.94 | 0.62 | 0.90 | | | | |
| fstl1b | follistatin-like 1b | ENSDARG00000039576 | 1.28 | 0.87 | 0.66 | 0.97 | | | | |
| ppp1r3b | protein phosphatase 1, regulatory subunit 3B | ENSDARG00000044691 | 1.27 | 0.57 | 0.44 | 0.98 | | | | |
| gps1 | G protein pathway suppressor 1 | ENSDARG00000040650 | 1.26 | 0.98 | 0.73 | 0.94 | | | | |
| ccdc6a | coiled-coil domain containing 6a | ENSDARG00000043334 | 1.21 | 0.87 | 0.70 | 0.97 | | | | |
| zgc:92744 | zgc:92744 | ENSDARG00000076223 | 1.06 | 0.76 | 0.69 | 0.96 | | | | |
| immt | inner membrane protein, mitochondrial | ENSDARG00000102874 | 1.03 | 0.79 | 0.69 | 0.90 | | | | |
| ACADSB | | ENSDARG00000098174 | 1.03 | 0.71 | 0.61 | 0.88 | | | | |
| btr12 | bloodthirsty-related gene family, member 12 | ENSDARG00000051809 | 0.97 | 3.43 | 3.88 | 1.10 | | | | |
| mfn2 | mitofusin 2 | ENSDARG00000079504 | 0.94 | 0.53 | 0.63 | 1.11 | | | | |
| tmem131 | transmembrane protein 131 | ENSDARG00000056259 | 0.93 | 1.40 | 1.53 | 1.02 | | | | |
| got1 | glutamic-oxaloacetic transaminase 1, soluble | ENSDARG00000039093 | 0.85 | 1.46 | 1.58 | 0.93 | | | | |
| galnt6 | UDP-N-acetyl-alpha-D-galactosamine: | ENSDARG00000014386 | 0.73 | 2.05 | 2.51 | 0.90 | | | | |
| rfx2 | regulatory factor X, 2 (influences HLA class II | ENSDARG00000013575 | 0.72 | 1.33 | 1.91 | 1.03 | | | | |
| pmp22a | peripheral myelin protein 22a | ENSDARG00000105223 | 0.58 | 0.88 | 1.57 | 1.03 | | | | |
| matn3b | | ENSDARG00000069265 | 0.57 | 0.39 | 0.66 | 0.97 | | | | |
| evpla | envoplakin a | ENSDARG00000019808 | 0.53 | 1.04 | 1.91 | 0.97 | | | | |
| fh | fumarate hydratase | ENSDARG00000075132 | 0.52 | 0.84 | 1.63 | 1.01 | | | | |
| lipg | lipase, endothelial | ENSDARG00000031044 | 0.42 | 0.74 | 1.89 | 1.07 | | | | |

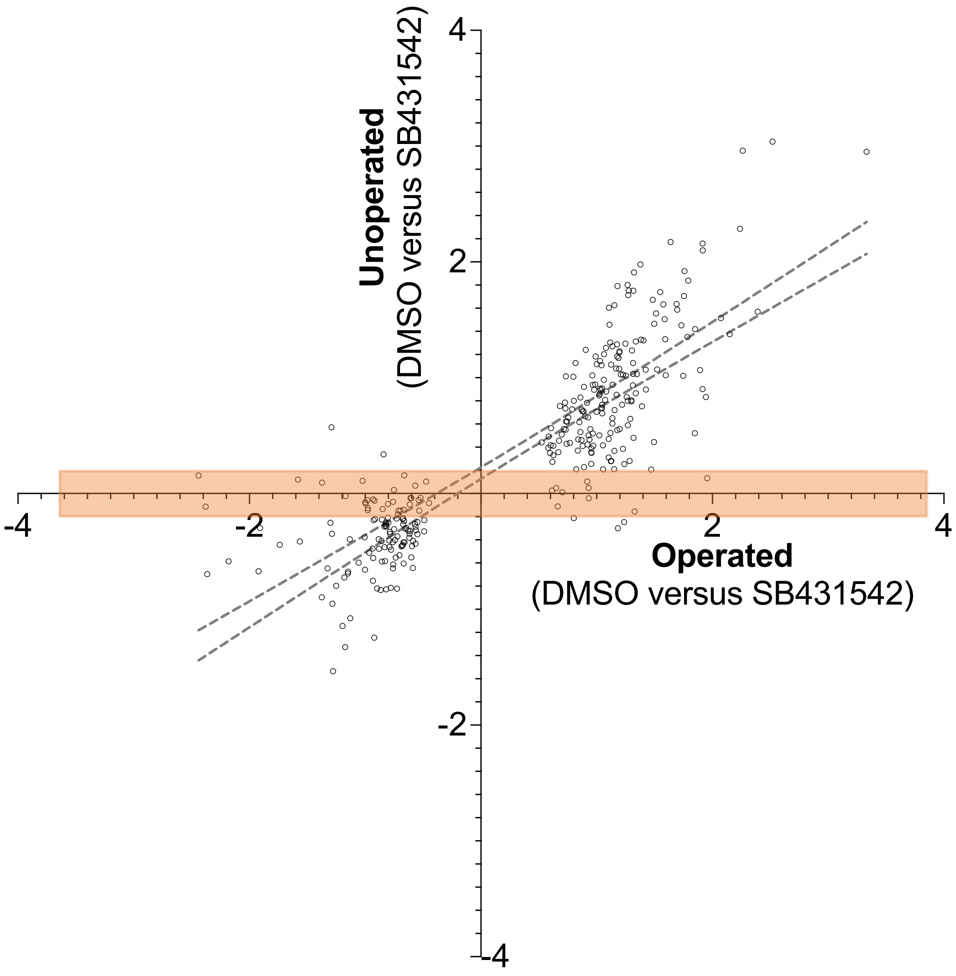


**Supplemental Figure 1.** Potential injury-specific TGF-beta target DEGs. DEGs with SB431542 operated versus DMSO operated (with $P_{adj}$ value < 0.05) and a lack of differential expression in SB431542 unoperated versus DMSO unoperated are presented in this table. The columns show the ensemble identifiers, the fold level of change and the adjusted counts for each sample. These are the DEGs that lie in the boxed area of the graph (as in Fig 3c) and have a $Log_2$ score of between 0.2 and -0.2 in SB431542 unoperated versus DMSO unoperated comparison (i.e. genes that are between 0.87 and 1.15 fold change). Note that the graph is in $Log_2$ scale and the table shows fold change.

Fold change
>5.0 = red
>3.0 = light red
>1.2 = pale red
<0.85 = pale blue
<0.65 = light blue
<0.5 = blue

Gene counts
>1
>10
>100
>1000

p<0.05 p<0.1

| Gene Symbol | Gene Title | ensemble | operated vs unoperated DMSO | operated vs unoperated SB43 | SB43 vs DMSO operated | SB43 vs DMSO unoperated | DMSO unoperated COUNT | DMSO operated COUNT | SB43 unoperated COUNT | SB43 operated COUNT |
|---|---|---|---|---|---|---|---|---|---|---|
| mmp9 | matrix metallopeptidase 9 | ENSDARG00000042816 | 10.35 | 7.44 | 2.42 | 3.37 | | | | |
| lama3 | laminin subunit alpha-3 | ENSDARG00000022615 | 7.41 | 1.23 | 0.18 | 1.11 | | | | |
| ctss2.1 | cathepsin Sb, tandem duplicate 1 | ENSDARG00000074656 | 6.91 | 6.61 | 2.06 | 2.15 | | | | |
| f13a1b | coagulation factor XIII, A1 polypeptide b | ENSDARG00000036893 | 6.28 | 1.73 | 0.41 | 1.49 | | | | |
| sp6 | *Sp6 transcription factor* | ENSDARG00000099880 | 5.89 | 1.95 | 0.22 | 0.67 | | | | |
| si:ch211 | Protein coding | ENSDARG00000060927 | 5.61 | 2.53 | 0.34 | 0.75 | | | | |
| lcp1 | *lymphocyte cytosolic protein 1 (L-plastin)* | ENSDARG00000023188 | 4.46 | 3.25 | 0.48 | 0.66 | | | | |
| f5 | coagulation factor V | ENSDARG00000055705 | 4.40 | 2.30 | 0.41 | 0.79 | | | | |
| apoeb | apolipoprotein Eb | ENSDARG00000040295 | 4.15 | 1.79 | 0.56 | 1.26 | | | | |
| c1qtnf5 | C1q and tumor necrosis factor related protein 5 | ENSDARG00000056134 | 3.88 | 2.42 | 0.40 | 0.64 | | | | |
| snorc | secondary ossification center associated | ENSDARG00000092383 | 3.71 | 1.17 | 0.19 | 0.62 | | | | |
| txn | thioredoxin | ENSDARG00000044125 | 3.36 | 4.22 | 1.81 | 1.44 | | | | |
| manf | mesencephalic astrocyte-derived neurotrophic | ENSDARG00000063177 | 3.22 | 1.87 | 0.59 | 1.02 | | | | |
| loxl5b | *lysyl oxidase-like 5b* | ENSDARG00000076904 | 3.16 | 1.33 | 0.26 | 0.63 | | | | |
| tram1 | translocation associated membrane protein 1 | ENSDARG00000019137 | 2.99 | 1.36 | 0.49 | 1.08 | | | | |
| efemp2a | EGF containing fibulin-like extracellular matrix | ENSDARG00000094324 | 2.92 | 1.98 | 0.61 | 0.90 | | | | |
| cmpk | cytidylate kinase | ENSDARG00000019924 | 2.90 | 2.11 | 0.52 | 0.72 | | | | |
| si:dkey | protein coding gene | ENSDARG00000090552 | 2.83 | 1.82 | 0.67 | 1.05 | | | | |
| hpcal4 | hippocalcin like 4 | ENSDARG00000070491 | 2.75 | 0.85 | 0.33 | 1.09 | | | | |
| krt97 | keratin 97 | ENSDARG00000000212 | 2.66 | 1.08 | 0.30 | 0.74 | | | | |
| slc39a7 | solute carrier family 39 (zinc transporter), | ENSDARG00000104451 | 2.46 | 1.53 | 0.59 | 0.95 | | | | |
| mcm10 | minichromosome maintenance 10 replication | ENSDARG00000045815 | 2.38 | 1.71 | 0.39 | 0.54 | | | | |
| s100a10b | S100 calcium binding protein A10b | ENSDARG00000025254 | 2.08 | 2.58 | 1.59 | 1.28 | | | | |
| prps1b | phosphoribosyl pyrophosphate synthetase 1B | ENSDARG00000037506 | 1.97 | 1.35 | 0.56 | 0.82 | | | | |
| rpa3 | replication protein A3 | ENSDARG00000002613 | 1.86 | 1.47 | 0.60 | 0.76 | | | | |
| ywhaqb | tyrosine 3-monooxygenase/tryptophan 5- | ENSDARG00000023323 | 1.85 | 2.27 | 1.54 | 1.26 | | | | |
| BX511082.1 | protein coding gene | ENSDARG00000113678 | 1.82 | 0.99 | 0.52 | 0.97 | | | | |
| arhgdig | Rho GDP dissociation inhibitor (GDI) gamma | ENSDARG00000004034 | 1.75 | 1.17 | 0.72 | 1.07 | | | | |
| ap1m1 | | ENSDARG00000096249 | 1.67 | 1.21 | 0.63 | 0.86 | | | | |
| cpm | carboxypeptidase M | ENSDARG00000011769 | 1.66 | 1.01 | 0.56 | 0.91 | | | | |
| ccn1l2 | | ENSDARG00000099985 | 1.65 | 1.15 | 0.60 | 0.86 | | | | |
| clta | clathrin, light chain A | ENSDARG00000045618 | 1.64 | 1.27 | 0.56 | 0.73 | | | | |
| arpp19b | cAMP-regulated phosphoprotein 19b /// cAMP- | ENSDARG00000039880 | 1.59 | 1.16 | 0.58 | 0.80 | | | | |
| pqbp1 | polyglutamine binding protein 1 | ENSDARG00000029724 | 1.58 | 1.20 | 0.54 | 0.72 | | | | |
| nars1 | asparaginyl-tRNA synthetase | ENSDARG00000061100 | 1.57 | 1.06 | 0.57 | 0.84 | | | | |
| sec23a | Sec23 homolog A, COPII coat complex | ENSDARG00000104230 | 1.56 | 1.09 | 0.65 | 0.94 | | | | |
| dnajc9 | DnaJ (Hsp40) homolog, subfamily C, member | ENSDARG00000031293 | 1.56 | 1.33 | 0.62 | 0.73 | | | | |
| cd81a | CD81 molecule a | ENSDARG00000036080 | 1.40 | 1.17 | 0.71 | 0.85 | | | | |
| abcb5 | ATP-binding cassette, sub-family B | ENSDARG00000021787 | 0.61 | 0.84 | 1.84 | 1.33 | | | | |
| pmp22a | peripheral myelin protein 22a | ENSDARG00000105223 | 0.58 | 0.88 | 1.57 | 1.03 | | | | |
| sdhaf4 | succinate dehydrogenase complex assembly | ENSDARG00000039390 | 0.58 | 0.94 | 1.94 | 1.19 | | | | |
| PPM1K | | ENSDARG00000076011 | 0.58 | 0.66 | 1.52 | 1.33 | | | | |
| matn3b | | ENSDARG00000069265 | 0.57 | 0.39 | 0.66 | 0.97 | | | | |
| tmem41ab | transmembrane protein 41ab | ENSDARG00000026771 | 0.53 | 0.67 | 1.70 | 1.35 | | | | |
| evpla | envoplakin a | ENSDARG00000019808 | 0.53 | 1.04 | 1.91 | 0.97 | | | | |
| fh | fumarate hydratase | ENSDARG00000075132 | 0.52 | 0.84 | 1.63 | 1.01 | | | | |
| ucp1 | uncoupling protein 1 | ENSDARG00000023151 | 0.49 | 0.77 | 1.85 | 1.17 | | | | |
| si:dkey | protein coding gene | ENSDARG00000101128 | 0.48 | 0.66 | 2.33 | 1.70 | | | | |
| nmrk2 | nicotinamide riboside kinase 2 | ENSDARG00000067848 | 0.45 | 0.61 | 1.96 | 1.43 | | | | |
| lipg | lipase, endothelial | ENSDARG00000031044 | 0.42 | 0.74 | 1.89 | 1.07 | | | | |
| ankrd9 | ankyrin repeat domain 9 | ENSDARG00000028804 | 0.36 | 0.53 | 4.21 | 2.86 | | | | |
| col7a1 | collagen, type VII, alpha 1 | ENSDARG00000021720 | 0.21 | 0.60 | 2.36 | 0.84 | | | | |

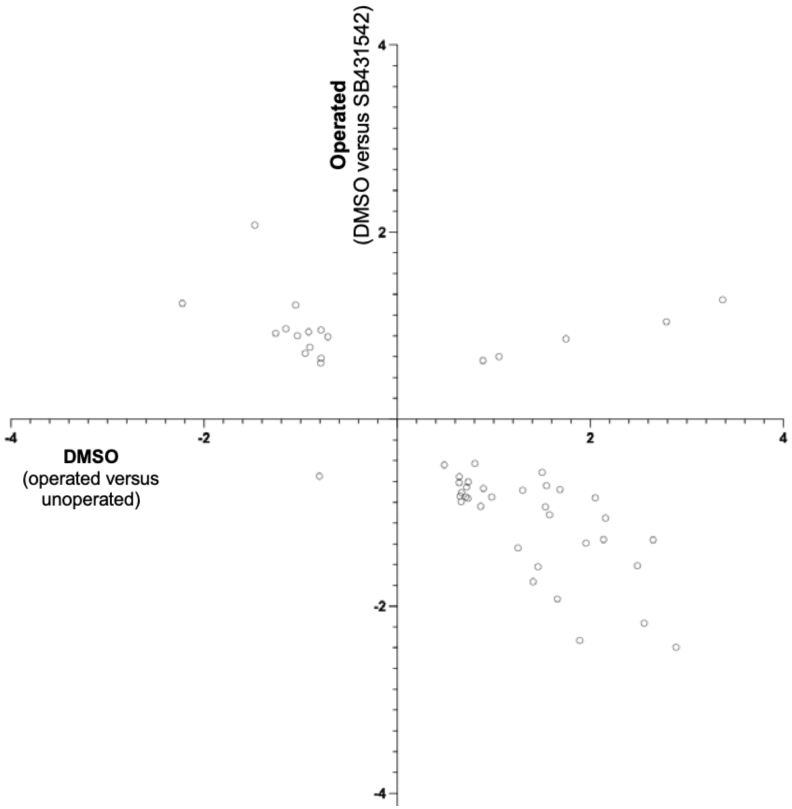


**Supplemental Figure 2.** Potential TGF-beta dependent DEGs. These are DEGs that have a $P_{adj}$ value < 0.05 for both DMSO operated versus DMSO unoperated and SB431542 operated versus DMSO operated. The columns show the ensemble identifiers, the fold level of change and the adjusted counts for each sample. The graph is the same as in Fig. 3d but only showings DEGs that are significant for both comparisons. Note that the graph is in $Log_2$ scale and the table shows fold change.

Fold change
>5.0 = red
>3.0 = light red
>1.2 = pale red
<0.85 = pale blue
<0.65 = light blue
<0.5 = blue

Gene counts
>1
>10
>100
>1000

p<0.05 p<0.1

| Gene Symbol | Gene Title | ensemble | operated vs unoperated DMSO | operated vs unoperated SB43 | SB43 vs DMSO operated | SB43 vs DMSO unoperated | DMSO unoperated COUNT | DMSO operated COUNT | SB43 unoperated COUNT | SB43 operated COUNT |
|---|---|---|---|---|---|---|---|---|---|---|
| fn1b | fibronectin 1b | ENSDARG00000006526 | 32.47 | 21.34 | 0.67 | 1.02 | | | | |
| cart3 | | ENSDARG00000035852 | 18.77 | 12.90 | 0.71 | 1.03 | | | | |
| timp2b | TIMP metallopeptidase inhibitor 2b | ENSDARG00000075261 | 16.15 | 13.34 | 0.74 | 0.90 | | | | |
| grn1 | granulin 1 | ENSDARG00000089362 | 16.13 | 10.23 | 0.80 | 1.26 | | | | |
| si:ch211 | | ENSDARG00000109648 | 9.76 | 5.54 | 0.75 | 1.32 | | | | |
| ms4a17a.4 | membrane-spanning 4-domains | ENSDARG00000014024 | 9.71 | 6.57 | 0.70 | 1.04 | | | | |
| slc7a7 | solute carrier family 7 (amino acid transporter | ENSDARG00000055226 | 7.70 | 5.15 | 0.67 | 1.00 | | | | |
| tgm2l | transglutaminase 2, like | ENSDARG00000093381 | 7.27 | 2.70 | 0.78 | 2.09 | | | | |
| adam8a | | ENSDARG00000001452 | 6.52 | 4.10 | 0.83 | 1.32 | | | | |
| clu | clusterin | ENSDARG00000010434 | 6.43 | 5.29 | 0.75 | 0.91 | | | | |
| ms4a17a.5 | membrane-spanning 4-domains | ENSDARG00000092204 | 6.24 | 3.21 | 0.67 | 1.30 | | | | |
| cbx7a | | ENSDARG00000038025 | 6.16 | 3.66 | 0.69 | 1.16 | | | | |
| c7a | | ENSDARG00000042172 | 5.95 | 4.73 | 0.60 | 0.75 | | | | |
| ms4a17a.7 | membrane-spanning 4-domains | ENSDARG00000043796 | 5.36 | 2.64 | 0.81 | 1.65 | | | | |
| krt18a.1 | keratin 18 | ENSDARG00000018404 | 5.05 | 3.20 | 0.75 | 1.18 | | | | |
| FO704661.1 | | ENSDARG00000102758 | 4.98 | 3.40 | 0.81 | 1.18 | | | | |
| ctsk | cathepsin K | ENSDARG00000040251 | 4.58 | 2.58 | 0.82 | 1.46 | | | | |
| lcp1 | *lymphocyte cytosolic protein 1 (L-plastin)* | ENSDARG00000023188 | 4.46 | 3.25 | 0.48 | 0.66 | | | | |
| mpeg1.1 | | ENSDARG00000055290 | 4.46 | 2.80 | 0.70 | 1.11 | | | | |
| kif23 | kinesin family member 23 | ENSDARG00000014943 | 4.37 | 2.50 | 0.71 | 1.24 | | | | |
| snap23.2 | | ENSDARG00000055252 | 4.35 | 3.77 | 0.70 | 0.81 | | | | |
| apoeb | apolipoprotein Eb | ENSDARG00000040295 | 4.15 | 1.79 | 0.56 | 1.26 | | | | |
| si:rp71 | dispanin subfamily A member 2b-like | ENSDARG00000097746 | 4.09 | 2.90 | 0.75 | 1.06 | | | | |
| ccdc88b | | ENSDARG00000076189 | 3.98 | 3.15 | 0.86 | 1.08 | | | | |
| tagln | transgelin | ENSDARG00000045408 | 3.96 | 2.87 | 0.66 | 0.92 | | | | |
| tnnt2c | troponin T2c, cardiac | ENSDARG00000032242 | 3.90 | 2.82 | 0.73 | 1.01 | | | | |
| c1qtnf5 | C1q and tumor necrosis factor related protein 5 | ENSDARG00000056134 | 3.88 | 2.42 | 0.40 | 0.64 | | | | |
| fcer1gl | | ENSDARG00000104077 | 3.68 | 2.33 | 0.68 | 1.07 | | | | |
| spi1b | Spi-1 proto-oncogene b | ENSDARG00000000767 | 3.65 | 2.49 | 0.72 | 1.06 | | | | |
| manf | mesencephalic astrocyte-derived neurotrophic | ENSDARG00000063177 | 3.22 | 1.87 | 0.59 | 1.02 | | | | |
| si:ch211 | si:ch211-1a19.3 | ENSDARG00000100968 | 3.10 | 2.39 | 0.76 | 0.99 | | | | |
| cdca8 | cell division cycle associated 8 | ENSDARG00000043137 | 3.00 | 2.18 | 0.73 | 1.01 | | | | |
| efemp2a | EGF containing fibulin-like extracellular matrix | ENSDARG00000094324 | 2.92 | 1.98 | 0.61 | 0.90 | | | | |
| cmpk | cytidylate kinase | ENSDARG00000019924 | 2.90 | 2.11 | 0.52 | 0.72 | | | | |
| si:dkey | protein coding gene | ENSDARG00000090552 | 2.83 | 1.82 | 0.67 | 1.05 | | | | |
| hyou1 | hypoxia up-regulated 1 | ENSDARG00000013670 | 2.78 | 1.84 | 0.83 | 1.26 | | | | |
| ms4a17a.12 | membrane-spanning 4-domains | ENSDARG00000053563 | 2.74 | 2.27 | 0.86 | 1.04 | | | | |
| clic2 | | ENSDARG00000010625 | 2.62 | 1.92 | 0.82 | 1.12 | | | | |
| pitpnaa | phosphatidylinositol transfer protein, alpha a | ENSDARG00000039490 | 2.61 | 2.26 | 0.76 | 0.88 | | | | |
| il6st | interleukin 6 signal transducer /// | ENSDARG00000104693 | 2.47 | 2.04 | 0.77 | 0.93 | | | | |
| capgb | capping protein (actin filament), gelsolin-like b | ENSDARG00000099672 | 2.41 | 1.80 | 0.83 | 1.12 | | | | |
| ptges3b | prostaglandin E synthase 3b (cytosolic) | ENSDARG00000089626 | 2.21 | 1.78 | 0.85 | 1.06 | | | | |
| rrbp1a | ribosome binding protein 1 homolog a (dog) | ENSDARG00000013763 | 2.19 | 1.72 | 0.73 | 0.94 | | | | |
| dad1 | defender against cell death 1 | ENSDARG00000102105 | 2.01 | 1.54 | 0.69 | 0.90 | | | | |
| thop1 | zgc:92139 | ENSDARG00000013776 | 1.96 | 1.66 | 0.79 | 0.93 | | | | |
| tmsb4x | thymosin, beta 4 x | ENSDARG00000077777 | 1.72 | 1.47 | 0.83 | 0.96 | | | | |
| atp2a1l | ATPase, Ca++ transporting, cardiac muscle, | ENSDARG00000035458 | 0.52 | 0.69 | 1.36 | 1.03 | | | | |
| cs | citrate synthase | ENSDARG00000103364 | 0.50 | 0.62 | 1.15 | 0.93 | | | | |
| krt4 | keratin 4 | ENSDARG00000017624 | 0.49 | 0.66 | 1.33 | 0.99 | | | | |
| casq1a | | ENSDARG00000038716 | 0.48 | 0.57 | 1.48 | 1.24 | | | | |
| dnajc18 | DnaJ (Hsp40) homolog, subfamily C, member | ENSDARG00000056005 | 0.47 | 0.58 | 1.19 | 0.97 | | | | |
| si:ch211 | | ENSDARG00000096616 | 0.46 | 0.62 | 1.33 | 0.98 | | | | |
| tuba8l2 | tubulin, alpha 8 like 2 | ENSDARG00000031164 | 0.45 | 0.56 | 1.17 | 0.94 | | | | |
| fndc1 | | ENSDARG00000002847 | 0.40 | 0.50 | 1.28 | 1.03 | | | | |
| nrap | | ENSDARG00000009341 | 0.35 | 0.41 | 1.27 | 1.08 | | | | |
| col2a1a | collagen, type II, alpha 1a | ENSDARG00000069093 | 0.32 | 0.37 | 1.77 | 1.51 | | | | |
| col11a2 | collagen, type XI, alpha 2 | ENSDARG00000012422 | 0.30 | 0.38 | 1.16 | 0.92 | | | | |
| epyc | | ENSDARG00000056950 | 0.27 | 0.42 | 1.21 | 0.77 | | | | |

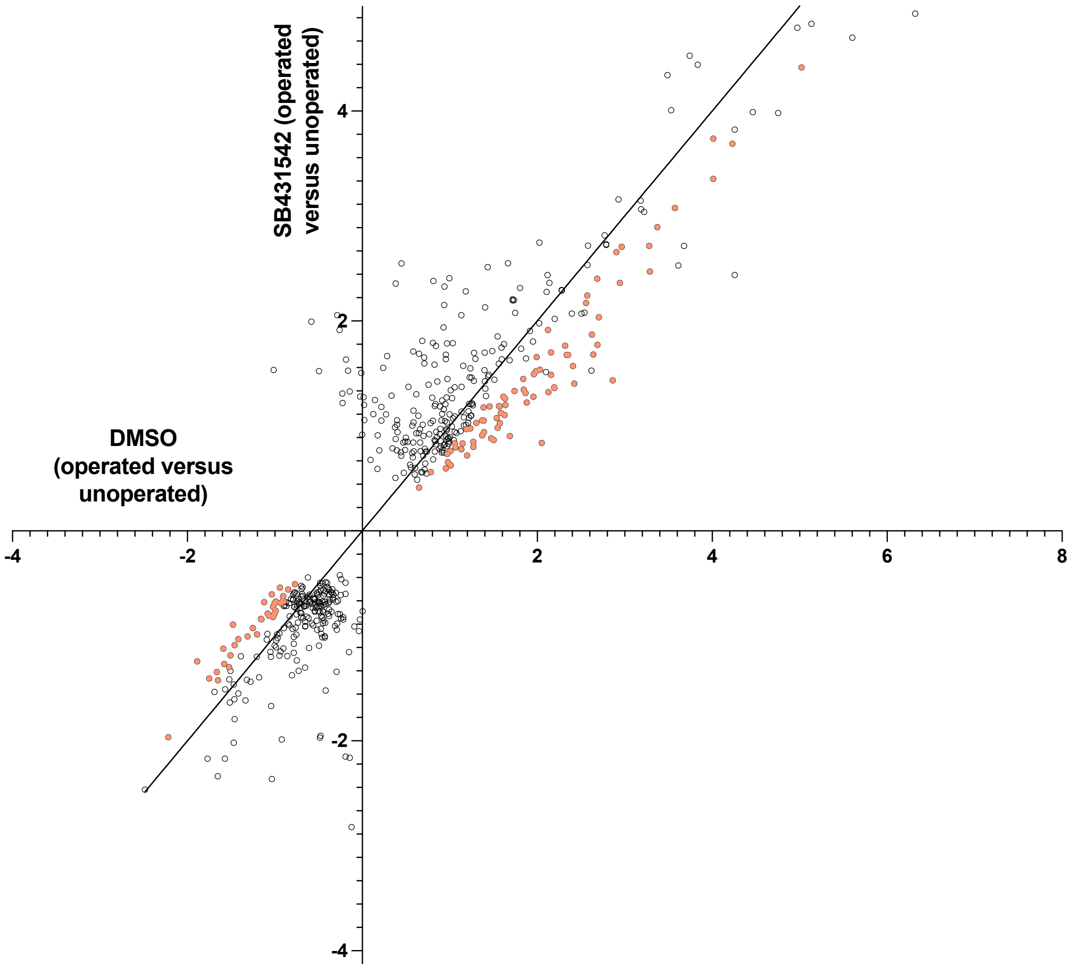


**Supplemental Figure 3. Potential regeneration DEGs.** To generate this table DEGs must meet all of the following criteria (1) a gene count of >10 for all conditions (2) a $P_{adj}$ value < 0.05 for DMSO operated versus unoperated and for SB431542 operated versus unoperated. (3) Either a $Log_2$ score above 0 for SB431542 operated versus unoperated and a ratio below 0.87 for SB431542 operated versus unoperated/ DMSO operated versus unoperated (coloured DEGs in the top right quadrant of graph) OR a $Log_2$ score below 0 for SB431542 operated versus unoperated and a ratio above 1.15 for SB431542 operated versus unoperated/ DMSO operated versus unoperated (coloured DEGs in the lower left quadrant of graph). The columns show the ensemble identifiers, the fold level of change and the adjusted counts for each sample. Note that the graph is the same data as in Fig 3e with the line indicating a slope of one. The graph is in $Log_2$ scale and the table shows fold change.

Fold change
>5.0 = red
>3.0 = light red
>1.2 = pale red
<0.85 = pale blue
<0.65 = light blue
<0.5 = blue
Gene counts
>1
>10
>100
>1000
p<0.05 p<0.1

| Gene Symbol | Gene Title | ensemble | operated vs unoperated DMSO | operated vs unoperated SB43 | SB43 vs DMSO operated | SB43 vs DMSO unoperated | DMSO unoperated COUNT | DMSO operated COUNT | SB43 unoperated COUNT | SB43 operated COUNT |
|---|---|---|---|---|---|---|---|---|---|---|
| si:ch211 | protein coding gene | ENSDARG00000074322 | 9.32 | 8.22 | 1.14 | 1.29 | | | | |
| rrad | Ras-related associated with diabetes | ENSDARG00000052011 | 5.98 | 6.58 | 1.05 | 0.95 | | | | |
| zgc:172053 | protein coding gene | ENSDARG00000038321 | 4.60 | 4.06 | 1.04 | 1.18 | | | | |
| lmna | lamin A | ENSDARG00000013415 | 3.52 | 3.33 | 1.11 | 1.18 | | | | |
| irg1l | immunoresponsive gene 1, like | ENSDARG00000062788 | 3.21 | 3.09 | 1.12 | 1.17 | | | | |
| fanci | Fanconi anemia, complementation group I | ENSDARG00000026224 | 3.07 | 3.43 | 0.93 | 0.83 | | | | |
| parp4 | | ENSDARG00000069934 | 3.02 | 3.36 | 1.05 | 0.94 | | | | |
| ms4a17a.11 | membrane-spanning 4-domains | ENSDARG00000094809 | 2.71 | 2.77 | 0.88 | 0.86 | | | | |
| lrpap1 | low density lipoprotein receptor-related protein | ENSDARG00000033604 | 2.42 | 2.69 | 0.90 | 0.81 | | | | |
| lect2l | | ENSDARG00000033227 | 2.39 | 2.32 | 0.97 | 1.00 | | | | |
| ube2c | ubiquitin-conjugating enzyme E2C | ENSDARG00000114670 | 2.33 | 2.11 | 0.88 | 0.97 | | | | |
| slc38a8b | putative sodium-coupled neutral amino acid | ENSDARG00000054196 | 2.31 | 2.04 | 1.11 | 1.26 | | | | |
| ctsc | cathepsin C | ENSDARG00000101334 | 2.27 | 2.01 | 0.90 | 1.01 | | | | |
| tcirg1b | T-cell, immune regulator 1, ATPase, H+ | ENSDARG00000105142 | 2.24 | 2.01 | 1.07 | 1.19 | | | | |
| hsp90b1 | heat shock protein 90, beta (grp94), member 1 | ENSDARG00000003570 | 2.00 | 1.86 | 0.95 | 1.02 | | | | |
| adamtsl5 | | ENSDARG00000052118 | 1.99 | 2.16 | 1.03 | 0.95 | | | | |
| thbs1a | thrombospondin-1-like | ENSDARG00000100775 | 1.99 | 1.81 | 1.07 | 1.18 | | | | |
| actn4 | actinin, alpha 4 | ENSDARG00000099786 | 1.95 | 2.05 | 1.02 | 0.97 | | | | |
| tgfb1a | transforming growth factor, beta 1a | ENSDARG00000041502 | 1.88 | 1.83 | 0.91 | 0.97 | | | | |
| si:ch1073 | | ENSDARG00000102589 | 1.88 | 1.70 | 0.98 | 1.06 | | | | |
| rnaseka | ribonuclease, RNase K a | ENSDARG00000069461 | 1.87 | 1.68 | 0.94 | 1.06 | | | | |
| ssrp1a | structure specific recognition protein 1a | ENSDARG00000037397 | 1.86 | 1.96 | 1.03 | 0.97 | | | | |
| prkcsh | protein kinase C substrate 80K-H | ENSDARG00000004470 | 1.85 | 1.90 | 1.12 | 1.09 | | | | |
| mbd2 | | ENSDARG00000075952 | 1.81 | 1.86 | 0.96 | 0.93 | | | | |
| anxa2a | annexin A2a | ENSDARG00000003216 | 1.77 | 1.82 | 1.06 | 1.04 | | | | |
| sat1a.2 | | ENSDARG00000095908 | 1.70 | 1.89 | 0.95 | 0.85 | | | | |
| ctsd | cathepsin D | ENSDARG00000057698 | 1.67 | 1.46 | 1.09 | 1.24 | | | | |
| cirbpa | cold inducible RNA binding protein a | ENSDARG00000103672 | 1.65 | 1.69 | 0.90 | 0.89 | | | | |
| arpc3 | actin related protein 2/3 complex, subunit 3 | ENSDARG00000057882 | 1.64 | 1.50 | 0.95 | 1.04 | | | | |
| eif5a2 | eukaryotic translation initiation factor 5A2 | ENSDARG00000056186 | 1.62 | 1.64 | 0.90 | 0.89 | | | | |
| col6a3 | | ENSDARG00000077139 | 1.58 | 1.79 | 1.02 | 0.90 | | | | |
| mkrn1 | makorin, ring finger protein, 1 | ENSDARG00000041665 | 1.52 | 1.52 | 0.91 | 0.92 | | | | |
| flna | | ENSDARG00000074201 | 1.47 | 1.52 | 0.93 | 0.90 | | | | |
| rab1ab | RAB1A, member RAS oncogene family b | ENSDARG00000029683 | 1.46 | 1.55 | 1.12 | 1.05 | | | | |
| hhatlb | hedgehog acyltransferase-like, b | ENSDARG00000005139 | 0.73 | 0.68 | 0.91 | 0.99 | | | | |
| atp5mea | ATP synthase, H+ transporting, mitochondrial | ENSDARG00000078113 | 0.73 | 0.71 | 1.02 | 1.06 | | | | |
| col1a1a | collagen, type I, alpha 1a | ENSDARG00000012405 | 0.72 | 0.68 | 1.01 | 1.07 | | | | |
| stxbp1a | | ENSDARG00000001994 | 0.72 | 0.71 | 0.94 | 0.94 | | | | |
| bckdha | branched chain keto acid dehydrogenase E1, | ENSDARG00000040555 | 0.71 | 0.63 | 0.92 | 1.05 | | | | |
| si:ch211 | | ENSDARG00000092035 | 0.70 | 0.65 | 1.11 | 1.19 | | | | |
| mt-cyb | | ENSDARG00000063924 | 0.70 | 0.66 | 1.04 | 1.09 | | | | |
| adss1 | adenylosuccinate synthase like 1 | ENSDARG00000099517 | 0.69 | 0.64 | 0.94 | 1.00 | | | | |
| pbx3b | pre-B-cell leukemia homeobox 3b | ENSDARG00000013615 | 0.68 | 0.64 | 0.91 | 0.98 | | | | |
| mt-nd4 | | ENSDARG00000063917 | 0.67 | 0.62 | 0.95 | 1.04 | | | | |
| aldoab | aldolase a, fructose-bisphosphate, b | ENSDARG00000034470 | 0.66 | 0.63 | 0.97 | 1.02 | | | | |
| gys1 | | ENSDARG00000016875 | 0.66 | 0.64 | 0.93 | 0.96 | | | | |
| pvalb4 | parvalbumin 4 | ENSDARG00000024433 | 0.66 | 0.66 | 0.99 | 0.99 | | | | |
| synpo2la | | ENSDARG00000077293 | 0.66 | 0.62 | 0.98 | 1.04 | | | | |
| mt-co2 | | ENSDARG00000063908 | 0.65 | 0.65 | 1.01 | 1.01 | | | | |
| tnni4b.2 | troponin I4b, tandem duplicate 2 | ENSDARG00000038671 | 0.65 | 0.74 | 1.14 | 1.01 | | | | |
| slc38a4 | solute carrier family 38, member 4 | ENSDARG00000018149 | 0.65 | 0.64 | 0.94 | 0.94 | | | | |
| ldb3a | LIM domain binding 3a | ENSDARG00000056322 | 0.64 | 0.62 | 0.81 | 0.95 | | | | |
| pdha1a | pyruvate dehydrogenase (lipoamide) alpha 1a | ENSDARG00000012387 | 0.63 | 0.61 | 0.93 | 0.97 | | | | |
| mybpc3 | myosin binding protein C, cardiac | ENSDARG00000011615 | 0.63 | 0.61 | 0.98 | 1.02 | | | | |
| cish | cytokine inducible SH2-containing protein | ENSDARG00000060316 | 0.62 | 0.56 | 0.88 | 0.98 | | | | |
| enah | enabled homolog (Drosophila) | ENSDARG00000032049 | 0.62 | 0.65 | 0.94 | 0.91 | | | | |
| tnmd | | ENSDARG00000052615 | 0.62 | 0.69 | 1.14 | 1.03 | | | | |
| pabpc4 | poly(A) binding protein, cytoplasmic 4 | ENSDARG00000059259 | 0.62 | 0.66 | 0.98 | 0.93 | | | | |
| camk2b1 | calcium/calmodulin-dependent protein kinase II | ENSDARG00000011065 | 0.62 | 0.58 | 1.01 | 1.08 | | | | |
| mbpa | myelin basic protein a | ENSDARG00000036186 | 0.62 | 0.65 | 0.98 | 0.94 | | | | |
| uqcrb | ubiquinol-cytochrome c reductase binding | ENSDARG00000011146 | 0.62 | 0.70 | 1.04 | 0.92 | | | | |
| tnni1d | troponin I, skeletal, slow d | ENSDARG00000073786 | 0.62 | 0.57 | 0.89 | 0.97 | | | | |
| actn3a | actinin alpha 3a | ENSDARG00000013755 | 0.61 | 0.58 | 1.03 | 1.09 | | | | |
| vimr2 | vimentin-related gene | ENSDARG00000105188 | 0.61 | 0.60 | 0.94 | 0.95 | | | | |
| mylz3 | myosin, light polypeptide 3, skeletal muscle | ENSDARG00000017441 | 0.61 | 0.64 | 0.99 | 0.94 | | | | |
| hsc70 | heat shock cognate 70 | ENSDARG00000012381 | 0.61 | 0.62 | 1.03 | 1.00 | | | | |
| cfl2 | | ENSDARG00000014106 | 0.60 | 0.61 | 1.00 | 0.97 | | | | |
| rbfox1l | RNA binding protein, fox-1 homolog (C. | ENSDARG00000021184 | 0.59 | 0.58 | 1.03 | 1.06 | | | | |
| chmd | | ENSDARG00000019342 | 0.59 | 0.60 | 0.89 | 0.88 | | | | |
| atp5f1b | ATP synthase, H+ transporting, mitochondrial | ENSDARG00000070083 | 0.59 | 0.67 | 0.96 | 0.84 | | | | |
| myom2a | si:dkey-78p20.4 | ENSDARG00000075433 | 0.58 | 0.60 | 0.90 | 0.87 | | | | |
| zgc:110843 | zgc:110843 | ENSDARG00000073845 | 0.58 | 0.61 | 0.97 | 0.93 | | | | |
| atp1b3b | ATPase, Na+/K+ transporting, beta 3b | ENSDARG00000042837 | 0.58 | 0.66 | 0.98 | 0.86 | | | | |
| proza | protein Z, vitamin K-dependent plasma | ENSDARG00000037783 | 0.58 | 0.63 | 1.09 | 0.99 | | | | |
| myhz2 | myosin, heavy polypeptide 2, fast muscle | ENSDARG00000012944 | 0.58 | 0.64 | 0.91 | 0.82 | | | | |
| ndst1a | | ENSDARG00000062397 | 0.56 | 0.57 | 0.94 | 0.93 | | | | |
| tnnt3b | troponin T type 3b (skeletal, fast) | ENSDARG00000068457 | 0.55 | 0.61 | 1.02 | 0.92 | | | | |
| mt-nd2 | | ENSDARG00000063899 | 0.55 | 0.62 | 1.00 | 0.87 | | | | |
| slc25a4 | solute carrier family 25 (mitochondrial carrier; | ENSDARG00000027355 | 0.54 | 0.61 | 0.97 | 0.85 | | | | |
| pvalb3 | parvalbumin 3 | ENSDARG00000022817 | 0.54 | 0.61 | 1.10 | 0.97 | | | | |
| krt1c19e | keratin type 1 c19e | ENSDARG00000090268 | 0.47 | 0.51 | 0.96 | 0.90 | | | | |
| sncga | | ENSDARG00000034423 | 0.37 | 0.34 | 1.07 | 1.17 | | | | |
| sparcl1 | | ENSDARG00000074888 | 0.36 | 0.33 | 1.03 | 1.14 | | | | |
| chad | | ENSDARG00000045071 | 0.36 | 0.36 | 0.98 | 0.97 | | | | |
| col9a1b | collagen, type IX, alpha 1b | ENSDARG00000031483 | 0.35 | 0.40 | 1.13 | 1.00 | | | | |
| palmda | palmdelphin a | ENSDARG00000001913 | 0.35 | 0.32 | 1.01 | 1.10 | | | | |
| si:dkey | protein coding gene | ENSDARG00000057504 | 0.18 | 0.18 | 1.03 | 1.01 | | | | |

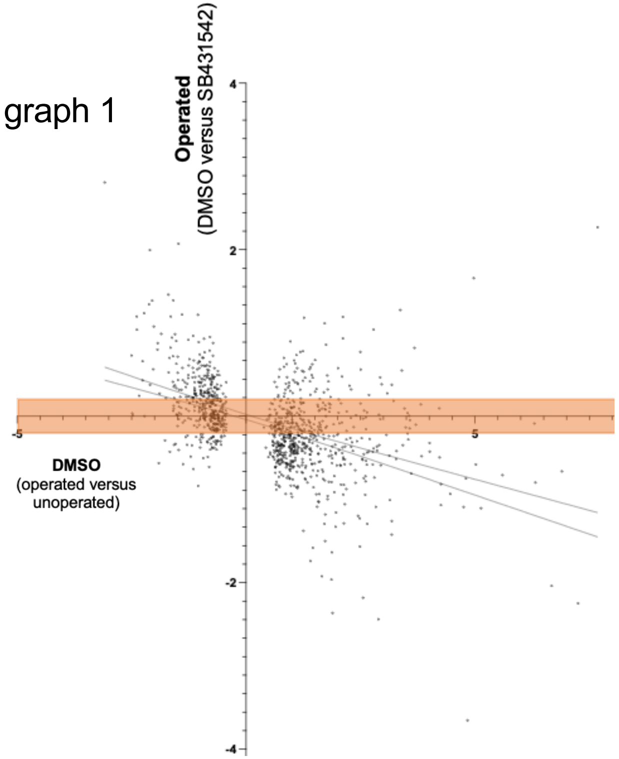


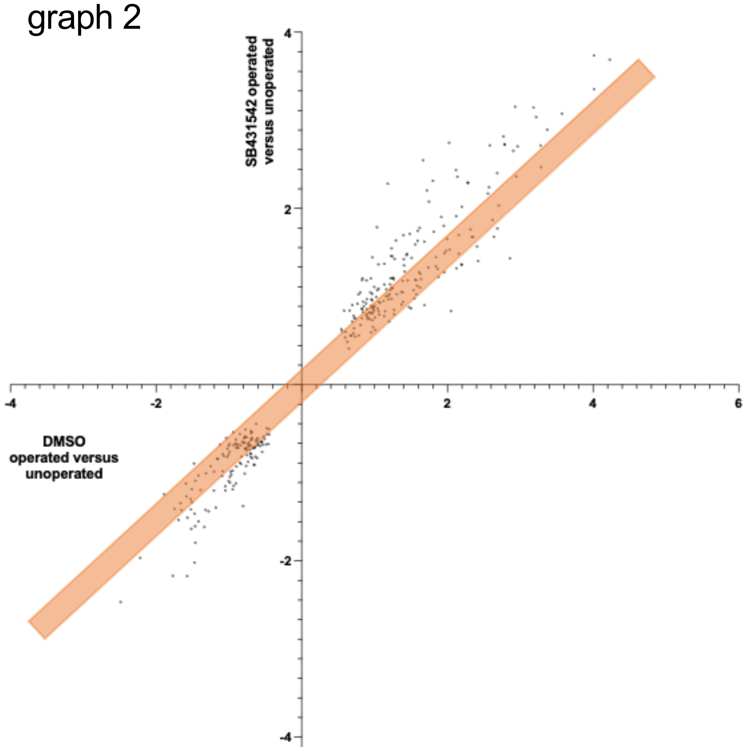


**Supplemental Figure 4. Potential regeneration DEGs that are not strongly regulated by TGF-beta.** To generate this table DEGs must meet all of the following criteria (1) a gene count of >10 for all conditions (2) a $P_{adj}$ value < 0.05 for DMSO operated versus DMSO unoperated and a $Log_2$ score of between 0.2 and -0.2 in the SB431542 operated versus DMSO operated comparison (box in graph 1) (3) a $P_{adj}$ value < 0.05 for SB431542 operated versus unoperated (4) a similar expression change between DMSO operated versus DMSO unoperated and SB431542 operated versus SB431542 unoperated (ratio between 0.87 and 1.15) (box in graph 2). The columns show the ensemble identifiers, the fold level of change and the adjusted counts for each sample. Note that the graph is in $Log_2$ scale and the table shows fold change. Graph 1 is data from Figure 3d and graph 2 is Figure 3e.

**Supplemental Figure 5.** Comparison of families of genes within RNA-seq data. This table shows families of genes with five or more members that were detected in our RNA-seq experiments. The columns show the ensemble identifiers, the fold level of change and the adjusted counts for each sample.

Fold change
>5.0 = red
>3.0 = light red
>1.2 = pale red
<0.85 = pale blue
<0.65 = light blue
<0.5 = blue

Gene counts
>1
>10
>100
>1000

p<0.05 p<0.1

| Gene Symbol | Gene Title | ensemble | (1) DMSO (operated vs unoperated) | (2) Operated (SB431542 vs DMSO) | (3) Unoperated (SB431542 vs DMSO) | (4) SB431542 (operated vs unoperated) |
|---|---|---|---|---|---|---|
| abcb4 | ATP-binding cassette, sub-family B | ENSDARG00000010936 | 7.14 | 0.47 | 1.15 | 2.92 |
| abcc10 | ATP-binding cassette, sub-family C | ENSDARG00000077988 | 2.16 | 1.90 | 2.18 | 1.88 |
| abcf2a | ATP-binding cassette, sub-family F (GCN20), | ENSDARG00000038785 | 1.73 | 0.93 | 1.10 | 1.47 |
| abca1a | ATP-binding cassette, sub-family A (ABC1), | ENSDARG00000074635 | 1.52 | 0.85 | 1.15 | 1.12 |
| abce1 | ATP-binding cassette, sub-family E (OABP), | ENSDARG00000007216 | 1.48 | 0.69 | 0.85 | 1.20 |
| abca1b | ATP-binding cassette, sub-family A (ABC1), | ENSDARG00000079009 | 1.27 | 0.90 | 0.66 | 1.73 |
| abcf1 | ATP-binding cassette, sub-family F (GCN20), | ENSDARG00000031795 | 1.20 | 0.82 | 0.73 | 1.34 |
| abca5 | ATP-binding cassette, sub-family A (ABC1), | ENSDARG00000074041 | 1.03 | 1.03 | 0.78 | 1.35 |
| abca3b | ATP-binding cassette, sub-family A (ABC1), | ENSDARG00000100524 | 1.02 | 1.13 | 0.94 | 1.23 |
| abcb7 | ATP-binding cassette, sub-family B | ENSDARG00000062795 | 0.96 | 0.87 | 1.07 | 0.78 |
| abcd3a | ATP-binding cassette, sub-family D (ALD), | ENSDARG00000104085 | 0.93 | 0.74 | 0.83 | 0.83 |
| abcc5 | ATP-binding cassette, sub-family C | ENSDARG00000061233 | 0.88 | 2.13 | 0.60 | 3.10 |
| abcb8 | ATP-binding cassette, sub-family B | ENSDARG00000056672 | 0.64 | 2.31 | 1.88 | 0.78 |
| abcb5 | ATP-binding cassette, sub-family B | ENSDARG00000021787 | 0.61 | 1.84 | 1.33 | 0.84 |
| abcd4 | ATP-binding cassette, sub-family D (ALD), | ENSDARG00000061770 | 0.57 | 4.75 | 2.48 | 1.10 |
| actr2a | ARP2 actin-related protein 2a homolog (yeast) | ENSDARG00000052438 | 2.98 | 1.08 | 1.58 | 2.04 |
| actb1 | actin, beta 1 | ENSDARG00000037746 | 1.72 | 0.99 | 0.83 | 2.04 |
| actb2 | actin, beta 2 | ENSDARG00000037870 | 1.24 | 1.02 | 1.10 | 1.15 |
| actl6a | actin-like 6A | ENSDARG00000070828 | 1.06 | 1.21 | 0.72 | 1.77 |
| actr6 | ARP6 actin-related protein 6 homolog (yeast) | ENSDARG00000021370 | 1.05 | 2.37 | 1.84 | 1.36 |
| actr1 | ARP1 actin-related protein 1, centractin (yeast) | ENSDARG00000011611 | 1.05 | 1.21 | 1.10 | 1.16 |
| actr8 | ARP8 actin-related protein 8 homolog (yeast) | ENSDARG00000103610 | 1.05 | 0.64 | 1.01 | 0.66 |
| actr10 | actin-related protein 10 homolog (S. cerevisiae) | ENSDARG00000038432 | 0.94 | 0.84 | 0.72 | 1.10 |
| actc1b | actin, alpha, cardiac muscle 1b | ENSDARG00000099197 | 0.69 | 0.83 | 0.97 | 0.59 |
| acta2 | actin, alpha 2, smooth muscle, aorta | ENSDARG00000045180 | 0.64 | 0.84 | 0.69 | 0.78 |
| actc1 | actin, alpha, cardiac muscle 2 | ENSDARG00000057911 | 0.64 | 0.79 | 1.23 | 0.41 |
| actc1a | actin alpha cardiac muscle 1a | ENSDARG00000042535 | 0.59 | 0.83 | 0.93 | 0.53 |
| acta1b | actin alpha 1, skeletal muscle b | ENSDARG00000055618 | 0.50 | 0.98 | 0.84 | 0.58 |
| acta1a | actin, alpha 1a, skeletal muscle | ENSDARG00000036371 | 0.35 | 1.01 | 0.81 | 0.44 |
| actc1c | actin alpha cardiac muscle 1c | ENSDARG00000079111 | 0.21 | 1.00 | 0.84 | 0.26 |
| actn1 | actinin, alpha 1 | ENSDARG00000007219 | 2.58 | 3.11 | 4.51 | 1.78 |
| actn4 | actinin, alpha 4 | ENSDARG00000099786 | 1.95 | 1.02 | 0.97 | 2.05 |
| actn3b | actinin alpha 3b | ENSDARG00000001431 | 0.78 | 1.15 | 1.35 | 0.67 |
| actn2b | actinin, alpha 2b | ENSDARG00000071090 | 0.73 | 0.70 | 0.74 | 0.69 |
| actn3a | actinin alpha 3a | ENSDARG00000013755 | 0.61 | 1.03 | 1.09 | 0.58 |
| ankrd10a | ankyrin repeat domain 10a | ENSDARG00000037100 | 1.81 | 0.79 | 0.96 | 1.50 |
| ankrd49 | ankyrin repeat domain 49 | ENSDARG00000023508 | 1.53 | 0.90 | 1.33 | 1.04 |
| ankmy2a | ankyrin repeat and MYND domain containing | ENSDARG00000005948 | 1.48 | 0.50 | 0.96 | 0.77 |
| ankfy1 | ankyrin repeat and FYVE domain containing 1 | ENSDARG00000061013 | 1.22 | 0.78 | 0.89 | 1.07 |
| ankrd28b | ankyrin repeat domain 28b | ENSDARG00000009023 | 1.12 | 1.13 | 1.27 | 1.00 |
| ankar | ankyrin and armadillo repeat containing | ENSDARG00000000516 | 1.00 | 2.31 | 1.00 | 1.70 |
| ankhd1 | ankyrin repeat and KH domain containing 1 | ENSDARG00000077860 | 0.97 | 1.05 | 0.86 | 1.18 |
| ankra2 | ankyrin repeat, family A (RFXANK-like), 2 | ENSDARG00000035399 | 0.95 | 0.97 | 1.33 | 0.69 |
| ankrd24 | ankyrin repeat domain 24 | ENSDARG00000062103 | 0.95 | 0.40 | 0.69 | 0.55 |
| ankmy1 | ankyrin repeat and MYND domain containing 1 | ENSDARG00000062702 | 0.94 | 2.15 | 0.78 | 2.61 |
| ankrd13c | ankyrin repeat domain 13C | ENSDARG00000103831 | 0.93 | 0.89 | 0.91 | 0.91 |
| anks1b | ankyrin repeat and sterile alpha motif domain | ENSDARG00000003512 | 0.79 | 0.79 | 0.90 | 0.69 |
| ankrd22 | ankyrin repeat domain 22 | ENSDARG00000002298 | 0.76 | 1.48 | 1.41 | 0.79 |
| ankrd46b | ankyrin repeat domain 46b | ENSDARG00000015780 | 0.71 | 1.38 | 1.56 | 0.63 |

Fold change
>5.0 = red
>3.0 = light red
>1.2 = pale red
<0.85 = pale blue
<0.65 = light blue
<0.5 = blue

Gene counts
>1
>10
>100
>1000

p<0.05 p<0.1

| Gene Symbol | Gene Title | ensemble | (1) DMSO (operated vs unoperated) | (2) Operated (SB431542 vs DMSO) | (3) Unoperated (SB431542 vs DMSO) | (4) SB431542 (operated vs unoperated) | DMSO unoperated COUNT | DMSO operated COUNT | SB43 unoperated COUNT | SB43 operated COUNT |
|---|---|---|---|---|---|---|---|---|---|---|
| ankrd12 | ankyrin repeat domain 12 | ENSDARG00000052419 | 0.68 | 1.00 | 0.67 | 1.02 | | | | |
| ank2b | ankyrin 2b, neuronal | ENSDARG00000043313 | 0.59 | 1.65 | 0.89 | 1.10 | | | | |
| ankrd1b | ankyrin repeat domain 1b (cardiac muscle) | ENSDARG00000076192 | 0.53 | 1.23 | 1.43 | 0.46 | | | | |
| ankrd11 | ankyrin repeat domain 11 | ENSDARG00000051886 | 0.52 | 1.86 | 0.83 | 1.16 | | | | |
| ank1a | ankyrin 1, erythrocytic a | ENSDARG00000092143 | 0.40 | 1.18 | 0.75 | 0.63 | | | | |
| ankrd9 | ankyrin repeat domain 9 | ENSDARG00000028804 | 0.36 | 4.21 | 2.86 | 0.53 | | | | |
| ankrd1a | ankyrin repeat domain 1a (cardiac muscle) | ENSDARG00000075263 | 0.28 | 1.99 | 1.15 | 0.48 | | | | |
| apoda.2 | apolipoprotein Da, duplicate 2 | ENSDARG00000060350 | 14.99 | 0.38 | 7.41 | 0.77 | | | | |
| apoba | apolipoprotein Ba | ENSDARG00000042780 | 11.88 | 0.07 | 3.53 | 0.24 | | | | |
| apobb.1 | apolipoprotein Bb, tandem duplicate 1 | ENSDARG00000022767 | 8.78 | 0.12 | 3.74 | 0.28 | | | | |
| apoea | apolipoprotein Ea | ENSDARG00000102004 | 7.70 | 0.38 | 4.57 | 0.63 | | | | |
| apoa4b.2 | apolipoprotein A-IV b, tandem duplicate 2 | ENSDARG00000020866 | 7.45 | 0.06 | 2.89 | 0.15 | | | | |
| apoa1a | apolipoprotein A-Ia | ENSDARG00000012076 | 7.29 | 0.75 | 3.44 | 1.59 | | | | |
| apoc1 | apolipoprotein C-I like | ENSDARG00000092170 | 5.77 | 0.83 | 2.29 | 2.10 | | | | |
| apobb.2 | apolipoprotein B-100 | ENSDARG00000075016 | 5.56 | 0.15 | 1.00 | 1.00 | | | | |
| apoa2 | apolipoprotein A-II | ENSDARG00000015866 | 4.40 | 0.43 | 1.55 | 1.23 | | | | |
| apoeb | apolipoprotein Eb | ENSDARG00000040295 | 4.15 | 0.56 | 1.26 | 1.79 | | | | |
| apoa4b.1 | apolipoprotein A-IV b, tandem duplicate 1 | ENSDARG00000040298 | 3.26 | 0.46 | 0.93 | 1.62 | | | | |
| apoa1b | apolipoprotein A-Ib | ENSDARG00000101324 | 1.61 | 1.06 | 1.54 | 1.11 | | | | |
| apodb | apolipoprotein Db | ENSDARG00000057437 | 1.34 | 1.74 | 2.01 | 1.16 | | | | |
| apoa4a | apolipoprotein A-IV a | ENSDARG00000101160 | 1.00 | 6.86 | 3.04 | 1.63 | | | | |
| apoob | apolipoprotein O, b | ENSDARG00000026444 | 0.99 | 0.76 | 0.90 | 0.84 | | | | |
| apol1 | apolipoprotein L, 1 | ENSDARG00000007425 | 0.96 | 1.24 | 1.68 | 0.71 | | | | |
| apool | apolipoprotein O-like | ENSDARG00000039374 | 0.94 | 1.05 | 1.42 | 0.69 | | | | |
| apoc2 | apolipoprotein C-II | ENSDARG00000092155 | 0.84 | 0.83 | 0.33 | 2.07 | | | | |
| apooa | apolipoprotein O, a | ENSDARG00000104696 | 0.74 | 1.15 | 1.11 | 0.76 | | | | |
| apof | apolipoprotein F | ENSDARG00000090980 | 0.68 | 2.83 | 3.01 | 0.64 | | | | |
| arpc5a | actin related protein 2/3 complex, subunit 5A | ENSDARG00000039142 | 2.95 | 0.78 | 0.76 | 3.01 | | | | |
| arpc5b | actin related protein 2/3 complex, subunit 5B | ENSDARG00000019062 | 2.61 | 0.95 | 2.39 | 1.04 | | | | |
| arpc1b | actin related protein 2/3 complex, subunit 1B | ENSDARG00000027063 | 2.24 | 1.01 | 1.31 | 1.73 | | | | |
| arpc3 | actin related protein 2/3 complex, subunit 3 | ENSDARG00000057882 | 1.64 | 0.95 | 1.04 | 1.50 | | | | |
| arpc4 | actin related protein 2/3 complex, subunit 4 | ENSDARG00000054063 | 1.61 | 0.94 | 1.17 | 1.29 | | | | |
| arpc2 | actin related protein 2/3 complex, subunit 2 | ENSDARG00000075989 | 1.61 | 0.78 | 0.93 | 1.35 | | | | |
| arpc4l | actin related protein 2/3 complex, subunit 4, | ENSDARG00000058225 | 1.32 | 1.41 | 1.34 | 1.39 | | | | |
| arpc1a | actin related protein 2/3 complex, subunit 1A | ENSDARG00000008383 | 1.18 | 0.83 | 0.76 | 1.29 | | | | |
| atp9b | ATPase, class II, type 9B | ENSDARG00000062521 | 20.41 | 0.47 | 16.43 | 0.58 | | | | |
| atp2a2b | ATPase, Ca++ transporting, cardiac muscle, | ENSDARG00000005122 | 2.82 | 0.89 | 1.61 | 1.57 | | | | |
| atp6v0b | ATPase H+ transporting V0 subunit b | ENSDARG00000031681 | 2.75 | 1.38 | 2.14 | 1.77 | | | | |
| atp6v1ba | ATPase, H+ transporting, lysosomal, V1 | ENSDARG00000013443 | 2.07 | 0.75 | 1.10 | 1.41 | | | | |
| atp6ap1b | ATPase, H+ transporting, lysosomal accessory | ENSDARG00000037153 | 2.07 | 0.73 | 1.17 | 1.30 | | | | |
| atp6v0ca | ATPase, H+ transporting, lysosomal, V0 | ENSDARG00000057853 | 1.96 | 0.65 | 0.80 | 1.58 | | | | |
| atp6v1ab | ATPase, H+ transporting, lysosomal, V1 | ENSDARG00000076318 | 1.75 | 0.86 | 1.33 | 1.13 | | | | |
| atp6v1f | ATPase, H+ transporting, lysosomal, V1 | ENSDARG00000045543 | 1.61 | 0.63 | 0.97 | 1.05 | | | | |
| atp6v1h | ATPase, H+ transporting, lysosomal V1 subunit | ENSDARG00000006370 | 1.59 | 0.75 | 1.04 | 1.15 | | | | |
| atp6v1aa | ATPase H+ transporting V1 subunit Aa | ENSDARG00000034534 | 1.43 | 0.80 | 0.64 | 1.80 | | | | |
| atp6v1d | ATPase, H+ transporting, lysosomal V1 subunit | ENSDARG00000011175 | 1.36 | 0.69 | 0.84 | 1.11 | | | | |
| atp6v1e1b | ATPase, H+ transporting, lysosomal, V1 | ENSDARG00000030694 | 1.33 | 0.85 | 0.94 | 1.20 | | | | |
| atp1a1a.1 | ATPase, Na+/K+ transporting, alpha 1a | ENSDARG00000002791 | 1.30 | 0.96 | 0.88 | 1.42 | | | | |
| atp6v1g1 | ATPase, H+ transporting, lysosomal, V1 | ENSDARG00000022315 | 1.29 | 0.95 | 1.13 | 1.08 | | | | |
| atp6ap2 | ATPase, H+ transporting, lysosomal accessory | ENSDARG00000008735 | 1.29 | 0.80 | 1.04 | 0.99 | | | | |

Fold change
>5.0 = red
>3.0 = light red
>1.2 = pale red
<0.85 = pale blue
<0.65 = light blue
<0.5 = blue

Gene counts
>1
>10
>100
>1000

p<0.05 p<0.1

| Gene Symbol | Gene Title | ensemble | (1) DMSO (operated vs unoperated) | (2) Operated (SB431542 vs DMSO) | (3) Unoperated (SB431542 vs DMSO) | (4) SB431542 (operated vs unoperated) | DMSO unoperated COUNT | DMSO operated COUNT | SB43 unoperated COUNT | SB43 operated COUNT |
|---|---|---|---|---|---|---|---|---|---|---|
| atp6v1c1b | ATPase, H+ transporting, lysosomal, V1 | ENSDARG00000035880 | 1.28 | 0.79 | 0.74 | 1.36 | | | | |
| atp1b1b | ATPase, Na+/K+ transporting, beta 1b | ENSDARG00000076833 | 1.24 | 1.06 | 1.49 | 0.88 | | | | |
| atp6v0d1 | ATPase, H+ transporting, lysosomal V0 subunit | ENSDARG00000069090 | 1.23 | 1.03 | 1.21 | 1.05 | | | | |
| atp6v1c1a | ATPase, H+ transporting, lysosomal, V1 | ENSDARG00000023967 | 1.13 | 1.40 | 1.46 | 1.09 | | | | |
| atp13a1 | ATPase type 13A1 | ENSDARG00000029931 | 1.13 | 1.00 | 0.60 | 1.88 | | | | |
| atp1b2b | ATPase, Na+/K+ transporting, beta 2b | ENSDARG00000034424 | 1.12 | 0.59 | 0.52 | 1.26 | | | | |
| atp1b1a | ATPase, Na+/K+ transporting, beta 1a | ENSDARG00000013144 | 1.12 | 0.79 | 0.81 | 1.09 | | | | |
| atp1b2a | ATPase, Na+/K+ transporting, beta 2a | ENSDARG00000099203 | 1.08 | 0.96 | 1.36 | 0.77 | | | | |
| atp1a1a.2 | ATPase, Na+/K+ transporting, alpha 1a | ENSDARG00000007739 | 1.03 | 0.88 | 1.15 | 0.79 | | | | |
| atp5if1b | ATPase inhibitory factor 1b | ENSDARG00000044092 | 0.97 | 1.15 | 1.50 | 0.75 | | | | |
| atp1a1b | ATPase, Na+/K+ transporting, alpha 1b | ENSDARG00000019856 | 0.96 | 0.83 | 0.79 | 1.01 | | | | |
| atpv0e2 | ATPase, H+ transporting V0 subunit e2 | ENSDARG00000059057 | 0.91 | 1.00 | 1.64 | 0.56 | | | | |
| atp10b | ATPase, class V, type 10B | ENSDARG00000076230 | 0.86 | 1.12 | 1.10 | 0.88 | | | | |
| atp1b3a | ATPase, Na+/K+ transporting, beta 3a | ENSDARG00000015790 | 0.84 | 1.08 | 1.07 | 0.84 | | | | |
| atp6v1b2 | ATPase, H+ transporting, lysosomal, V1 | ENSDARG00000043465 | 0.82 | 1.17 | 1.00 | 0.96 | | | | |
| atp2b4 | ATPase, Ca++ transporting, plasma membrane | ENSDARG00000044902 | 0.80 | 1.01 | 0.76 | 1.07 | | | | |
| atp2a1 | ATPase, Ca++ transporting, cardiac muscle, | ENSDARG00000020574 | 0.80 | 1.12 | 1.13 | 0.79 | | | | |
| atp2b3b | ATPase plasma membrane Ca2+ transporting | ENSDARG00000023445 | 0.76 | 0.94 | 1.16 | 0.62 | | | | |
| atp2b2 | ATPase, Ca++ transporting, plasma membrane | ENSDARG00000063433 | 0.76 | 0.96 | 0.82 | 0.89 | | | | |
| atp1a3b | ATPase, Na+/K+ transporting, alpha 3b | ENSDARG00000104139 | 0.76 | 1.03 | 0.77 | 1.01 | | | | |
| atp2b3a | ATPase, Ca++ transporting, plasma membrane | ENSDARG00000043474 | 0.76 | 1.06 | 0.75 | 1.08 | | | | |
| atp1a2a | ATPase, Na+/K+ transporting, alpha 2a | ENSDARG00000010472 | 0.74 | 1.72 | 1.63 | 0.79 | | | | |
| atp1a3a | ATPase, Na+/K+ transporting, alpha 3a | ENSDARG00000018259 | 0.74 | 1.13 | 0.93 | 0.89 | | | | |
| atp2b1a | ATPase, Ca++ transporting, plasma membrane | ENSDARG00000012684 | 0.71 | 0.83 | 0.75 | 0.79 | | | | |
| atp5if1a | ATPase inhibitory factor 1a | ENSDARG00000067975 | 0.70 | 0.76 | 0.70 | 0.76 | | | | |
| atp6ap1la | ATPase, H+ transporting, lysosomal accessory | ENSDARG00000091509 | 0.63 | 0.76 | 0.61 | 0.79 | | | | |
| atp1b3b | ATPase, Na+/K+ transporting, beta 3b | ENSDARG00000042837 | 0.58 | 0.98 | 0.86 | 0.66 | | | | |
| atp2a2a | ATPase, Ca++ transporting, cardiac muscle, | ENSDARG00000029439 | 0.56 | 0.98 | 0.73 | 0.75 | | | | |
| atp6v0cb | ATPase, H+ transporting, lysosomal, V0 | ENSDARG00000036577 | 0.54 | 1.07 | 0.80 | 0.73 | | | | |
| atp2a1l | ATPase, Ca++ transporting, cardiac muscle, | ENSDARG00000035458 | 0.52 | 1.36 | 1.03 | 0.69 | | | | |
| atp1a1a.4 | ATPase, Na+/K+ transporting, alpha 1a | ENSDARG00000001870 | 0.52 | 1.70 | 0.85 | 1.04 | | | | |
| chd4a | chromodomain helicase DNA binding protein | ENSDARG00000063535 | 1.32 | 0.98 | 0.76 | 1.70 | | | | |
| chd1 | chromodomain helicase DNA binding protein 1 | ENSDARG00000103787 | 1.24 | 0.93 | 0.94 | 1.23 | | | | |
| chd1l | chromodomain helicase DNA binding protein 1- | ENSDARG00000015471 | 1.18 | 0.85 | 1.00 | 1.01 | | | | |
| chd4b | chromodomain helicase DNA binding protein | ENSDARG00000025789 | 1.05 | 1.05 | 0.94 | 1.17 | | | | |
| chd7 | chromodomain helicase DNA binding protein 7 | ENSDARG00000075211 | 0.88 | 0.98 | 0.78 | 1.11 | | | | |
| chd2 | chromodomain helicase DNA binding protein 2 | ENSDARG00000060687 | 0.81 | 1.25 | 0.91 | 1.11 | | | | |
| chd9 | chromodomain helicase DNA binding protein 9 | ENSDARG00000074498 | 0.77 | 1.03 | 0.86 | 0.92 | | | | |
| chd6 | chromodomain helicase DNA binding protein 6 | ENSDARG00000017244 | 0.75 | 0.87 | 0.65 | 1.00 | | | | |
| chd5 | chromodomain helicase DNA binding protein 5 | ENSDARG00000105083 | 0.57 | 0.74 | 0.45 | 0.93 | | | | |
| chd3 | chromodomain helicase DNA binding protein 3 | ENSDARG00000021405 | 0.53 | 0.98 | 0.72 | 0.72 | | | | |
| cldn5a | claudin 5a | ENSDARG00000043716 | 1.52 | 2.35 | 0.93 | 3.85 | | | | |
| cldni | claudin i | ENSDARG00000054616 | 1.42 | 1.01 | 1.04 | 1.38 | | | | |
| cldne | claudin e | ENSDARG00000043128 | 1.23 | 1.37 | 1.54 | 1.10 | | | | |
| cldn7b | claudin 7b | ENSDARG00000014047 | 1.19 | 1.45 | 1.52 | 1.13 | | | | |
| cldn15la | claudin 15-like a | ENSDARG00000016081 | 1.15 | 1.63 | 2.68 | 0.70 | | | | |
| cldn12 | claudin 12 | ENSDARG00000003927 | 1.10 | 0.77 | 0.88 | 0.96 | | | | |
| cldna | claudin a | ENSDARG00000069888 | 0.91 | 3.36 | 1.21 | 2.51 | | | | |
| cldnb | claudin b | ENSDARG00000009544 | 0.88 | 1.84 | 1.57 | 1.03 | | | | |
| cldnh | claudin h | ENSDARG00000069503 | 0.79 | 1.42 | 1.11 | 1.02 | | | | |
| cldn11b | claudin 11b | ENSDARG00000030723 | 0.78 | 0.91 | 0.92 | 0.76 | | | | |

Fold change
>5.0 = red
>3.0 = light red
>1.2 = pale red
<0.85 = pale blue
<0.65 = light blue
<0.5 = blue

Gene counts
>1
>10
>100
>1000

p<0.05 p<0.1

| Gene Symbol | Gene Title | ensemble | (1) DMSO (operated vs unoperated) | (2) Operated (SB431542 vs DMSO) | (3) Unoperated (SB431542 vs DMSO) | (4) SB431542 (operated vs unoperated) | DMSO unoperated COUNT | DMSO operated COUNT | SB43 unoperated COUNT | SB43 operated COUNT |
|---|---|---|---|---|---|---|---|---|---|---|
| cldng | claudin g | ENSDARG00000003701 | 0.74 | 0.87 | 0.92 | 0.71 | | | | |
| cldnk | claudin k | ENSDARG00000042357 | 0.74 | 1.05 | 1.33 | 0.58 | | | | |
| cldn8.3 | claudin 8.3 | ENSDARG00000099950 | 0.72 | 1.60 | 4.43 | 0.26 | | | | |
| cldnd1a | claudin domain containing 1a | ENSDARG00000104439 | 0.71 | 0.74 | 0.45 | 1.16 | | | | |
| cldnc | claudin c | ENSDARG00000015955 | 0.70 | 2.99 | 1.86 | 1.13 | | | | |
| cldn19 | claudin 19 | ENSDARG00000044569 | 0.61 | 1.87 | 0.86 | 1.34 | | | | |
| cldnd | claudin d | ENSDARG00000006580 | 0.59 | 0.85 | 1.00 | 0.50 | | | | |
| cldn11a | claudin 11a | ENSDARG00000020031 | 0.50 | 0.91 | 0.86 | 0.53 | | | | |
| col17a1b | collagen, type XVII, alpha 1b | ENSDARG00000079011 | 1.46 | 0.87 | 1.04 | 1.23 | | | | |
| col12a1a | collagen, type XII, alpha 1a | ENSDARG00000078322 | 1.45 | 0.81 | 0.94 | 1.26 | | | | |
| col5a3a | collagen type V alpha-3a | ENSDARG00000098294 | 1.33 | 0.58 | 0.20 | 3.92 | | | | |
| col5a2a | collagen, type V, alpha 2a | ENSDARG00000031678 | 1.13 | 0.96 | 1.09 | 1.00 | | | | |
| col8a1a | collagen, type VIII, alpha 1a | ENSDARG00000077403 | 1.00 | 19.40 | 1.00 | 14.29 | | | | |
| col4a1 | collagen, type IV, alpha 1 | ENSDARG00000055009 | 1.00 | 1.04 | 0.83 | 1.25 | | | | |
| col5a1 | procollagen, type V, alpha 1 | ENSDARG00000012593 | 0.91 | 0.84 | 0.85 | 0.90 | | | | |
| col28a2a | collagen type XXVIII alpha 1 a | ENSDARG00000076321 | 0.88 | 0.77 | 1.04 | 0.65 | | | | |
| col15a1b | collagen, type XV, alpha 1b | ENSDARG00000061848 | 0.84 | 0.58 | 1.29 | 0.38 | | | | |
| col1a2 | collagen, type I, alpha 2 | ENSDARG00000020007 | 0.77 | 0.89 | 0.90 | 0.76 | | | | |
| col1a1b | collagen, type I, alpha 1b | ENSDARG00000035809 | 0.76 | 1.17 | 1.22 | 0.73 | | | | |
| col14a1a | collagen, type XIV, alpha 1a | ENSDARG00000005762 | 0.74 | 1.17 | 1.08 | 0.80 | | | | |
| col4a6 | collagen, type IV, alpha 6 | ENSDARG00000052061 | 0.72 | 1.49 | 1.35 | 0.80 | | | | |
| col1a1a | collagen, type I, alpha 1a | ENSDARG00000012405 | 0.72 | 1.01 | 1.07 | 0.68 | | | | |
| col4a5 | collagen, type IV, alpha 5 (Alport syndrome) | ENSDARG00000052063 | 0.68 | 1.29 | 1.06 | 0.83 | | | | |
| col10a1a | collagen, type X, alpha 1a | ENSDARG00000054753 | 0.50 | 0.57 | 0.54 | 0.53 | | | | |
| col9a2 | procollagen, type IX, alpha 2 | ENSDARG00000024492 | 0.40 | 0.84 | 0.90 | 0.37 | | | | |
| col9a3 | collagen, type IX, alpha 3 | ENSDARG00000037845 | 0.40 | 0.97 | 1.17 | 0.33 | | | | |
| col9a1b | collagen, type IX, alpha 1b | ENSDARG00000031483 | 0.35 | 1.13 | 1.00 | 0.40 | | | | |
| col11a1a | collagen, type XI, alpha 1a | ENSDARG00000026165 | 0.33 | 1.05 | 0.76 | 0.46 | | | | |
| col2a1a | collagen, type II, alpha 1a | ENSDARG00000069093 | 0.32 | 1.77 | 1.51 | 0.37 | | | | |
| col11a2 | collagen, type XI, alpha 2 | ENSDARG00000012422 | 0.30 | 1.16 | 0.92 | 0.38 | | | | |
| col7a1 | collagen, type VII, alpha 1 | ENSDARG00000021720 | 0.21 | 2.36 | 0.84 | 0.60 | | | | |
| ctsl.1 | cathepsin L.1 | ENSDARG00000003902 | 11.91 | 0.91 | 1.28 | 8.44 | | | | |
| ctss2.1 | cathepsin Sb, tandem duplicate 1 | ENSDARG00000074656 | 6.91 | 2.06 | 2.15 | 6.61 | | | | |
| ctsk | cathepsin K | ENSDARG00000040251 | 4.58 | 0.82 | 1.46 | 2.58 | | | | |
| ctss2.2 | cathepsin Sb, tandem duplicate 2 | ENSDARG00000013771 | 3.58 | 1.46 | 3.70 | 1.42 | | | | |
| ctsh | cathepsin H | ENSDARG00000041108 | 2.99 | 0.86 | 1.41 | 1.83 | | | | |
| ctsba | cathepsin Ba | ENSDARG00000055120 | 2.29 | 1.16 | 2.44 | 1.09 | | | | |
| ctsc | cathepsin C | ENSDARG00000101334 | 2.27 | 0.90 | 1.01 | 2.01 | | | | |
| ctsz | cathepsin Z | ENSDARG00000043081 | 2.02 | 0.79 | 1.08 | 1.48 | | | | |
| ctsla | cathepsin La | ENSDARG00000007836 | 1.94 | 1.12 | 1.43 | 1.51 | | | | |
| ctsd | cathepsin D | ENSDARG00000057698 | 1.67 | 1.09 | 1.24 | 1.46 | | | | |
| ctss1 | cathepsin Sa | ENSDARG00000036940 | 1.07 | 1.04 | 1.24 | 0.89 | | | | |
| ctsll | cathepsin L, like | ENSDARG00000011701 | 0.73 | 1.17 | 5.20 | 0.16 | | | | |
| ctsf | cathepsin F | ENSDARG00000063095 | 0.70 | 1.06 | 1.05 | 0.72 | | | | |
| cyba | cytochrome b-245, alpha polypeptide | ENSDARG00000018283 | 3.59 | 1.17 | 1.66 | 2.54 | | | | |
| cybb | cytochrome b-245, beta polypeptide (chronic | ENSDARG00000056615 | 3.51 | 0.58 | 1.17 | 1.74 | | | | |
| cyb5b | cytochrome b5 type B | ENSDARG00000099774 | 2.36 | 0.75 | 0.83 | 2.14 | | | | |
| cyb5d1 | cytochrome b5 domain containing 1 | ENSDARG00000056007 | 1.66 | 1.03 | 0.37 | 4.66 | | | | |
| cyb5r3 | cytochrome b5 reductase 3 | ENSDARG00000005891 | 1.65 | 1.11 | 1.74 | 1.05 | | | | |
| cyb5a | cytochrome b5 type A (microsomal) | ENSDARG00000098589 | 1.51 | 1.30 | 1.59 | 1.23 | | | | |

Fold change
>5.0 = red
>3.0 = light red
>1.2 = pale red
<0.85 = pale blue
<0.65 = light blue
<0.5 = blue

Gene counts
>1
>10
>100
>1000

p<0.05 p<0.1

| Gene Symbol | Gene Title | ensemble | (1) DMSO (operated vs unoperated) | (2) Operated (SB431542 vs DMSO) | (3) Unoperated (SB431542 vs DMSO) | (4) SB431542 (operated vs unoperated) | DMSO unoperated COUNT | DMSO operated COUNT | SB43 unoperated COUNT | SB43 operated COUNT |
|---|---|---|---|---|---|---|---|---|---|---|
| cyb5r1 | cytochrome b5 reductase 1 | ENSDARG00000018966 | 1.06 | 1.11 | 1.10 | 1.07 | | | | |
| dnajb9b | | ENSDARG00000016886 | 3.27 | 1.16 | 0.50 | 7.50 | | | | |
| dnajc25 | | ENSDARG00000067613 | 2.75 | 4.37 | 1.00 | 12.25 | | | | |
| dnajc3a | DnaJ (Hsp40) homolog, subfamily C, member | ENSDARG00000041110 | 2.05 | 0.80 | 0.98 | 1.67 | | | | |
| dnajb11 | DnaJ heat shock protein family (Hsp40) | ENSDARG00000015088 | 1.98 | 0.74 | 0.79 | 1.85 | | | | |
| dnajc12 | DnaJ (Hsp40) homolog, subfamily C, member | ENSDARG00000086691 | 1.72 | 1.44 | 2.93 | 0.84 | | | | |
| dnajc5b | | ENSDARG00000058147 | 1.65 | 1.14 | 1.64 | 1.14 | | | | |
| dnajb6b | DnaJ (Hsp40) homolog, subfamily B, member | ENSDARG00000020953 | 1.61 | 0.86 | 1.17 | 1.19 | | | | |
| dnajc9 | DnaJ (Hsp40) homolog, subfamily C, member | ENSDARG00000031293 | 1.56 | 0.62 | 0.73 | 1.33 | | | | |
| dnajc16 | | ENSDARG00000059699 | 1.40 | 0.68 | 0.17 | 5.79 | | | | |
| dnaja1 | DnaJ (Hsp40) homolog, subfamily A, member 1 | ENSDARG00000030972 | 1.34 | 0.93 | 0.98 | 1.28 | | | | |
| dnajb12a | DnaJ (Hsp40) homolog, subfamily B, member | ENSDARG00000039363 | 1.33 | 1.03 | 1.73 | 0.80 | | | | |
| dnajc3b | DnaJ (Hsp40) homolog, subfamily C, member | ENSDARG00000017874 | 1.30 | 1.00 | 0.96 | 1.35 | | | | |
| dnajb1a | DnaJ (Hsp40) homolog, subfamily B, member | ENSDARG00000099383 | 1.28 | 0.96 | 1.03 | 1.19 | | | | |
| dnajc10 | DnaJ (Hsp40) homolog, subfamily C, member | ENSDARG00000074727 | 1.26 | 0.76 | 0.76 | 1.26 | | | | |
| dnaja2b | DnaJ (Hsp40) homolog, subfamily A, member | ENSDARG00000010745 | 1.26 | 0.76 | 1.02 | 0.94 | | | | |
| dnajc8 | DnaJ (Hsp40) homolog, subfamily C, member | ENSDARG00000059373 | 1.26 | 0.60 | 0.68 | 1.11 | | | | |
| dnajb1b | DnaJ (Hsp40) homolog, subfamily B, member | ENSDARG00000041394 | 1.24 | 6.57 | 4.66 | 1.75 | | | | |
| dnajc5gb | DnaJ (Hsp40) homolog, subfamily C, member | ENSDARG00000017687 | 1.22 | 2.53 | 2.48 | 1.25 | | | | |
| dnajc17 | DnaJ (Hsp40) homolog, subfamily C, member | ENSDARG00000104959 | 1.16 | 0.55 | 0.63 | 1.01 | | | | |
| dnajb12b | | ENSDARG00000087473 | 1.15 | 1.26 | 1.09 | 1.33 | | | | |
| dnajc21 | DnaJ (Hsp40) homolog, subfamily C, member | ENSDARG00000105195 | 1.14 | 0.72 | 0.77 | 1.06 | | | | |
| dnajc16l | | ENSDARG00000060725 | 1.12 | 1.12 | 1.47 | 0.86 | | | | |
| dnaja3a | DnaJ (Hsp40) homolog, subfamily A, member | ENSDARG00000058494 | 1.11 | 0.90 | 1.15 | 0.86 | | | | |
| dnaja2a | DnaJ (Hsp40) homolog, subfamily A, member 2 | ENSDARG00000104066 | 1.09 | 0.82 | 0.86 | 1.04 | | | | |
| dnajc11a | DnaJ (Hsp40) homolog, subfamily C, member | ENSDARG00000011196 | 1.04 | 0.94 | 0.97 | 1.00 | | | | |
| dnajc11b | | ENSDARG00000102696 | 1.02 | 1.08 | 1.01 | 1.09 | | | | |
| dnajb14 | | ENSDARG00000069996 | 1.00 | 0.88 | 1.31 | 0.68 | | | | |
| dnajc1 | DnaJ (Hsp40) homolog, subfamily C, member | ENSDARG00000001940 | 0.99 | 0.94 | 0.91 | 1.02 | | | | |
| dnajb2 | | ENSDARG00000058644 | 0.96 | 1.69 | 2.52 | 0.65 | | | | |
| dnajc15 | | ENSDARG00000038309 | 0.92 | 1.38 | 1.39 | 0.92 | | | | |
| dnajc7 | DnaJ (Hsp40) homolog, subfamily C, member | ENSDARG00000058148 | 0.89 | 0.93 | 0.89 | 0.93 | | | | |
| dnajc5ga | DnaJ (Hsp40) homolog, subfamily C, member | ENSDARG00000041896 | 0.89 | 0.71 | 0.87 | 0.72 | | | | |
| dnajc6 | | ENSDARG00000079891 | 0.88 | 1.02 | 1.14 | 0.79 | | | | |
| dnajc27 | DnaJ (Hsp40) homolog, subfamily C, member | ENSDARG00000070916 | 0.84 | 0.95 | 1.23 | 0.65 | | | | |
| dnajc19 | DnaJ (Hsp40) homolog, subfamily C, member | ENSDARG00000044420 | 0.83 | 0.78 | 0.89 | 0.73 | | | | |
| dnajc5ab | DnaJ (Hsp40) homolog, subfamily C, member | ENSDARG00000004836 | 0.83 | 0.82 | 0.80 | 0.85 | | | | |
| dnajc5aa | | ENSDARG00000042948 | 0.82 | 1.15 | 1.14 | 0.82 | | | | |
| dnajb4 | | ENSDARG00000038978 | 0.80 | 1.35 | 1.16 | 0.93 | | | | |
| dnajc24 | DnaJ (Hsp40) homolog, subfamily C, member | ENSDARG00000023927 | 0.79 | 0.68 | 0.59 | 0.89 | | | | |
| dnajc28 | | ENSDARG00000018181 | 0.78 | 1.30 | 1.19 | 0.86 | | | | |
| dnajc4 | | ENSDARG00000024090 | 0.78 | 1.44 | 1.38 | 0.81 | | | | |
| dnaja3b | DnaJ (Hsp40) homolog, subfamily A, member | ENSDARG00000102295 | 0.78 | 2.28 | 2.25 | 0.79 | | | | |
| dnajc2 | | ENSDARG00000070477 | 0.71 | 0.94 | 0.74 | 0.90 | | | | |
| dnajc14 | | ENSDARG00000105398 | 0.67 | 2.08 | 0.35 | 4.02 | | | | |
| dnajb6a | DnaJ (Hsp40) homolog, subfamily B, member | ENSDARG00000004680 | 0.67 | 2.48 | 2.72 | 0.61 | | | | |
| dnajb13 | | ENSDARG00000043157 | 0.66 | 1.38 | 0.81 | 1.14 | | | | |
| dnajc22 | | ENSDARG00000037067 | 0.57 | 2.10 | 0.49 | 2.44 | | | | |
| dnajc30b | DnaJ (Hsp40) homolog, subfamily C, member | ENSDARG00000079415 | 0.56 | 1.17 | 2.49 | 0.26 | | | | |
| dnajc18 | DnaJ (Hsp40) homolog, subfamily C, member | ENSDARG00000056005 | 0.47 | 1.19 | 0.97 | 0.58 | | | | |
| dnajb1 | DnaJ (Hsp40) homolog, subfamily B, member 1 | ENSDARG00000015831 | | | | | | | | |

Fold change
>5.0 = red
>3.0 = light red
>1.2 = pale red
<0.85 = pale blue
<0.65 = light blue
<0.5 = blue

Gene counts
>1
>10
>100
>1000

p<0.05 p<0.1

| Gene Symbol | Gene Title | ensemble | (1) DMSO (operated vs unoperated) | (2) Operated (SB431542 vs DMSO) | (3) Unoperated (SB431542 vs DMSO) | (4) SB431542 (operated vs unoperated) | DMSO unoperated COUNT | DMSO operated COUNT | SB43 unoperated COUNT | SB43 operated COUNT |
|---|---|---|---|---|---|---|---|---|---|---|
| hoxc5a | homeobox C5a | ENSDARG00000070340 | 8.15 | 1.47 | 1.00 | 12.17 | | | | |
| hoxa1a | homeobox A1a | ENSDARG00000104307 | 7.96 | 1.06 | 1.00 | 8.62 | | | | |
| hoxa13b | homeobox A13b | ENSDARG00000036254 | 1.80 | 0.99 | 1.35 | 1.32 | | | | |
| hoxd13a | homeobox D13a | ENSDARG00000059256 | 1.70 | 0.84 | 1.72 | 0.82 | | | | |
| hoxa11a | homeobox A11a | ENSDARG00000104162 | 1.58 | 1.69 | 1.99 | 1.34 | | | | |
| hoxb7a | homeobox B7a | ENSDARG00000056030 | 1.53 | 1.24 | 2.61 | 0.73 | | | | |
| hoxd12a | homeobox D12a | ENSDARG00000059263 | 1.52 | 1.57 | 2.63 | 0.91 | | | | |
| hoxc13a | homeobox C13a | ENSDARG00000070353 | 1.37 | 0.88 | 0.73 | 1.66 | | | | |
| hoxb8b | homeobox B8b | ENSDARG00000054025 | 1.31 | 0.72 | 2.58 | 0.37 | | | | |
| hoxb2a | homeobox B2a | ENSDARG00000000175 | 1.31 | 1.02 | 1.37 | 0.97 | | | | |
| hoxb2a | homeobox B2a | ENSDARG00000000175 | 1.31 | 1.02 | 1.37 | 0.97 | | | | |
| hoxb6b | homeobox B6b | ENSDARG00000026513 | 1.26 | 1.13 | 1.28 | 1.11 | | | | |
| hoxa9b | homeobox A9b | ENSDARG00000056819 | 1.15 | 1.04 | 1.59 | 0.76 | | | | |
| hoxa2b | homeobox A2b | ENSDARG00000023031 | 1.13 | 1.14 | 1.14 | 1.13 | | | | |
| hoxa9a | homeobox A9a | ENSDARG00000105013 | 1.09 | 1.15 | 1.51 | 0.83 | | | | |
| hoxb4a | homeobox B4a | ENSDARG00000013533 | 1.00 | 2.31 | 1.00 | 1.70 | | | | |
| hoxc4a | homeobox C4a | ENSDARG00000070338 | 1.00 | 1.00 | 1.91 | 0.45 | | | | |
| hoxa10b | homeobox A10b | ENSDARG00000031337 | 1.00 | 0.87 | 1.07 | 0.81 | | | | |
| hoxd10a | homeobox D10a | ENSDARG00000057859 | 0.94 | 1.04 | 1.08 | 0.90 | | | | |
| hoxc12b | homeobox C12b | ENSDARG00000103133 | 0.93 | 0.89 | 0.88 | 0.93 | | | | |
| hoxc9a | homeobox C9a | ENSDARG00000092809 | 0.91 | 1.40 | 0.70 | 1.81 | | | | |
| hoxb8a | homeobox B8a | ENSDARG00000056027 | 0.90 | 0.83 | 0.98 | 0.77 | | | | |
| hoxb5b | homeobox B5b | ENSDARG00000054030 | 0.88 | 0.71 | 0.63 | 0.99 | | | | |
| hoxb5a | homeobox B5a | ENSDARG00000013057 | 0.87 | 0.97 | 0.97 | 0.86 | | | | |
| hoxc11b | homeobox C11b | ENSDARG00000102631 | 0.86 | 1.14 | 1.19 | 0.82 | | | | |
| hoxd9a | homeobox D9a | ENSDARG00000059274 | 0.84 | 0.63 | 0.75 | 0.72 | | | | |
| hoxb3a | homeobox B3a | ENSDARG00000029263 | 0.82 | 0.86 | 1.01 | 0.70 | | | | |
| hoxd3a | homeobox D3a | ENSDARG00000059280 | 0.82 | 1.56 | 0.86 | 1.47 | | | | |
| hoxb6a | homeobox B6a | ENSDARG00000010630 | 0.79 | 0.73 | 0.59 | 0.98 | | | | |
| hoxa11b | homeobox A11b | ENSDARG00000007009 | 0.77 | 0.97 | 0.86 | 0.86 | | | | |
| hoxa4a | homeobox A3a /// homeobox A4a | ENSDARG00000103862 | 0.72 | 0.94 | 0.88 | 0.76 | | | | |
| hoxd11a | homeobox D11a | ENSDARG00000059267 | 0.70 | 0.95 | 1.28 | 0.52 | | | | |
| hoxc6b | homeobox C6b | ENSDARG00000101954 | 0.67 | 1.23 | 0.86 | 0.97 | | | | |
| hoxd4a | homeobox D4a | ENSDARG00000059276 | 0.67 | 1.18 | 0.94 | 0.84 | | | | |
| hoxc10a | homeobox C10a | ENSDARG00000070348 | 0.65 | 0.94 | 0.83 | 0.74 | | | | |
| hoxc6a | homeobox C6a | ENSDARG00000070343 | 0.59 | 1.20 | 0.59 | 1.19 | | | | |
| hoxc8a | homeobox C8a | ENSDARG00000070346 | 0.49 | 1.16 | 0.46 | 1.24 | | | | |
| hsp90b1 | heat shock protein 90, beta (grp94), member 1 | ENSDARG00000114206 | 11.57 | 0.87 | 1.04 | 9.60 | | | | |
| hsp90b1 | heat shock protein 90, beta (grp94), member 1 | ENSDARG00000003570 | 2.00 | 0.95 | 1.02 | 1.86 | | | | |
| hspa4a | heat shock protein 4a | ENSDARG00000004754 | 1.69 | 0.87 | 0.93 | 1.57 | | | | |
| hspd1 | heat shock 60 protein 1 | ENSDARG00000056160 | 1.55 | 1.06 | 1.24 | 1.33 | | | | |
| hspa4b | heat shock protein 4b | ENSDARG00000018989 | 1.50 | 0.69 | 0.87 | 1.18 | | | | |
| hspa5 | heat shock protein 5 | ENSDARG00000103846 | 1.45 | 0.86 | 0.99 | 1.25 | | | | |
| hspa5 | heat shock protein 5 | ENSDARG00000103846 | 1.45 | 0.86 | 0.99 | 1.25 | | | | |
| hspb1 | heat shock protein, alpha-crystallin-related, 1 | ENSDARG00000041065 | 1.34 | 0.78 | 0.90 | 1.16 | | | | |
| hsp90aa1.2 | heat shock protein 90, alpha (cytosolic), class | ENSDARG00000024746 | 1.33 | 0.87 | 0.84 | 1.38 | | | | |
| hspa14 | heat shock protein 14 | ENSDARG00000058030 | 1.23 | 0.75 | 0.88 | 1.05 | | | | |
| hspe1 | heat shock 10 protein 1 | ENSDARG00000056167 | 1.17 | 0.91 | 0.99 | 1.08 | | | | |
| hspa9 | heat shock protein 9 | ENSDARG00000003035 | 1.14 | 1.29 | 1.60 | 0.92 | | | | |
| hspa8 | heat shock protein 8 | ENSDARG00000068992 | 1.11 | 0.94 | 0.93 | 1.13 | | | | |
| hsp90ab1 | heat shock protein 90, alpha (cytosolic), class | ENSDARG00000029150 | 0.91 | 0.72 | 0.66 | 0.99 | | | | |

Fold change
>5.0 = red
>3.0 = light red
>1.2 = pale red
<0.85 = pale blue
<0.65 = light blue
<0.5 = blue
p<0.05 p<0.1

Gene counts
>1
>10
>100
>1000

| Gene Symbol | Gene Title | ensemble | (1) DMSO (operated vs unoperated) | (2) Operated (SB431542 vs DMSO) | (3) Unoperated (SB431542 vs DMSO) | (4) SB431542 (operated vs unoperated) | DMSO unoperated COUNT | DMSO operated COUNT | SB43 unoperated COUNT | SB43 operated COUNT |
|---|---|---|---|---|---|---|---|---|---|---|
| hspb6 | heat shock protein, alpha-crystallin-related, b6 | ENSDARG00000077236 | 0.91 | 1.55 | 1.44 | 0.97 | | | | |
| hspbp1 | HSPA (heat shock 70kDa) binding protein, | ENSDARG00000102937 | 0.84 | 1.02 | 1.28 | 0.67 | | | | |
| hspb8 | heat shock protein b8 | ENSDARG00000058365 | 0.78 | 1.14 | 1.09 | 0.82 | | | | |
| hsp90aa1.1 | heat shock protein 90, alpha (cytosolic), class | ENSDARG00000010478 | 0.74 | 0.91 | 1.02 | 0.66 | | | | |
| hspa4l | heat shock protein 4 like | ENSDARG00000053544 | 0.70 | 1.11 | 0.90 | 0.86 | | | | |
| hspa12a | heat shock protein 12A | ENSDARG00000070603 | 0.65 | 0.98 | 1.06 | 0.60 | | | | |
| hspb15 | heat shock protein, alpha-crystallin-related, b15 | ENSDARG00000078411 | 0.57 | 1.20 | 0.87 | 0.79 | | | | |
| igfbp5a | | ENSDARG00000039264 | 2.75 | 0.12 | 1.52 | 0.22 | | | | |
| igfbp1a | insulin-like growth factor binding protein 1a | ENSDARG00000099351 | 2.01 | 1.59 | 1.27 | 2.53 | | | | |
| igfbp1b | insulin-like growth factor binding protein 1b | ENSDARG00000038666 | 1.36 | 5.17 | 1.21 | 5.85 | | | | |
| igfbp6a | | ENSDARG00000070941 | 1.15 | 0.41 | 1.60 | 0.29 | | | | |
| igfbp7 | | ENSDARG00000104138 | 1.00 | 2.31 | 1.91 | 0.87 | | | | |
| igfbp6b | insulin-like growth factor binding protein 6b | ENSDARG00000090833 | 0.90 | 0.50 | 0.63 | 0.71 | | | | |
| igfbp5b | insulin-like growth factor binding protein 5b | ENSDARG00000025348 | 0.87 | 1.89 | 1.72 | 0.96 | | | | |
| igfbp2b | | ENSDARG00000031422 | 0.85 | 0.68 | 0.13 | 4.62 | | | | |
| igfbp3 | insulin-like growth factor binding protein 3 | ENSDARG00000099144 | 0.82 | 3.38 | 3.79 | 0.73 | | | | |
| igfbp2a | insulin-like growth factor binding protein 2a | ENSDARG00000052470 | 0.75 | 0.80 | 1.09 | 0.55 | | | | |
| krt93 | xkeratin 93 | ENSDARG00000044976 | 5.95 | 0.14 | 1.00 | 1.00 | | | | |
| krt18a.1 | keratin 18 | ENSDARG00000018404 | 5.05 | 0.75 | 1.18 | 3.20 | | | | |
| krt96 | keratin 96 | ENSDARG00000095147 | 2.86 | 0.19 | 0.92 | 0.59 | | | | |
| krt97 | keratin 97 | ENSDARG00000000212 | 2.66 | 0.30 | 0.74 | 1.08 | | | | |
| krt94 | keratin 94 | ENSDARG00000044975 | 1.93 | 0.64 | 0.54 | 2.31 | | | | |
| krt18b | keratin zgc:77517 krtt1c6 | ENSDARG00000028618 | 1.91 | 0.86 | 0.93 | 1.76 | | | | |
| krt8 | keratin 8 | ENSDARG00000058358 | 1.67 | 0.75 | 0.95 | 1.32 | | | | |
| krt1-c5 | keratin 1 | ENSDARG00000026979 | 1.22 | 0.71 | 0.74 | 1.17 | | | | |
| krt222 | keratin 222 | ENSDARG00000071518 | 1.18 | 0.90 | 1.25 | 0.85 | | | | |
| krt99 | keratin 99 zgc:110712 | ENSDARG00000019365 | 0.94 | 0.58 | 1.41 | 0.39 | | | | |
| krt98 | keratin 98 | ENSDARG00000044973 | 0.71 | 0.31 | 0.66 | 0.33 | | | | |
| krt5 | keratin 5 | ENSDARG00000058371 | 0.56 | 1.44 | 1.30 | 0.62 | | | | |
| krt92 | keratin 96? | ENSDARG00000036834 | 0.53 | 2.05 | 1.01 | 1.08 | | | | |
| krt4 | keratin 4 | ENSDARG00000017624 | 0.49 | 1.33 | 0.99 | 0.66 | | | | |
| krtt1c19e | keratin type 1 c19e | ENSDARG00000090268 | 0.47 | 0.96 | 0.90 | 0.51 | | | | |
| krt91 | keratin 91 | ENSDARG00000036830 | 0.47 | 1.19 | 0.83 | 0.67 | | | | |
| krt17 | keratin 17 | ENSDARG00000094041 | 0.46 | 1.24 | 0.85 | 0.68 | | | | |
| krt15 | keratin 15 | ENSDARG00000036840 | 0.36 | 1.35 | 0.65 | 0.74 | | | | |
| krt1-19d | keratin, type 1, gene 19d | ENSDARG00000023082 | 0.23 | 1.97 | 0.82 | 0.55 | | | | |
| map1lc3c | microtubule-associated protein 1 light chain 3 | ENSDARG00000100528 | 6.03 | 0.14 | 0.25 | 3.45 | | | | |
| mapre1b | microtubule-associated protein, RP/EB family, | ENSDARG00000002659 | 1.67 | 0.67 | 0.91 | 1.24 | | | | |
| map1lc3a | microtubule-associated protein 1 light chain 3 | ENSDARG00000033609 | 1.37 | 1.30 | 1.89 | 0.94 | | | | |
| mapre1a | microtubule-associated protein, RP/EB family, | ENSDARG00000042927 | 1.14 | 1.11 | 1.11 | 1.14 | | | | |
| map1lc3b | microtubule-associated protein 1 light chain 3 | ENSDARG00000101127 | 1.02 | 1.01 | 1.00 | 1.03 | | | | |
| map9 | microtubule-associated protein 9 | ENSDARG00000037276 | 1.02 | 1.08 | 0.65 | 1.68 | | | | |
| map1sa | microtubule-associated protein 1Sa | ENSDARG00000060805 | 0.79 | 0.86 | 0.84 | 0.81 | | | | |
| maptb | microtubule-associated protein tau b | ENSDARG00000087616 | 0.73 | 0.58 | 0.73 | 0.58 | | | | |
| map1ab | microtubule-associated protein 1Ab | ENSDARG00000022045 | 0.62 | 0.94 | 0.84 | 0.69 | | | | |
| mapre3b | microtubule-associated protein, RP/EB family, | ENSDARG00000102878 | 0.58 | 1.01 | 0.93 | 0.64 | | | | |
| map1lc3cl | microtubule-associated protein 1 light chain 3 | ENSDARG00000075727 | 0.29 | 0.82 | 1.08 | 0.22 | | | | |
| mmp13b | matrix metallopeptidase 13b | ENSDARG00000100794 | 78.68 | 0.94 | 6.93 | 10.68 | | | | |
| mmp9 | matrix metallopeptidase 9 | ENSDARG00000042816 | 10.35 | 2.42 | 3.37 | 7.44 | | | | |

Fold change
>5.0 = red
>3.0 = light red
>1.2 = pale red
<0.85 = pale blue
<0.65 = light blue
<0.5 = blue

Gene counts
>1
>10
>100
>1000

p<0.05 p<0.1

| Gene Symbol | Gene Title | ensemble | (1) DMSO (operated vs unoperated) | (2) Operated (SB431542 vs DMSO) | (3) Unoperated (SB431542 vs DMSO) | (4) SB431542 (operated vs unoperated) | DMSO unoperated COUNT | DMSO operated COUNT | SB43 unoperated COUNT | SB43 operated COUNT |
|---|---|---|---|---|---|---|---|---|---|---|
| mmp13a | matrix metallopeptidase 13a | ENSDARG00000012395 | 7.49 | 1.01 | 1.20 | 6.30 | | | | |
| mmp13a | matrix metallopeptidase 13a | ENSDARG00000114451 | 4.34 | 1.91 | 1.53 | 5.42 | | | | |
| mmp14b | matrix metallopeptidase 14b (membrane- | ENSDARG00000008388 | 2.01 | 0.94 | 1.88 | 1.01 | | | | |
| mmp2 | matrix metallopeptidase 2 | ENSDARG00000017676 | 1.64 | 1.61 | 3.05 | 0.87 | | | | |
| mmp14a | matrix metallopeptidase 14a (membrane- | ENSDARG00000002235 | 1.27 | 0.96 | 1.17 | 1.04 | | | | |
| MMP23B | matrix metallopeptidase 23B | ENSDARG00000043079 | 1.00 | 3.92 | 1.00 | 2.89 | | | | |
| mmp17b | matrix metallopeptidase 17b | ENSDARG00000102956 | 0.88 | 1.28 | 1.18 | 0.96 | | | | |
| mmp30 | matrix metallopeptidase 30 | ENSDARG00000045887 | 0.62 | 2.44 | 1.22 | 1.25 | | | | |
| ms4a17a.4 | membrane-spanning 4-domains | ENSDARG00000014024 | 9.71 | 0.70 | 1.04 | 6.57 | | | | |
| ms4a17a.5 | membrane-spanning 4-domains | ENSDARG00000092204 | 6.24 | 0.67 | 1.30 | 3.21 | | | | |
| ms4a17a.2 | membrane-spanning 4-domains | ENSDARG00000105674 | 5.82 | 0.70 | 0.96 | 4.23 | | | | |
| ms4a17a.7 | membrane-spanning 4-domains | ENSDARG00000043796 | 5.36 | 0.81 | 1.65 | 2.64 | | | | |
| ms4a17a.1 | membrane-spanning 4-domains, subfamily A, | ENSDARG00000043798 | 4.40 | 0.57 | 0.48 | 5.14 | | | | |
| ms4a17a.12 | membrane-spanning 4-domains | ENSDARG00000053563 | 2.74 | 0.86 | 1.04 | 2.27 | | | | |
| ms4a17a.11 | membrane-spanning 4-domains | ENSDARG00000094809 | 2.71 | 0.88 | 0.86 | 2.77 | | | | |
| ms4a17a.9 | membrane-spanning 4-domains | ENSDARG00000094854 | 2.32 | 0.87 | 1.23 | 1.64 | | | | |
| mybphb | myosin binding protein Hb | ENSDARG00000003081 | 0.99 | 2.04 | 2.21 | 0.92 | | | | |
| mybpc2b | myosin binding protein C, fast type b | ENSDARG00000021265 | 0.88 | 0.66 | 0.77 | 0.75 | | | | |
| mybpc3 | myosin binding protein C, cardiac | ENSDARG00000011615 | 0.63 | 0.98 | 1.02 | 0.61 | | | | |
| mybpc1 | myosin binding protein C1 | ENSDARG00000045560 | 0.52 | 0.71 | 0.82 | 0.45 | | | | |
| mybpha | myosin binding protein Ha | ENSDARG00000058799 | 0.33 | 1.33 | 0.99 | 0.44 | | | | |
| myl12.1 | myosin, light chain 12, genome duplicate 1 | ENSDARG00000099766 | 1.56 | 1.18 | 1.55 | 1.19 | | | | |
| mylipa | myosin regulatory light chain interacting protein | ENSDARG00000008859 | 1.01 | 1.45 | 1.45 | 1.01 | | | | |
| myl7 | myosin, light chain 7, regulatory | ENSDARG00000019096 | 1.00 | 2.31 | 1.91 | 0.87 | | | | |
| myl12.2 | myosin, light chain 12, genome duplicate 2 | ENSDARG00000025326 | 0.99 | 0.84 | 0.95 | 0.87 | | | | |
| myl6 | myosin, light chain 6, alkali, smooth muscle | ENSDARG00000008494 | 0.86 | 0.92 | 0.87 | 0.91 | | | | |
| myl10 | myosin, light chain 10, regulatory | ENSDARG00000062592 | 0.77 | 0.75 | 0.91 | 0.64 | | | | |
| myl1 | myosin, light chain 1, alkali; skeletal, fast | ENSDARG00000014196 | 0.73 | 0.80 | 0.99 | 0.59 | | | | |
| mylpfb | myosin light chain, phosphorylatable, fast | ENSDARG00000002589 | 0.72 | 1.06 | 1.25 | 0.61 | | | | |
| myl13 | myosin, light chain 13 | ENSDARG00000042245 | 0.69 | 1.07 | 1.02 | 0.72 | | | | |
| mylpfa | myosin light chain, phosphorylatable, fast | ENSDARG00000053254 | 0.62 | 1.22 | 1.06 | 0.72 | | | | |
| mylz3 | myosin, light polypeptide 3, skeletal muscle | ENSDARG00000017441 | 0.61 | 0.99 | 0.94 | 0.64 | | | | |
| mylk4a | myosin light chain kinase 3-like | ENSDARG00000091260 | 0.54 | 2.73 | 0.76 | 1.93 | | | | |
| ndufs1 | NADH dehydrogenase (ubiquinone) Fe-S | ENSDARG00000028546 | 4.06 | 0.21 | 1.00 | 1.00 | | | | |
| ndufa4 | NADH dehydrogenase (ubiquinone) 1 alpha | ENSDARG00000056108 | 1.03 | 2.49 | 3.37 | 0.76 | | | | |
| ndufaf4 | NADH dehydrogenase (ubiquinone) complex I, | ENSDARG00000077859 | 0.97 | 1.16 | 1.29 | 0.88 | | | | |
| ndufs8a | NADH dehydrogenase (ubiquinone) Fe-S | ENSDARG00000051986 | 0.93 | 1.03 | 1.14 | 0.84 | | | | |
| ndufs5 | NADH dehydrogenase (ubiquinone) Fe-S | ENSDARG00000006290 | 0.90 | 0.80 | 0.88 | 0.82 | | | | |
| ndufb4 | NADH dehydrogenase (ubiquinone) 1 beta | ENSDARG00000019332 | 0.86 | 1.36 | 1.36 | 0.87 | | | | |
| ndufb8 | NADH dehydrogenase (ubiquinone) 1 beta | ENSDARG00000010113 | 0.86 | 0.81 | 0.98 | 0.71 | | | | |
| ndufb9 | NADH dehydrogenase (ubiquinone) 1 beta | ENSDARG00000041314 | 0.85 | 0.82 | 0.91 | 0.76 | | | | |
| ndufv2 | NADH dehydrogenase (ubiquinone) | ENSDARG00000013044 | 0.84 | 0.90 | 1.31 | 0.58 | | | | |
| ndufc2 | NADH dehydrogenase (ubiquinone) 1, | ENSDARG00000102115 | 0.84 | 0.94 | 1.05 | 0.75 | | | | |
| ndufa6 | NADH dehydrogenase (ubiquinone) 1 alpha | ENSDARG00000038028 | 0.82 | 0.73 | 0.75 | 0.80 | | | | |
| ndufb10 | NADH dehydrogenase (ubiquinone) 1 beta | ENSDARG00000028889 | 0.82 | 0.96 | 0.99 | 0.79 | | | | |
| ndufaf1 | NADH dehydrogenase (ubiquinone) complex I, | ENSDARG00000025549 | 0.80 | 1.66 | 1.47 | 0.90 | | | | |
| ndufab1a | NADH dehydrogenase (ubiquinone) 1, | ENSDARG00000058463 | 0.79 | 1.19 | 1.10 | 0.85 | | | | |
| ndufv1 | NADH dehydrogenase (ubiquinone) | ENSDARG00000036438 | 0.78 | 1.12 | 1.07 | 0.82 | | | | |
| ndufs2 | NADH dehydrogenase (ubiquinone) Fe-S | ENSDARG00000007526 | 0.78 | 1.03 | 1.04 | 0.77 | | | | |

Fold change
- >5.0 = red
- >3.0 = light red
- >1.2 = pale red
- <0.85 = pale blue
- <0.65 = light blue
- <0.5 = blue

Gene counts
- >1
- >10
- >100
- >1000

p<0.05 p<0.1

| Gene Symbol | Gene Title | ensemble | (1) DMSO (operated vs unoperated) | (2) Operated (SB431542 vs DMSO) | (3) Unoperated (SB431542 vs DMSO) | (4) SB431542 (operated vs unoperated) | DMSO unoperated COUNT | DMSO operated COUNT | SB43 unoperated COUNT | SB43 operated COUNT |
|---|---|---|---|---|---|---|---|---|---|---|
| ndufb5 | NADH dehydrogenase (ubiquinone) 1 beta | ENSDARG00000070824 | 0.76 | 1.13 | 1.26 | 0.69 | | | | |
| ndufaf6 | NADH dehydrogenase (ubiquinone) complex I, | ENSDARG00000053652 | 0.75 | 0.85 | 0.69 | 0.92 | | | | |
| ndufb6 | NADH dehydrogenase (ubiquinone) 1 beta | ENSDARG00000037259 | 0.75 | 1.12 | 1.10 | 0.76 | | | | |
| ndufs6 | NADH dehydrogenase (ubiquinone) Fe-S | ENSDARG00000056583 | 0.75 | 1.46 | 2.00 | 0.54 | | | | |
| ndufa8 | NADH dehydrogenase (ubiquinone) 1 alpha | ENSDARG00000058041 | 0.73 | 1.19 | 1.22 | 0.71 | | | | |
| ndufb3 | NADH dehydrogenase (ubiquinone) 1 beta | ENSDARG00000075709 | 0.72 | 0.86 | 0.89 | 0.70 | | | | |
| ndufb7 | NADH dehydrogenase (ubiquinone) 1 beta | ENSDARG00000033789 | 0.71 | 0.85 | 0.88 | 0.69 | | | | |
| ndufs4 | NADH dehydrogenase (ubiquinone) Fe-S | ENSDARG00000052840 | 0.71 | 0.89 | 0.85 | 0.74 | | | | |
| ndufa10 | NADH dehydrogenase (ubiquinone) 1 alpha | ENSDARG00000013333 | 0.69 | 0.94 | 0.82 | 0.80 | | | | |
| ndufa4l | NADH dehydrogenase (ubiquinone) 1 alpha | ENSDARG00000099499 | 0.68 | 1.13 | 1.03 | 0.74 | | | | |
| NDUFB1 | NADH dehydrogenase (ubiquinone) 1 beta | ENSDARG00000087456 | 0.67 | 1.05 | 0.87 | 0.81 | | | | |
| ndufab1b | NADH dehydrogenase (ubiquinone) 1, | ENSDARG00000014915 | 0.66 | 0.99 | 0.95 | 0.69 | | | | |
| ndufa5 | NADH dehydrogenase (ubiquinone) 1 alpha | ENSDARG00000039346 | 0.65 | 1.16 | 0.96 | 0.79 | | | | |
| ndufb2 | NADH dehydrogenase (ubiquinone) 1 beta | ENSDARG00000045490 | 0.65 | 1.01 | 0.89 | 0.74 | | | | |
| ndufv3 | NADH:ubiquinone oxidoreductase subunit V3 | ENSDARG00000090389 | 0.64 | 0.89 | 0.84 | 0.68 | | | | |
| ndufaf5 | NADH dehydrogenase (ubiquinone) complex I, | ENSDARG00000061629 | 0.64 | 0.90 | 0.77 | 0.75 | | | | |
| ndufa2 | NADH dehydrogenase (ubiquinone) 1 alpha | ENSDARG00000021984 | 0.62 | 0.90 | 0.75 | 0.75 | | | | |
| ndufa11 | NADH dehydrogenase (ubiquinone) 1 alpha | ENSDARG00000042777 | 0.62 | 1.01 | 0.89 | 0.70 | | | | |
| ndufs7 | NADH dehydrogenase (ubiquinone) Fe-S | ENSDARG00000074552 | 0.54 | 1.06 | 0.77 | 0.75 | | | | |
| ndufa1 | NADH dehydrogenase (ubiquinone) 1 alpha | ENSDARG00000036329 | 0.52 | 1.10 | 0.80 | 0.71 | | | | |
| nr5a5 | nuclear receptor subfamily 5, group A, member | ENSDARG00000039116 | 1.60 | 6.65 | 2.67 | 3.98 | | | | |
| nr5a1b | nuclear receptor subfamily 5, group A, member | ENSDARG00000023362 | 1.41 | 2.32 | 4.27 | 0.76 | | | | |
| nr1h3 | nuclear receptor subfamily 1, group H, member | ENSDARG00000098439 | 1.34 | 1.06 | 1.26 | 1.13 | | | | |
| nrip1b | nuclear receptor interacting protein 1b | ENSDARG00000068894 | 1.30 | 1.47 | 2.00 | 0.95 | | | | |
| nrf1 | nuclear respiratory factor 1 | ENSDARG00000000018 | 1.22 | 0.89 | 1.18 | 0.93 | | | | |
| nr2f6a | nuclear receptor subfamily 2, group F, member | ENSDARG00000003607 | 1.19 | 0.69 | 0.96 | 0.85 | | | | |
| nr2f1a | nuclear receptor subfamily 2, group F, member | ENSDARG00000052695 | 1.11 | 0.76 | 1.03 | 0.83 | | | | |
| nrbf2b | nuclear receptor binding factor 2b | ENSDARG00000023591 | 1.05 | 1.10 | 1.14 | 1.01 | | | | |
| nr2f5 | nuclear receptor subfamily 2, group F, member | ENSDARG00000033172 | 1.01 | 1.61 | 1.34 | 1.22 | | | | |
| nr2e1 | nuclear receptor subfamily 2, group E, member | ENSDARG00000017107 | 1.00 | 1.00 | 3.04 | 0.28 | | | | |
| nr2f6b | nuclear receptor subfamily 2, group F, member | ENSDARG00000003165 | 0.98 | 1.04 | 0.98 | 1.04 | | | | |
| nr1h5 | nuclear receptor subfamily 1, group H, member | ENSDARG00000031046 | 0.91 | 2.98 | 3.21 | 0.84 | | | | |
| nr3c1 | nuclear receptor subfamily 3, group C, member | ENSDARG00000025032 | 0.85 | 0.87 | 0.85 | 0.86 | | | | |
| nr2f2 | nuclear receptor subfamily 2, group F, member | ENSDARG00000040926 | 0.78 | 1.08 | 1.08 | 0.78 | | | | |
| nr4a2b | nuclear receptor subfamily 4, group A, member | ENSDARG00000044532 | 0.67 | 1.63 | 1.13 | 0.97 | | | | |
| nr2f1b | nuclear receptor subfamily 2, group F, member | ENSDARG00000017168 | 0.67 | 1.11 | 0.67 | 1.10 | | | | |
| nr1d2b | nuclear receptor subfamily 1, group D, member | ENSDARG00000009594 | 0.64 | 1.23 | 1.94 | 0.41 | | | | |
| nr1d2a | nuclear receptor subfamily 1, group D, member | ENSDARG00000003820 | 0.62 | 1.39 | 1.15 | 0.75 | | | | |
| nr0b2a | nuclear receptor subfamily 0, group B, member | ENSDARG00000044685 | 0.61 | 2.92 | 2.29 | 0.77 | | | | |
| nr2c2 | nuclear receptor subfamily 2, group C, member | ENSDARG00000042477 | 0.53 | 0.97 | 0.93 | 0.55 | | | | |
| nr4a1 | nuclear receptor subfamily 4, group A, member | ENSDARG00000000796 | 0.52 | 3.44 | 2.55 | 0.70 | | | | |
| nr1d1 | nuclear receptor subfamily 1, group d, member | ENSDARG00000033160 | 0.49 | 0.92 | 1.41 | 0.31 | | | | |
| nr6a1a | nuclear receptor subfamily 6, group A, member | ENSDARG00000101508 | 0.47 | 0.61 | 0.33 | 1.00 | | | | |
| nr5a2 | nuclear receptor subfamily 5, group A, member | ENSDARG00000100940 | 0.42 | 1.11 | 0.50 | 0.95 | | | | |
| nr5a1a | nuclear receptor subfamily 5, group A, member | ENSDARG00000103176 | 0.24 | 3.76 | 0.77 | 1.16 | | | | |
| ppp1r18 | protein phosphatase 1, regulatory subunit 18 | ENSDARG00000071251 | 4.57 | 0.55 | 1.16 | 2.15 | | | | |
| ppp1r11 | protein phosphatase 1, regulatory (inhibitor) | ENSDARG00000036063 | 4.50 | 0.72 | 4.57 | 0.71 | | | | |
| ppp2r2ab | protein phosphatase 2, regulatory subunit B, | ENSDARG00000006624 | 2.48 | 1.81 | 1.56 | 2.88 | | | | |
| ppp2r2ab | protein phosphatase 2, regulatory subunit B, | ENSDARG00000006624 | 2.48 | 1.81 | 1.56 | 2.88 | | | | |
| ppp4r2b | protein phosphatase 4, regulatory subunit 2b | ENSDARG00000053447 | 2.15 | 0.79 | 1.56 | 1.08 | | | | |
| ppp1r3ca | protein phosphatase 1, regulatory subunit 3Ca | ENSDARG00000071005 | 2.05 | 1.35 | 0.80 | 3.46 | | | | |

Fold change
- >5.0 = red
- >3.0 = light red
- >1.2 = pale red
- <0.85 = pale blue
- <0.65 = light blue
- <0.5 = blue

p<0.05   p<0.1

Gene counts
- >1
- >10
- >100
- >1000

| Gene Symbol | Gene Title | ensemble | (1) DMSO (operated vs unoperated) | (2) Operated (SB431542 vs DMSO) | (3) Unoperated (SB431542 vs DMSO) | (4) SB431542 (operated vs unoperated) | DMSO unoperated COUNT | DMSO operated COUNT | SB43 unoperated COUNT | SB43 operated COUNT |
|---|---|---|---|---|---|---|---|---|---|---|
| ppp1r3da | protein phosphatase 1, regulatory subunit 3Da | ENSDARG00000077513 | 2.00 | 0.95 | 1.69 | 1.12 | | | | |
| ppp4ca | protein phosphatase 4, catalytic subunit a | ENSDARG00000070570 | 1.86 | 0.91 | 1.54 | 1.10 | | | | |
| ppp4r2a | protein phosphatase 4, regulatory subunit 2a | ENSDARG00000026540 | 1.75 | 0.77 | 1.23 | 1.10 | | | | |
| ppp1caa | protein phosphatase 1, catalytic subunit, alpha | ENSDARG00000003486 | 1.48 | 1.43 | 1.94 | 1.09 | | | | |
| ppp1r15a | protein phosphatase 1, regulatory subunit 15A | ENSDARG00000069135 | 1.42 | 2.59 | 1.58 | 2.33 | | | | |
| ppp1cab | protein phosphatase 1, catalytic subunit, alpha | ENSDARG00000071566 | 1.41 | 0.82 | 1.02 | 1.13 | | | | |
| ppp1r8b | protein phosphatase 1, regulatory subunit 8b | ENSDARG00000022430 | 1.37 | 0.80 | 0.87 | 1.25 | | | | |
| ppp1r14bb | protein phosphatase 1, regulatory (inhibitor) | ENSDARG00000030161 | 1.36 | 1.06 | 1.67 | 0.87 | | | | |
| ppp1r3b | protein phosphatase 1, regulatory subunit 3B | ENSDARG00000044691 | 1.27 | 0.44 | 0.98 | 0.57 | | | | |
| ppp2cb | protein phosphatase 2, catalytic subunit, beta | ENSDARG00000099241 | 1.18 | 0.85 | 0.80 | 1.25 | | | | |
| ppp2r1ba | protein phosphatase 2, regulatory subunit A, | ENSDARG00000007791 | 1.16 | 0.70 | 0.77 | 1.05 | | | | |
| ppp4r3b | protein phosphatase 4, regulatory subunit 3B | ENSDARG00000103415 | 1.16 | 0.63 | 0.50 | 1.45 | | | | |
| ppp1r37 | protein phosphatase 1, regulatory subunit 37 | ENSDARG00000078458 | 1.13 | 1.57 | 1.34 | 1.32 | | | | |
| ppp1r13ba | protein phosphatase 1, regulatory subunit 13Ba | ENSDARG00000004377 | 1.11 | 0.99 | 1.33 | 0.82 | | | | |
| ppp1r13l | protein phosphatase 1, regulatory subunit 13 | ENSDARG00000013777 | 1.09 | 0.82 | 0.74 | 1.21 | | | | |
| ppp6r3 | protein phosphatase 6, regulatory subunit 3 | ENSDARG00000013379 | 1.06 | 1.06 | 1.02 | 1.11 | | | | |
| ppp1r9bb | neurabin-2-like /// protein phosphatase 1, | ENSDARG00000071709 | 1.04 | 0.65 | 0.99 | 0.69 | | | | |
| ppp2r3c | protein phosphatase 2, regulatory subunit B'', | ENSDARG00000043972 | 1.04 | 1.38 | 1.53 | 0.93 | | | | |
| ppp6r2a | protein phosphatase 6, regulatory subunit 2a | ENSDARG00000045540 | 1.03 | 1.14 | 1.12 | 1.04 | | | | |
| ppp2r5d | protein phosphatase 2, regulatory subunit B', | ENSDARG00000014428 | 1.03 | 0.96 | 0.98 | 1.01 | | | | |
| ppp5c | protein phosphatase 5, catalytic subunit | ENSDARG00000034313 | 1.02 | 0.76 | 0.80 | 0.96 | | | | |
| ppp1r2 | protein phosphatase 1, regulatory (inhibitor) | ENSDARG00000054007 | 0.98 | 1.71 | 1.40 | 1.20 | | | | |
| ppp1cb | protein phosphatase 1, catalytic subunit, beta | ENSDARG00000044153 | 0.98 | 1.05 | 1.16 | 0.88 | | | | |
| ppp1r12a | protein phosphatase 1, regulatory subunit 12A | ENSDARG00000010784 | 0.98 | 1.46 | 1.27 | 1.13 | | | | |
| ppp2r1bb | protein phosphatase 2, regulatory subunit A, | ENSDARG00000032430 | 0.97 | 0.87 | 0.85 | 0.99 | | | | |
| ppp2r5eb | protein phosphatase 2, regulatory subunit B', | ENSDARG00000069118 | 0.96 | 1.45 | 1.78 | 0.78 | | | | |
| ppp2r3b | protein phosphatase 2, regulatory subunit B'', | ENSDARG00000076004 | 0.89 | 1.27 | 1.09 | 1.04 | | | | |
| ppp1r7 | protein phosphatase 1, regulatory (inhibitor) | ENSDARG00000009740 | 0.89 | 0.84 | 0.84 | 0.89 | | | | |
| ppp1r10 | protein phosphatase 1, regulatory subunit 10 | ENSDARG00000032651 | 0.89 | 1.20 | 1.01 | 1.06 | | | | |
| ppp3r1b | protein phosphatase 3 (formerly 2B), regulatory | ENSDARG00000069360 | 0.88 | 0.81 | 1.00 | 0.71 | | | | |
| ppp1r14ab | protein phosphatase 1, regulatory (inhibitor) | ENSDARG00000017710 | 0.86 | 1.02 | 0.96 | 0.91 | | | | |
| ppp1r14aa | protein phosphatase 1, regulatory (inhibitor) | ENSDARG00000011239 | 0.82 | 0.94 | 1.17 | 0.66 | | | | |
| ppp1r42 | protein phosphatase 1, regulatory subunit 42 | ENSDARG00000057632 | 0.80 | 0.70 | 1.03 | 0.54 | | | | |
| ppp3cb | protein phosphatase 3, catalytic subunit, beta | ENSDARG00000025106 | 0.79 | 1.00 | 1.09 | 0.72 | | | | |
| ppp6c | protein phosphatase 6, catalytic subunit | ENSDARG00000002949 | 0.78 | 2.39 | 1.78 | 1.05 | | | | |
| ppp2r5ea | protein phosphatase 2, regulatory subunit B', | ENSDARG00000015474 | 0.77 | 1.13 | 0.98 | 0.88 | | | | |
| ppp4r4 | protein phosphatase 4, regulatory subunit 4 | ENSDARG00000010407 | 0.74 | 2.05 | 1.46 | 1.04 | | | | |
| ppp1r13bb | protein phosphatase 1, regulatory subunit 13Bb | ENSDARG00000009142 | 0.71 | 0.92 | 0.70 | 0.94 | | | | |
| ppp2r2ca | protein phosphatase 2, regulatory subunit B, | ENSDARG00000056797 | 0.71 | 0.87 | 0.55 | 1.12 | | | | |
| ppp1r15b | protein phosphatase 1, regulatory subunit 15B | ENSDARG00000068128 | 0.69 | 2.17 | 0.90 | 1.67 | | | | |
| ppp3cca | protein phosphatase 3, catalytic subunit, | ENSDARG00000014962 | 0.66 | 0.90 | 0.64 | 0.94 | | | | |
| ppp2r5b | protein phosphatase 2, regulatory subunit B', | ENSDARG00000054931 | 0.61 | 1.79 | 0.98 | 1.11 | | | | |
| ppp3ca | protein phosphatase 3, catalytic subunit, alpha | ENSDARG00000004988 | 0.55 | 1.40 | 1.01 | 0.76 | | | | |
| ppp1r3ab | protein phosphatase 1, regulatory subunit 3Ab | ENSDARG00000088813 | 0.54 | 1.52 | 0.86 | 0.96 | | | | |
| ppp2r5ca | protein phosphatase 2, regulatory subunit B', | ENSDARG00000059083 | 0.50 | 2.93 | 0.50 | 2.90 | | | | |
| ppp1r1b | protein phosphatase 1, regulatory (inhibitor) | ENSDARG00000076280 | 0.46 | 0.93 | 0.58 | 0.73 | | | | |
| ppp1r3cb | protein phosphatase 1, regulatory subunit 3Cb | ENSDARG00000014554 | 0.42 | 1.18 | 0.72 | 0.69 | | | | |
| pvalb8 | parvalbumin 8 | ENSDARG00000037790 | 0.94 | 0.82 | 1.00 | 0.78 | | | | |
| pvalb2 | parvalbumin 2 | ENSDARG00000002768 | 0.69 | 1.41 | 1.40 | 0.69 | | | | |
| pvalb4 | parvalbumin 4 | ENSDARG00000024433 | 0.66 | 0.99 | 0.99 | 0.66 | | | | |
| pvalb5 | parvalbumin 5 | ENSDARG00000032836 | 0.66 | 1.01 | 0.91 | 0.73 | | | | |
| pvalb9 | parvalbumin 9 | ENSDARG00000071601 | 0.59 | 0.54 | 0.82 | 0.39 | | | | |

Fold change
>5.0 = red
>3.0 = light red
>1.2 = pale red
<0.85 = pale blue
<0.65 = light blue
<0.5 = blue

Gene counts
>1
>10
>100
>1000

p<0.05 p<0.1

| Gene Symbol | Gene Title | ensemble | (1) DMSO (operated vs unoperated) | (2) Operated (SB431542 vs DMSO) | (3) Unoperated (SB431542 vs DMSO) | (4) SB431542 (operated vs unoperated) | DMSO unoperated COUNT | DMSO operated COUNT | SB43 unoperated COUNT | SB43 operated COUNT |
|---|---|---|---|---|---|---|---|---|---|---|
| pvalb6 | parvalbumin 6 | ENSDARG00000009311 | 0.59 | 1.12 | 0.99 | 0.67 | | | | |
| pvalb3 | parvalbumin 3 | ENSDARG00000022817 | 0.54 | 1.10 | 0.97 | 0.61 | | | | |
| pvalb7 | parvalbumin 7 | ENSDARG00000034705 | 0.52 | 0.86 | 0.88 | 0.51 | | | | |
| pvalb1 | parvalbumin 1 | ENSDARG00000037789 | 0.49 | 1.08 | 0.79 | 0.67 | | | | |
| slc7a7 | solute carrier family 7 (amino acid transporter | ENSDARG00000055226 | 7.70 | 0.67 | 1.00 | 5.15 | | | | |
| slc46a1 | solute carrier family 46 (folate transporter), | ENSDARG00000026149 | 6.06 | 0.87 | 7.83 | 0.67 | | | | |
| slc22a7b.1 | solute carrier family 22 (organic anion | ENSDARG00000056643 | 2.85 | 0.15 | 1.57 | 0.28 | | | | |
| slc39a7 | solute carrier family 39 (zinc transporter), | ENSDARG00000104451 | 2.46 | 0.59 | 0.95 | 1.53 | | | | |
| slc12a8 | solute carrier family 12, member 8 | ENSDARG00000074384 | 2.16 | 1.15 | 1.58 | 1.57 | | | | |
| slc30a1a | solute carrier family 30 (zinc transporter), | ENSDARG00000005463 | 2.13 | 0.83 | 1.17 | 1.52 | | | | |
| slc51a | solute carrier family 51 member A | ENSDARG00000045306 | 2.05 | 0.82 | 0.80 | 2.10 | | | | |
| slc13a2 | solute carrier family 13 (sodium-dependent | ENSDARG00000053853 | 1.98 | 2.55 | 2.32 | 2.17 | | | | |
| slc52a3 | solute carrier family 52 (riboflavin transporter), | ENSDARG00000042737 | 1.97 | 1.51 | 1.39 | 2.15 | | | | |
| slc46a3 | solute carrier family 46, member 3 | ENSDARG00000077313 | 1.96 | 1.48 | 2.10 | 1.39 | | | | |
| slco1d1 | solute carrier organic anion transporter family, | ENSDARG00000104108 | 1.95 | 0.68 | 1.50 | 0.88 | | | | |
| slco2b1 | solute carrier organic anion transporter family, | ENSDARG00000054609 | 1.92 | 1.29 | 1.20 | 2.07 | | | | |
| slc25a19 | solute carrier family 25 (mitochondrial thiamine | ENSDARG00000100385 | 1.89 | 1.76 | 4.65 | 0.71 | | | | |
| slc10a2 | solute carrier family 10 (sodium/bile acid | ENSDARG00000014916 | 1.87 | 1.86 | 2.63 | 1.32 | | | | |
| slc2a10 | solute carrier family 2 (facilitated glucose | ENSDARG00000090820 | 1.82 | 0.54 | 0.69 | 1.43 | | | | |
| slc24a2 | solute carrier family 24 | ENSDARG00000042988 | 1.79 | 1.71 | 1.58 | 1.95 | | | | |
| slc29a3 | solute carrier family 29 (equilibrative nucleoside | ENSDARG00000077828 | 1.77 | 0.92 | 1.05 | 1.55 | | | | |
| slc10a7 | solute carrier family 10, member 7 | ENSDARG00000104508 | 1.74 | 0.69 | 0.95 | 1.27 | | | | |
| slc12a9 | solute carrier family 12, member 9 | ENSDARG00000060366 | 1.74 | 0.90 | 1.48 | 1.05 | | | | |
| slc20a1a | solute carrier family 20, member 1a | ENSDARG00000020114 | 1.73 | 1.10 | 1.80 | 1.06 | | | | |
| slc14a2 | solute carrier family 14 member 2 | ENSDARG00000051914 | 1.70 | 2.45 | 5.38 | 0.78 | | | | |
| slc9a3r1a | solute carrier family 9, subfamily A (NHE3, | ENSDARG00000000068 | 1.67 | 0.57 | 1.04 | 0.92 | | | | |
| slc35f6 | solute carrier family 35, member F6 | ENSDARG00000016745 | 1.66 | 0.85 | 0.94 | 1.52 | | | | |
| slc26a1 | solute carrier family 26 (anion exchanger), | ENSDARG00000029832 | 1.65 | 0.47 | 1.07 | 0.73 | | | | |
| slc12a4 | solute carrier family 12 (potassium/chloride | ENSDARG00000014378 | 1.64 | 1.52 | 1.94 | 1.29 | | | | |
| slc43a3b | solute carrier family 43 member 3b | ENSDARG00000057949 | 1.61 | 1.40 | 1.12 | 2.02 | | | | |
| slc52a2 | solute carrier family 52 (riboflavin transporter), | ENSDARG00000102035 | 1.61 | 0.07 | 0.76 | 0.14 | | | | |
| slc4a1ap | solute carrier family 4 (anion exchanger), | ENSDARG00000034160 | 1.60 | 0.81 | 1.02 | 1.26 | | | | |
| slc2a11b | solute carrier family 2 (facilitated glucose | ENSDARG00000063288 | 1.59 | 0.79 | 1.59 | 0.79 | | | | |
| slc4a4b | solute carrier family 4 (sodium bicarbonate | ENSDARG00000044808 | 1.54 | 0.33 | 0.33 | 1.53 | | | | |
| slc26a11 | solute carrier family 26 (anion exchanger), | ENSDARG00000043021 | 1.48 | 0.71 | 0.78 | 1.34 | | | | |
| slc12a3 | solute carrier family 12 (sodium/chloride | ENSDARG00000013855 | 1.46 | 1.83 | 1.45 | 1.84 | | | | |
| slc39a9 | solute carrier family 39, member 9 | ENSDARG00000070447 | 1.46 | 0.87 | 0.59 | 2.14 | | | | |
| slc35f2 | solute carrier family 35, member F2 | ENSDARG00000069745 | 1.45 | 0.79 | 1.00 | 1.14 | | | | |
| slc31a1 | solute carrier family 31 (copper transporter), | ENSDARG00000013961 | 1.41 | 0.65 | 0.78 | 1.18 | | | | |
| slc35b2 | solute carrier family 35 (adenosine 3'-phospho | ENSDARG00000007886 | 1.41 | 1.47 | 2.30 | 0.90 | | | | |
| slc22a31 | solute carrier family 22, member 31 | ENSDARG00000078882 | 1.40 | 0.77 | 1.12 | 0.97 | | | | |
| slc30a7 | solute carrier family 30 (zinc transporter), | ENSDARG00000019998 | 1.39 | 0.80 | 0.97 | 1.14 | | | | |
| slc35d1a | solute carrier family 35 (UDP-GlcA/UDP- | ENSDARG00000011973 | 1.36 | 0.89 | 1.44 | 0.84 | | | | |
| slc38a2 | solute carrier family 38, member 2 | ENSDARG00000045886 | 1.34 | 0.57 | 0.87 | 0.89 | | | | |
| slc17a9b | solute carrier family 17 (vesicular nucleotide | ENSDARG00000011049 | 1.33 | 0.85 | 0.61 | 1.84 | | | | |
| slc38a7 | solute carrier family 38, member 7 | ENSDARG00000012002 | 1.26 | 0.87 | 0.86 | 1.27 | | | | |
| slc25a33 | solute carrier family 25 (pyrimidine nucleotide | ENSDARG00000039931 | 1.23 | 1.34 | 2.08 | 0.79 | | | | |
| slc16a9a | solute carrier family 16, member 9a | ENSDARG00000013926 | 1.22 | 2.42 | 0.93 | 3.17 | | | | |
| slc39a6 | solute carrier family 39 (zinc transporter), | ENSDARG00000068143 | 1.20 | 0.76 | 0.73 | 1.25 | | | | |
| slc15a4 | solute carrier family 15 (oligopeptide | ENSDARG00000102612 | 1.19 | 0.78 | 1.06 | 0.87 | | | | |
| slc13a5a | info solute carrier family 13 (sodium-dependent | ENSDARG00000077691 | 1.17 | 0.16 | 1.02 | 0.18 | | | | |
| slc40a1 | solute carrier family 40 (iron-regulated | ENSDARG00000000241 | 1.16 | 1.10 | 1.34 | 0.95 | | | | |

Fold change
- >5.0 = red
- >3.0 = light red
- >1.2 = pale red
- <0.85 = pale blue
- <0.65 = light blue
- <0.5 = blue

Gene counts
- >1
- >10
- >100
- >1000

p<0.05 p<0.1

| Gene Symbol | Gene Title | ensemble | (1) DMSO (operated vs unoperated) | (2) Operated (SB431542 vs DMSO) | (3) Unoperated (SB431542 vs DMSO) | (4) SB431542 (operated vs unoperated) | DMSO unoperated COUNT | DMSO operated COUNT | SB43 unoperated COUNT | SB43 operated COUNT |
|---|---|---|---|---|---|---|---|---|---|---|
| slc20a1b | solute carrier family 20 (phosphate transporter), | ENSDARG00000010641 | 1.16 | 1.37 | 1.29 | 1.24 | | | | |
| slc9a7 | solute carrier family 9, subfamily A (NHE7, | ENSDARG00000076754 | 1.15 | 0.57 | 0.82 | 0.80 | | | | |
| slc25a40 | solute carrier family 25 member 40 | ENSDARG00000015856 | 1.15 | 2.26 | 2.27 | 1.14 | | | | |
| slc4a1a | solute carrier family 4 (anion exchanger), | ENSDARG00000012881 | 1.14 | 0.75 | 1.05 | 0.81 | | | | |
| slc12a7a | solute carrier family 12 (potassium/chloride | ENSDARG00000073756 | 1.13 | 1.21 | 1.18 | 1.15 | | | | |
| slc48a1b | solute carrier family 48 (heme transporter), | ENSDARG00000026109 | 1.11 | 1.29 | 2.08 | 0.69 | | | | |
| slc16a9b | solute carrier family 16, member 9b | ENSDARG00000104687 | 1.11 | 1.64 | 2.21 | 0.82 | | | | |
| slc25a20 | solute carrier family 25 (carnitine/acylcarnitine | ENSDARG00000040401 | 1.09 | 0.74 | 0.84 | 0.97 | | | | |
| slc12a7b | solute carrier family 12 (potassium/chloride | ENSDARG00000062058 | 1.09 | 0.97 | 1.19 | 0.89 | | | | |
| slc16a6b | solute carrier family 16, member 6b | ENSDARG00000060246 | 1.07 | 1.48 | 1.30 | 1.22 | | | | |
| slc5a6b | solute carrier family 5 (sodium/multivitamin and | ENSDARG00000014599 | 1.06 | 1.24 | 1.37 | 0.96 | | | | |
| slc6a9 | solute carrier family 6 (neurotransmitter | ENSDARG00000018534 | 1.06 | 1.03 | 1.12 | 0.97 | | | | |
| slc43a2b | solute carrier family 43 (amino acid system L | ENSDARG00000061120 | 1.04 | 1.04 | 1.28 | 0.84 | | | | |
| slc30a5 | solute carrier family 30 (zinc transporter), | ENSDARG00000051921 | 1.03 | 1.20 | 1.31 | 0.95 | | | | |
| slc2a15b | solute carrier family 2 (facilitated glucose | ENSDARG00000053269 | 1.03 | 1.60 | 3.19 | 0.52 | | | | |
| slc35b4 | solute carrier family 35, member B4 | ENSDARG00000071087 | 1.03 | 1.00 | 0.93 | 1.12 | | | | |
| slc16a1b | solute carrier family 16 (monocarboxylate | ENSDARG00000068572 | 1.02 | 0.56 | 0.68 | 0.84 | | | | |
| slc35c2 | solute carrier family 35 (GDP-fucose | ENSDARG00000037517 | 1.02 | 1.14 | 1.27 | 0.92 | | | | |
| slc25a48 | solute carrier family 25, member 48 | ENSDARG00000021250 | 1.01 | 0.56 | 0.83 | 0.68 | | | | |
| slc27a1a | solute carrier family 27 (fatty acid transporter), | ENSDARG00000006240 | 1.01 | 1.15 | 1.58 | 0.73 | | | | |
| slc25a43 | solute carrier family 25, member 43 | ENSDARG00000102048 | 1.00 | 1.62 | 1.76 | 0.92 | | | | |
| slc39a5 | solute carrier family 39 (zinc transporter), | ENSDARG00000079525 | 1.00 | 2.31 | 1.00 | 1.70 | | | | |
| slc25a26 | solute carrier family 25 (S-adenosylmethionine | ENSDARG00000058208 | 1.00 | 0.92 | 0.93 | 0.99 | | | | |
| slc25a55a | solute carrier family 25 (mitochondrial carrier: | ENSDARG00000020893 | 0.99 | 0.86 | 0.88 | 0.97 | | | | |
| slc4a2a | solute carrier family 4 (anion exchanger), | ENSDARG00000028173 | 0.99 | 0.74 | 1.05 | 0.69 | | | | |
| slc25a17 | solute carrier family 25 (mitochondrial carrier; | ENSDARG00000061684 | 0.98 | 0.82 | 0.61 | 1.31 | | | | |
| slc3a2a | solute carrier family 3 (amino acid transporter | ENSDARG00000036427 | 0.97 | 1.10 | 1.26 | 0.85 | | | | |
| slc30a9 | solute carrier family 30 (zinc transporter), | ENSDARG00000057272 | 0.96 | 1.20 | 1.28 | 0.90 | | | | |
| slc37a4a | solute carrier family 37 (glucose-6-phosphate | ENSDARG00000038106 | 0.96 | 0.82 | 0.96 | 0.82 | | | | |
| slc26a5 | solute carrier family 26 (anion exchanger), | ENSDARG00000022424 | 0.95 | 1.21 | 1.62 | 0.71 | | | | |
| slc16a10 | solute carrier family 16 member 10 | ENSDARG00000020984 | 0.95 | 0.63 | 0.85 | 0.70 | | | | |
| slc25a44b | solute carrier family 25, member 44 b | ENSDARG00000035905 | 0.94 | 0.95 | 0.92 | 0.98 | | | | |
| slc25a1a | solute carrier family 25 (mitochondrial carrier; | ENSDARG00000057110 | 0.94 | 1.10 | 1.23 | 0.84 | | | | |
| slc6a1b | solute carrier family 6 (neurotransmitter | ENSDARG00000039647 | 0.93 | 0.78 | 0.74 | 0.98 | | | | |
| slc1a4 | solute carrier family 1 (glutamate/neutral amino | ENSDARG00000000551 | 0.90 | 0.68 | 0.98 | 0.63 | | | | |
| slc22a2 | solute carrier family 22 (organic cation | ENSDARG00000030530 | 0.89 | 1.56 | 1.66 | 0.84 | | | | |
| slc25a38b | solute carrier family 25, member 38b | ENSDARG00000074533 | 0.88 | 0.59 | 0.73 | 0.71 | | | | |
| slc6a19b | solute carrier family 6 (neutral amino acid | ENSDARG00000056719 | 0.87 | 3.33 | 0.32 | 9.20 | | | | |
| slc1a2b | solute carrier family 1 (glial high affinity | ENSDARG00000102453 | 0.87 | 0.80 | 0.94 | 0.75 | | | | |
| slc43a1a | solute carrier family 43 (amino acid system L | ENSDARG00000037393 | 0.85 | 0.70 | 0.76 | 0.79 | | | | |
| slc35e1 | solute carrier family 35, member E1 | ENSDARG00000011945 | 0.85 | 0.99 | 0.86 | 0.98 | | | | |
| slc29a2 | solute carrier family 29 (equilibrative nucleoside | ENSDARG00000001767 | 0.85 | 1.24 | 1.21 | 0.87 | | | | |
| slc7a6os | solute carrier family 7, member 6 opposite | ENSDARG00000010596 | 0.83 | 1.06 | 0.81 | 1.09 | | | | |
| slc25a32a | solute carrier family 25 (mitochondrial folate | ENSDARG00000089791 | 0.82 | 1.09 | 1.01 | 0.88 | | | | |
| slc24a4a | solute carrier family 24 | ENSDARG00000015425 | 0.82 | 1.55 | 1.14 | 1.11 | | | | |
| slc25a14 | solute carrier family 25 (mitochondrial carrier, | ENSDARG00000026680 | 0.81 | 1.13 | 0.95 | 0.97 | | | | |
| slc25a28 | solute carrier family 25 (mitochondrial iron | ENSDARG00000052994 | 0.80 | 1.22 | 1.09 | 0.90 | | | | |
| slc25a39 | solute carrier family 25, member 39 | ENSDARG00000007449 | 0.80 | 1.03 | 1.45 | 0.57 | | | | |
| slc39a14 | solute carrier family 39 (zinc transporter), | ENSDARG00000102387 | 0.79 | 1.03 | 0.35 | 2.36 | | | | |
| slc6a17 | solute carrier family 6 (neutral amino acid | ENSDARG00000068787 | 0.79 | 1.01 | 0.90 | 0.88 | | | | |
| slc37a2 | solute carrier family 37 (glucose-6-phosphate | ENSDARG00000023394 | 0.79 | 0.97 | 0.76 | 1.01 | | | | |
| slc32a1 | solute carrier family 32 (GABA vesicular | ENSDARG00000059775 | 0.79 | 1.00 | 0.81 | 0.97 | | | | |
| slc20a2 | solute carrier family 20 (phosphate transporter), | ENSDARG00000060796 | 0.79 | 1.27 | 1.02 | 0.98 | | | | |

Fold change
>5.0 = red
>3.0 = light red
>1.2 = pale red
<0.85 = pale blue
<0.65 = light blue
<0.5 = blue

p<0.05 p<0.1

Gene counts
>1
>10
>100
>1000

| Gene Symbol | Gene Title | ensemble | (1) DMSO (operated vs unoperated) | (2) Operated (SB431542 vs DMSO) | (3) Unoperated (SB431542 vs DMSO) | (4) SB431542 (operated vs unoperated) | DMSO unoperated COUNT | DMSO operated COUNT | SB43 unoperated COUNT | SB43 operated COUNT |
|---|---|---|---|---|---|---|---|---|---|---|
| slc22a7b.2 | solute carrier family 22 member 7-like | ENSDARG00000091252 | 0.78 | 0.68 | 0.24 | 2.27 | | | | |
| slc27a2a | solute carrier family 27 (fatty acid transporter), | ENSDARG00000036237 | 0.78 | 0.53 | 1.11 | 0.37 | | | | |
| slc7a6 | solute carrier family 7 member 6 | ENSDARG00000054423 | 0.78 | 1.20 | 1.60 | 0.59 | | | | |
| slc7a8a | solute carrier family 7 (cationic amino acid | ENSDARG00000075831 | 0.78 | 1.33 | 1.45 | 0.71 | | | | |
| slc25a12 | solute carrier family 25 (aspartate/glutamate | ENSDARG00000102362 | 0.77 | 0.97 | 1.26 | 0.59 | | | | |
| slc25a11 | solute carrier family 25 (mitochondrial carrier; | ENSDARG00000035741 | 0.77 | 0.86 | 0.90 | 0.73 | | | | |
| slc48a1a | solute carrier family 48 (heme transporter), | ENSDARG00000026907 | 0.75 | 2.03 | 1.54 | 1.00 | | | | |
| slc25a27 | solute carrier family 25, member 27 | ENSDARG00000042873 | 0.75 | 1.28 | 1.32 | 0.73 | | | | |
| slc25a5 | solute carrier family 25 (mitochondrial carrier; | ENSDARG00000092553 | 0.75 | 1.30 | 1.16 | 0.84 | | | | |
| slc7a3a | solute carrier family 7 (cationic amino acid | ENSDARG00000020645 | 0.75 | 2.39 | 2.02 | 0.88 | | | | |
| slc2a3a | solute carrier family 2 (facilitated glucose | ENSDARG00000013295 | 0.75 | 1.39 | 1.56 | 0.67 | | | | |
| slc2a2 | solute carrier family 2 (facilitated glucose | ENSDARG00000056196 | 0.72 | 1.19 | 0.98 | 0.87 | | | | |
| slc6a11a | solute carrier family 6 (neurotransmitter | ENSDARG00000074002 | 0.72 | 9.84 | 7.75 | 0.92 | | | | |
| slc1a3a | solute carrier family 1 (glial high affinity | ENSDARG00000104431 | 0.71 | 1.09 | 1.06 | 0.73 | | | | |
| slc38a5b | solute carrier family 38, member 5b | ENSDARG00000014587 | 0.71 | 0.81 | 0.78 | 0.74 | | | | |
| slc16a3 | solute carrier family 16 (monocarboxylate | ENSDARG00000045051 | 0.71 | 1.15 | 1.15 | 0.71 | | | | |
| slc25a3b | solute carrier family 25 (mitochondrial carrier; | ENSDARG00000025566 | 0.70 | 0.86 | 0.79 | 0.76 | | | | |
| slc25a25a | solute carrier family 25 (mitochondrial carrier; | ENSDARG00000010572 | 0.70 | 2.26 | 1.46 | 1.08 | | | | |
| slc18a3a | solute carrier family 18 (vesicular acetylcholine | ENSDARG00000006356 | 0.69 | 0.92 | 0.83 | 0.76 | | | | |
| slc25a55b | solute carrier family 25 member 55b | ENSDARG00000013749 | 0.68 | 0.74 | 0.83 | 0.60 | | | | |
| slc39a13 | solute carrier family 39 (zinc transporter), | ENSDARG00000000442 | 0.67 | 0.75 | 1.01 | 0.50 | | | | |
| slc16a12b | solute carrier family 16, member 12b | ENSDARG00000089885 | 0.65 | 1.14 | 0.94 | 0.78 | | | | |
| slc38a4 | solute carrier family 38, member 4 | ENSDARG00000018149 | 0.65 | 0.94 | 0.94 | 0.64 | | | | |
| slc43a2a | solute carrier family 43 (amino acid system L | ENSDARG00000036848 | 0.64 | 1.97 | 0.55 | 2.30 | | | | |
| slc35a4 | solute carrier family 35, member A4 | ENSDARG00000062379 | 0.63 | 1.17 | 0.85 | 0.87 | | | | |
| slc38a6 | solute carrier family 38, member 6 | ENSDARG00000054312 | 0.63 | 3.23 | 1.87 | 1.08 | | | | |
| slc3a2b | solute carrier family 3 (amino acid transporter | ENSDARG00000037012 | 0.62 | 1.75 | 0.71 | 1.53 | | | | |
| slc9a8 | solute carrier family 9, subfamily A (NHE8, | ENSDARG00000020699 | 0.62 | 1.19 | 0.98 | 0.76 | | | | |
| slc37a4b | solute carrier family 37 (glucose-6-phosphate | ENSDARG00000077180 | 0.59 | 0.99 | 0.80 | 0.74 | | | | |
| slc6a19a.1 | solute carrier family 6 (neutral amino acid | ENSDARG00000018621 | 0.59 | 2.23 | 1.79 | 0.74 | | | | |
| slc44a5b | solute carrier family 44 member 5b | ENSDARG00000057419 | 0.58 | 1.89 | 0.72 | 1.53 | | | | |
| slc25a51b | solute carrier family 25 member 51-like | ENSDARG00000040463 | 0.58 | 1.05 | 0.82 | 0.75 | | | | |
| slc25a36b | solute carrier family 25 member 36b | ENSDARG00000015915 | 0.57 | 1.18 | 0.86 | 0.78 | | | | |
| slc25a4 | solute carrier family 25 (mitochondrial carrier; | ENSDARG00000027355 | 0.54 | 0.97 | 0.85 | 0.61 | | | | |
| slc16a4 | solute carrier family 16 member 4 | ENSDARG00000042807 | 0.52 | 1.29 | 0.75 | 0.89 | | | | |
| slc8a4b | solute carrier family 8 (sodium/calcium | ENSDARG00000037145 | 0.49 | 1.49 | 0.63 | 1.16 | | | | |
| slc2a11l | solute carrier family 2 (facilitated glucose | ENSDARG00000062873 | 0.44 | 0.21 | 0.33 | 0.28 | | | | |
| slc13a1 | solute carrier family 13 (sodium/sulphate | ENSDARG00000045638 | 0.40 | 1.53 | 0.36 | 1.68 | | | | |
| slc5a12 | solute carrier family 5 | ENSDARG00000005004 | 0.40 | 6.00 | 1.29 | 1.85 | | | | |
| slc25a3a | solute carrier family 25 (mitochondrial carrier; | ENSDARG00000027424 | 0.35 | 2.76 | 0.69 | 1.38 | | | | |
| slc25a44a | solute carrier family 25, member 44 a | ENSDARG00000045927 | 0.16 | 3.92 | 0.62 | 1.03 | | | | |
| slc25a37 | solute carrier family 25 (mitochondrial iron | ENSDARG00000073743 | 0.12 | 2.31 | 0.50 | 0.54 | | | | |
| tgm2l | transglutaminase 2, like | ENSDARG00000093381 | 7.27 | 0.78 | 2.09 | 2.70 | | | | |
| tgm2b | transglutaminase 2b | ENSDARG00000074094 | 4.20 | 0.66 | 3.23 | 0.86 | | | | |
| tgm2a | transglutaminase 2, C polypeptide A | ENSDARG00000070157 | 0.34 | 0.79 | 1.19 | 0.22 | | | | |
| tgm1l4 | transglutaminase 1 like 4 | ENSDARG00000101407 | 0.27 | 0.68 | 0.25 | 0.74 | | | | |
| tnnt2a | troponin T type 2a (cardiac) | ENSDARG00000020610 | 4.17 | 0.21 | 1.00 | 1.00 | | | | |
| tnnt2c | troponin T2c, cardiac | ENSDARG00000032242 | 3.90 | 0.73 | 1.01 | 2.82 | | | | |
| tnni2b.1 | troponin I type 2b (skeletal, fast), tandem | ENSDARG00000035958 | 2.34 | 0.97 | 1.35 | 1.68 | | | | |
| tnni1b | troponin I type 1b (skeletal, slow) | ENSDARG00000052708 | 1.94 | 0.59 | 0.62 | 1.86 | | | | |
| tnnc1a | troponin C type 1a (slow) | ENSDARG00000011400 | 0.70 | 1.16 | 0.12 | 6.67 | | | | |

Fold change
>5.0 = red
>3.0 = light red
>1.2 = pale red
<0.85 = pale blue
<0.65 = light blue
<0.5 = blue

Gene counts
>1
>10
>100
>1000

p<0.05 p<0.1

| Gene Symbol | Gene Title | ensemble | (1) DMSO (operated vs unoperated) | (2) Operated (SB431542 vs DMSO) | (3) Unoperated (SB431542 vs DMSO) | (4) SB431542 (operated vs unoperated) | DMSO unoperated COUNT | DMSO operated COUNT | SB43 unoperated COUNT | SB43 operated COUNT |
|---|---|---|---|---|---|---|---|---|---|---|
| tnnc1b | troponin C type 1b (slow) | ENSDARG00000037539 | 0.65 | 0.90 | 0.84 | 0.69 | | | | |
| tnni4b.2 | troponin I4b, tandem duplicate 2 | ENSDARG00000036671 | 0.65 | 1.14 | 1.01 | 0.74 | | | | |
| tnni1d | troponin I, skeletal, slow d | ENSDARG00000073766 | 0.62 | 0.89 | 0.97 | 0.57 | | | | |
| tnnt3a | troponin T type 3a (skeletal, fast) | ENSDARG00000030270 | 0.60 | 0.91 | 0.77 | 0.71 | | | | |
| tnnc2 | troponin C type 2 (fast) | ENSDARG00000070835 | 0.56 | 1.22 | 0.99 | 0.69 | | | | |
| tnni2a.4 | troponin I type 2a (skeletal, fast), tandem | ENSDARG00000029069 | 0.56 | 1.12 | 0.91 | 0.68 | | | | |
| tnnt3b | troponin T type 3b (skeletal, fast) | ENSDARG00000068457 | 0.55 | 1.02 | 0.92 | 0.61 | | | | |
| tnni2a.3 | troponin I type 2a (skeletal, fast), tandem | ENSDARG00000013752 | 0.48 | 1.29 | 0.70 | 0.89 | | | | |
| tnnt2d | troponin T2d, cardiac | ENSDARG00000002988 | 0.43 | 0.98 | 0.84 | 0.51 | | | | |
| tnni1c | troponin I, skeletal, slow c | ENSDARG00000042559 | 0.32 | 0.74 | 0.60 | 0.39 | | | | |
| tnni2b.2 | troponin I type 2b (skeletal, fast), tandem | ENSDARG00000029995 | 0.31 | 1.63 | 0.89 | 0.58 | | | | |
| tnni4b.1 | troponin I4b, tandem duplicate 1 | ENSDARG00000092999 | 0.25 | 2.11 | 0.60 | 0.87 | | | | |
| tubb6 | tubulin, beta 6 class V | ENSDARG00000104801 | 5.22 | 0.64 | 1.20 | 2.78 | | | | |
| tuba1b | tubulin, alpha 1b | ENSDARG00000045367 | 2.64 | 0.73 | 0.71 | 2.68 | | | | |
| tubd1 | tubulin, delta 1 | ENSDARG00000058219 | 2.30 | 0.68 | 1.07 | 1.45 | | | | |
| tuba8l | tubulin, alpha 8 like | ENSDARG00000042708 | 1.86 | 0.67 | 0.69 | 1.81 | | | | |
| tuba8l4 | tubulin, alpha 8 like 4 | ENSDARG00000006260 | 1.58 | 0.75 | 0.83 | 1.43 | | | | |
| tubb4b | tubulin, beta 4B class IVb | ENSDARG00000002344 | 1.37 | 0.97 | 1.24 | 1.08 | | | | |
| tubb4b | tubulin, beta 4B class IVb | ENSDARG00000002344 | 1.37 | 0.97 | 1.24 | 1.08 | | | | |
| tubb2b | tubulin, beta 2b | ENSDARG00000098591 | 1.37 | 0.81 | 0.90 | 1.23 | | | | |
| tubg1 | tubulin, gamma 1 | ENSDARG00000015610 | 1.31 | 0.61 | 0.97 | 0.82 | | | | |
| tubgcp4 | tubulin, gamma complex associated protein 4 | ENSDARG00000005374 | 1.30 | 0.82 | 0.92 | 1.15 | | | | |
| tubgcp5 | tubulin, gamma complex associated protein 5 | ENSDARG00000077442 | 1.28 | 0.88 | 1.21 | 0.93 | | | | |
| tubgcp2 | tubulin, gamma complex associated protein 2 | ENSDARG00000013079 | 1.25 | 1.14 | 1.43 | 1.00 | | | | |
| tuba4l | tubulin, alpha 4 like | ENSDARG00000074289 | 1.22 | 1.09 | 0.93 | 1.43 | | | | |
| tuba7l | tubulin, alpha 7 like | ENSDARG00000104643 | 1.00 | 1.00 | 3.04 | 0.28 | | | | |
| tuba8l3 | tubulin, alpha 8 like 3 | ENSDARG00000070155 | 0.93 | 0.78 | 0.83 | 0.87 | | | | |
| tubb5 | tubulin, beta 5 | ENSDARG00000037997 | 0.91 | 1.67 | 1.31 | 1.15 | | | | |
| tubgcp3 | tubulin, gamma complex associated protein 3 | ENSDARG00000029133 | 0.89 | 2.13 | 2.08 | 0.91 | | | | |
| tuba2 | tubulin, alpha 2 | ENSDARG00000045014 | 0.79 | 1.11 | 1.29 | 0.68 | | | | |
| tubb2 | tubulin, beta 2A class IIa | ENSDARG00000039522 | 0.70 | 1.20 | 1.09 | 0.77 | | | | |
| tuba1a | tubulin, alpha 1a /// zgc:123298 | ENSDARG00000001889 | 0.67 | 0.89 | 0.79 | 0.75 | | | | |
| tuba1c | tubulin, alpha 1c | ENSDARG00000055216 | 0.59 | 1.00 | 0.92 | 0.64 | | | | |
| tuba8l2 | tubulin, alpha 8 like 2 | ENSDARG00000031164 | 0.45 | 1.17 | 0.94 | 0.56 | | | | |
| ube2c | ubiquitin-conjugating enzyme E2C | ENSDARG00000114670 | 2.33 | 0.88 | 0.97 | 2.11 | | | | |
| ube2d1b | ubiquitin-conjugating enzyme E2D 1b | ENSDARG00000038576 | 2.10 | 1.45 | 1.93 | 1.58 | | | | |
| ube2d2l | ubiquitin-conjugating enzyme E2D 2 (UBC4/5 | ENSDARG00000099749 | 1.72 | 0.70 | 0.95 | 1.27 | | | | |
| ube3d | ubiquitin protein ligase E3D | ENSDARG00000026178 | 1.69 | 0.88 | 0.94 | 1.58 | | | | |
| ube2q2 | ubiquitin-conjugating enzyme E2Q family | ENSDARG00000013990 | 1.60 | 2.18 | 2.16 | 1.62 | | | | |
| ube2d1a | ubiquitin-conjugating enzyme E2D 1a | ENSDARG00000029107 | 1.56 | 1.86 | 2.85 | 1.02 | | | | |
| ube2s | ubiquitin-conjugating enzyme E2S | ENSDARG00000031775 | 1.46 | 0.93 | 0.98 | 1.38 | | | | |
| ube2v2 | ubiquitin-conjugating enzyme E2 variant 2 | ENSDARG00000028198 | 1.36 | 1.02 | 0.84 | 1.64 | | | | |
| ube2l3b | ubiquitin-conjugating enzyme E2L 3b | ENSDARG00000027141 | 1.36 | 0.86 | 1.01 | 1.15 | | | | |
| ube4b | ubiquitination factor E4B, UFD2 homolog (S. | ENSDARG00000037017 | 1.29 | 0.76 | 0.90 | 1.09 | | | | |
| ube2na | ubiquitin-conjugating enzyme E2Na | ENSDARG00000008748 | 1.28 | 1.12 | 1.16 | 1.23 | | | | |
| ube2ia | ubiquitin-conjugating enzyme E2Ia | ENSDARG00000052649 | 1.25 | 1.17 | 1.47 | 0.99 | | | | |
| ube2r2 | ubiquitin-conjugating enzyme E2R 2 | ENSDARG00000058740 | 1.24 | 1.19 | 1.28 | 1.16 | | | | |
| ube2d2 | ubiquitin-conjugating enzyme E2D 2 (UBC4/5 | ENSDARG00000043484 | 1.24 | 0.96 | 0.91 | 1.30 | | | | |
| ube2g2 | ubiquitin-conjugating enzyme E2G 2 (UBC7 | ENSDARG00000025404 | 1.18 | 1.68 | 2.18 | 0.91 | | | | |
| ube2ib | ubiquitin-conjugating enzyme E2Ib | ENSDARG00000007438 | 1.15 | 0.72 | 0.78 | 1.06 | | | | |
| ube2kb | ubiquitin-conjugating enzyme E2Kb (UBC1 | ENSDARG00000013505 | 1.11 | 0.87 | 0.76 | 1.28 | | | | |

Fold change
>5.0 = red
>3.0 = light red
>1.2 = pale red
<0.85 = pale blue
<0.65 = light blue
<0.5 = blue
p<0.05 p<0.1

Gene counts
>1
>10
>100
>1000

| Gene Symbol | Gene Title | ensemble | (1) DMSO (operated vs unoperated) | (2) Operated (SB431542 vs DMSO) | (3) Unoperated (SB431542 vs DMSO) | (4) SB431542 (operated vs unoperated) | DMSO unoperated COUNT | DMSO operated COUNT | SB43 unoperated COUNT | SB43 operated COUNT |
|---|---|---|---|---|---|---|---|---|---|---|
| ube2g1a | ubiquitin-conjugating enzyme E2G 1a (UBC7 | ENSDARG00000015292 | 1.09 | 1.01 | 1.23 | 0.89 | | | | |
| ube2e3 | ubiquitin-conjugating enzyme E2E 3 (UBC4/5 | ENSDARG00000012244 | 1.07 | 1.02 | 1.13 | 0.97 | | | | |
| ube2ql1 | ubiquitin-conjugating enzyme E2Q family-like 1 | ENSDARG00000079276 | 1.05 | 1.56 | 0.82 | 2.00 | | | | |
| ube2e2 | ubiquitin-conjugating enzyme E2E 2 | ENSDARG00000034670 | 1.04 | 1.11 | 1.27 | 0.91 | | | | |
| ube2q1 | ubiquitin-conjugating enzyme E2Q family | ENSDARG00000100766 | 0.98 | 0.83 | 0.75 | 1.08 | | | | |
| ube2a | ubiquitin-conjugating enzyme E2A (RAD6 | ENSDARG00000098466 | 0.95 | 0.97 | 1.15 | 0.80 | | | | |
| ube2nb | ubiquitin-conjugating enzyme E2Nb | ENSDARG00000045877 | 0.87 | 0.91 | 0.91 | 0.87 | | | | |
| ube2h | ubiquitin-conjugating enzyme E2H (UBC8 | ENSDARG00000000019 | 0.80 | 0.86 | 0.98 | 0.71 | | | | |
| ube2d3 | ubiquitin-conjugating enzyme E2D 3 | ENSDARG00000038473 | 0.79 | 1.18 | 0.92 | 1.00 | | | | |
| ube2g1b | ubiquitin-conjugating enzyme E2G 1b (UBC7 | ENSDARG00000069527 | 0.69 | 1.15 | 0.95 | 0.83 | | | | |
| ube2d4 | ubiquitin-conjugating enzyme E2D 4 | ENSDARG00000015057 | 0.38 | 0.95 | 0.37 | 0.98 | | | | |
| ywhabb | tyrosine 3-monooxygenase/tryptophan 5- | ENSDARG00000075758 | 2.08 | 1.13 | 1.36 | 1.72 | | | | |
| ywhaqb | tyrosine 3-monooxygenase/tryptophan 5- | ENSDARG00000023323 | 1.85 | 1.54 | 1.26 | 2.27 | | | | |
| ywhaba | tyrosine 3-monooxygenase/tryptophan 5- | ENSDARG00000013078 | 1.57 | 0.98 | 1.15 | 1.33 | | | | |
| ywhaqa | tyrosine 3-monooxygenase/tryptophan 5- | ENSDARG00000042539 | 1.50 | 0.81 | 0.94 | 1.30 | | | | |
| ywhaz | tyrosine 3-monooxygenase/tryptophan 5- | ENSDARG00000032575 | 1.49 | 0.95 | 0.71 | 2.00 | | | | |
| ywhae2 | tyrosine 3-monooxygenase/tryptophan 5- | ENSDARG00000017014 | 1.25 | 0.92 | 0.93 | 1.24 | | | | |
| ywhah | tyrosine 3-monooxygenase/tryptophan 5- | ENSDARG00000005560 | 0.93 | 0.75 | 0.81 | 0.87 | | | | |
| ywhae1 | tyrosine 3-monooxygenase/tryptophan 5- | ENSDARG00000006399 | 0.82 | 0.87 | 0.67 | 1.07 | | | | |
| ywhabl | tyrosine 3-monooxygenase/tryptophan 5- | ENSDARG00000040287 | 0.81 | 1.07 | 0.80 | 1.09 | | | | |

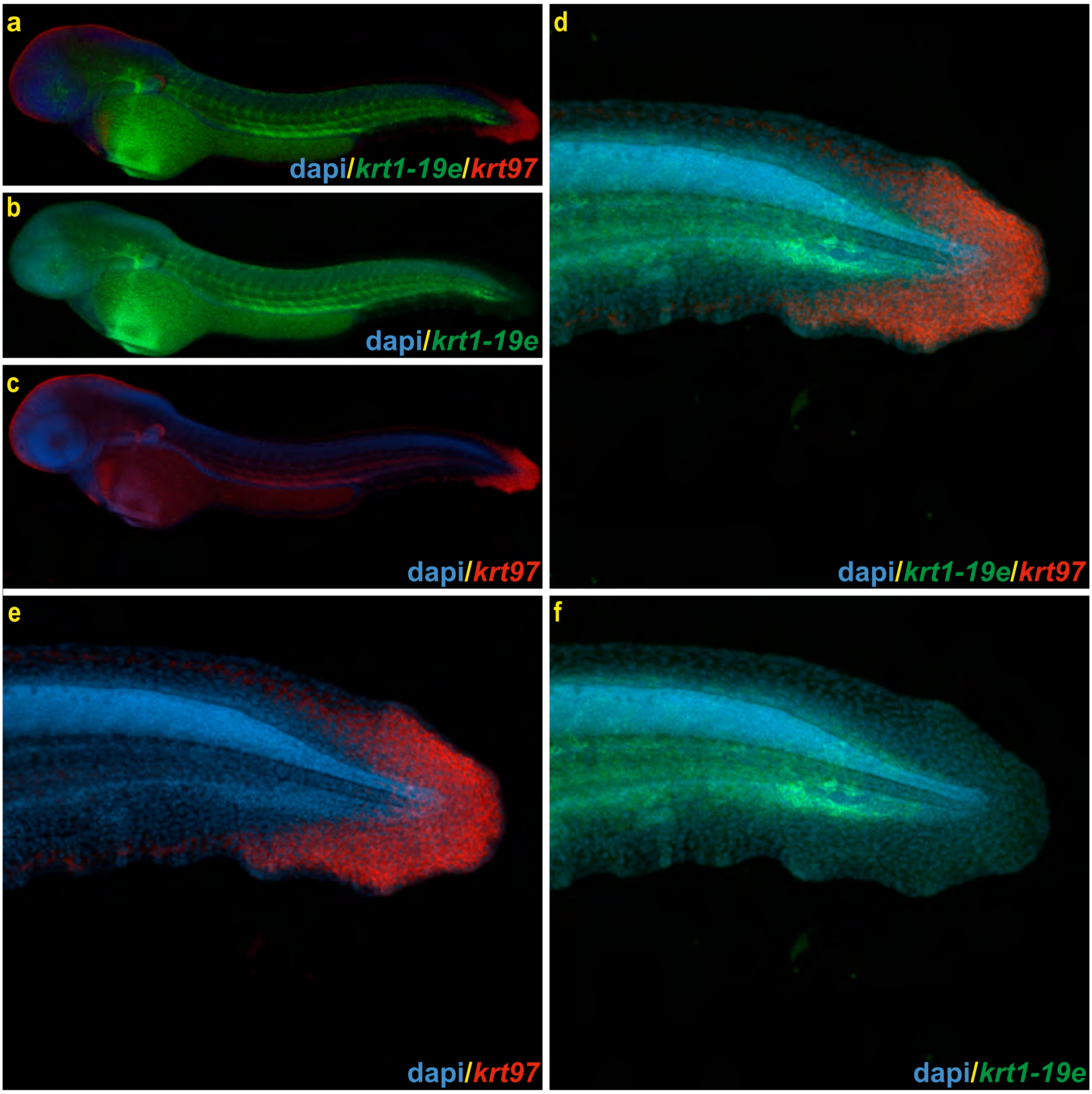


**Supplemental Figure 6: *krt1-19e* is expressed in the trunk region and does not overlap with *krt97* at 48hpf.** (a-c) Low magnification and (d-f) high magnification images of *krt1-19e* expression is shown in green, DAPI staining of nuclei in blue and *krt97* in red. *krt1-19e* is expressed in the trunk and does not overlap with *krt97* at this stage.

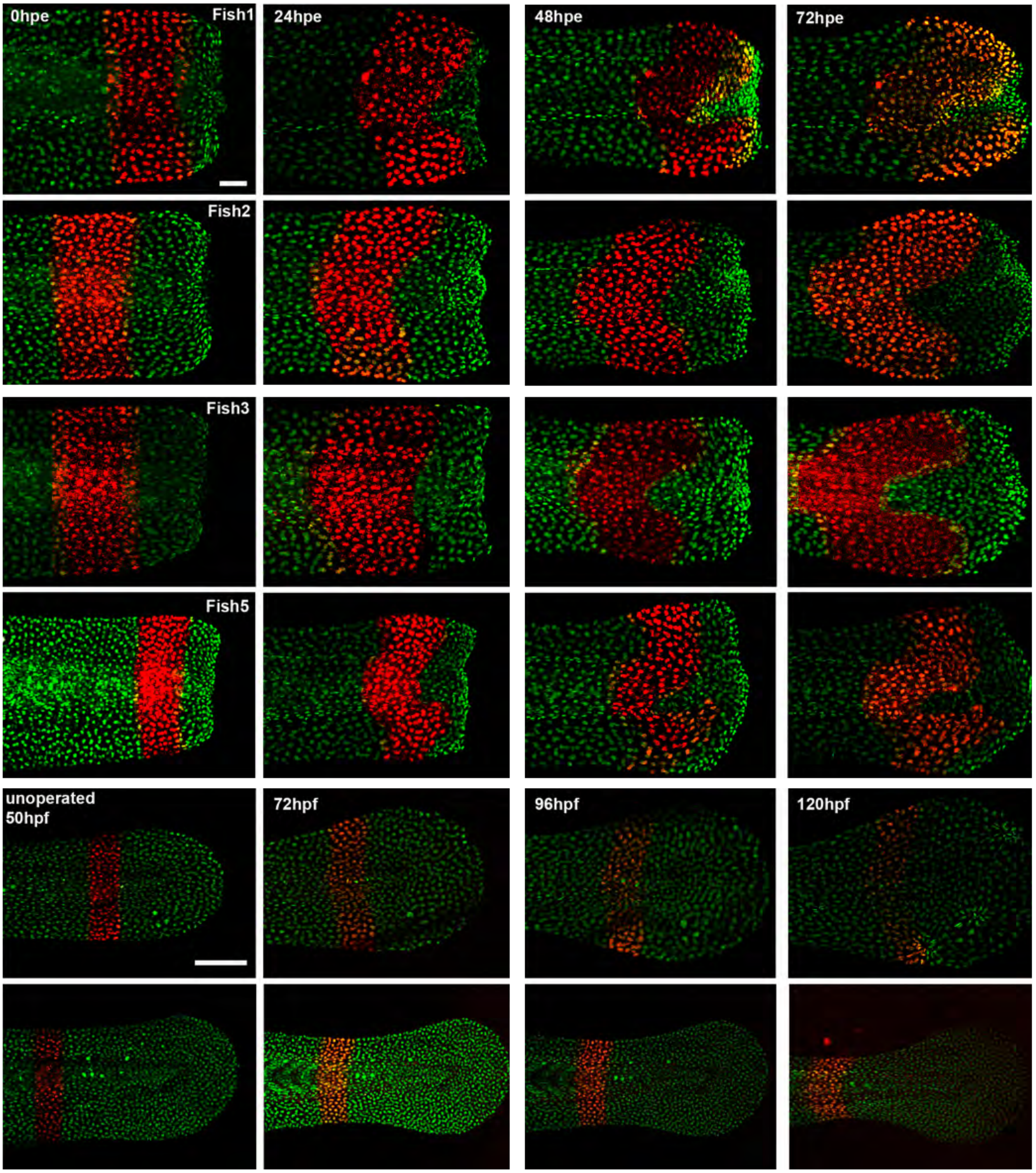


**Supplemental Figure 7:** Examples of photoconversion experiments during regeneration.

**Supplemental Figure 8. R packages.** This table lists the packages and versions used for the scRNA-seq analysis presented in the study.

| Package | Version |
|---|---|
| abind | 1.4-8 |
| askpass | 1.2.0 |
| assertthat | 0.2.1 |
| assorthead | 0.99.8 |
| backports | 1.5.0 |
| base64enc | 0.1-3 |
| beachmat | 2.21.6 |
| beeswarm | 0.4.0 |
| BH | 1.84.0-0 |
| Biobase | 2.65.1 |
| BiocGenerics | 0.51.1 |
| BiocManager | 1.30.25 |
| BiocParallel | 1.39.0 |
| BiocSingular | 1.21.3 |
| BiocVersion | 3.20.0 |
| bit | 4.5.0 |
| bit64 | 4.5.2 |
| bitops | 1.0-8 |
| blob | 1.2.4 |
| brew | 1.0-10 |
| brio | 1.1.5 |
| broom | 1.0.6 |
| bslib | 0.8.0 |
| cachem | 1.1.0 |
| Cairo | 1.6-2 |
| callr | 3.7.6 |
| car | 3.1-2 |
| carData | 3.0-5 |
| caTools | 1.18.3 |
| cellranger | 1.1.0 |
| circlize | 0.4.16 |
| cli | 3.6.3 |
| clipr | 0.8.0 |
| clock | 0.7.1 |
| clue | 0.3-65 |
| colorspace | 2.1-1 |
| commonmark | 1.9.1 |
| ComplexHeatmap | 2.21.0 |
| conflicted | 1.2.0 |
| corrplot | 0.94 |
| COTAN | 2.5.7 |
| cowplot | 1.1.3 |

| | |
|---|---|
| cpp11 | 0.5.0 |
| crayon | 1.5.3 |
| credentials | 2.0.1 |
| crosstalk | 1.2.1 |
| curl | 5.2.3 |
| data.table | 1.16.0 |
| DBI | 1.2.3 |
| dbplyr | 2.5.0 |
| DelayedArray | 0.31.11 |
| DelayedMatrixStats | 1.27.3 |
| deldir | 2.0-4 |
| dendextend | 1.17.1 |
| Deriv | 4.1.6 |
| desc | 1.4.3 |
| DESeq2 | 1.45.3 |
| devtools | 2.4.5 |
| diagram | 1.6.5 |
| dials | 1.3.0 |
| DiceDesign | 1.1 |
| diffobj | 0.3.5 |
| digest | 0.6.37 |
| doBy | 4.6.22 |
| doFuture | 1.0.1 |
| doParallel | 1.0.17 |
| dotCall64 | 1.1-1 |
| downlit | 0.4.4 |
| dplyr | 1.1.4 |
| dqrng | 0.4.1 |
| DropletUtils | 1.25.2 |
| dtplyr | 1.3.1 |
| edgeR | 4.3.15 |
| ellipsis | 0.3.2 |
| evaluate | 1.0.0 |
| fansi | 1.0.6 |
| farver | 2.1.2 |
| fastDummies | 1.7.4 |
| fastmap | 1.2.0 |
| fitdistrplus | 1.2-1 |
| FNN | 1.1.4.1 |
| fontawesome | 0.5.2 |
| forcats | 1.0.0 |
| foreach | 1.5.2 |
| formatR | 1.14 |
| fs | 1.6.4 |
| furrr | 0.3.1 |
| futile.logger | 1.4.3 |
| futile.options | 1.0.1 |
| future | 1.34.0 |

| | |
|---|---|
| future.apply | 1.11.2 |
| gargle | 1.5.2 |
| generics | 0.1.3 |
| GenomeInfoDb | 1.41.1 |
| GenomeInfoDbData | 1.2.12 |
| GenomicRanges | 1.57.1 |
| gert | 2.1.2 |
| GetoptLong | 1.0.5 |
| ggbeeswarm | 0.7.2 |
| ggplot2 | 3.4.4 |
| ggplotify | 0.1.2 |
| ggprism | 1.0.5 |
| ggpubr | 0.6.0 |
| ggrastr | 1.0.2 |
| ggrepel | 0.9.6 |
| ggridges | 0.5.6 |
| ggsci | 3.2.0 |
| ggsignif | 0.6.4 |
| ggthemes | 5.1.0 |
| gh | 1.4.1 |
| gitcreds | 0.1.2 |
| GlobalOptions | 0.1.2 |
| globals | 0.16.3 |
| glue | 1.7.0 |
| goftest | 1.2-3 |
| googledrive | 2.1.1 |
| googlesheets4 | 1.1.1 |
| gower | 1.0.1 |
| GPfit | 1.0-8 |
| gplots | 3.1.3.1 |
| gridExtra | 2.3 |
| gridGraphics | 0.5-1 |
| gtable | 0.3.5 |
| gtools | 3.9.5 |
| hardhat | 1.4.0 |
| haven | 2.5.4 |
| HDF5Array | 1.33.6 |
| hdf5r | 1.3.11 |
| here | 1.0.1 |
| highr | 0.11 |
| hms | 1.1.3 |
| htmltools | 0.5.8.1 |
| htmlwidgets | 1.6.4 |
| httpuv | 1.6.15 |
| httr | 1.4.7 |
| httr2 | 1.0.4 |
| ica | 1.0-3 |
| ids | 1.0.1 |

| | |
|---|---|
| igraph | 2.0.3 |
| ini | 0.3.1 |
| ipred | 0.9-15 |
| IRanges | 2.39.2 |
| irlba | 2.3.5.1 |
| isoband | 0.2.7 |
| iterators | 1.0.14 |
| janitor | 2.2.0 |
| jquerylib | 0.1.4 |
| jsonlite | 1.8.9 |
| kernlab | 0.9-33 |
| knitr | 1.48 |
| ks | 1.14.3 |
| labeling | 0.4.3 |
| lambda.r | 1.2.4 |
| later | 1.3.2 |
| latex2exp | 0.9.6 |
| lava | 1.8.0 |
| lazyeval | 0.2.2 |
| leiden | 0.4.3.1 |
| lhs | 1.2.0 |
| lifecycle | 1.0.4 |
| limma | 3.61.9 |
| listenv | 0.9.1 |
| lme4 | 1.1-35.5 |
| lmtest | 0.9-40 |
| locfit | 1.5-9.10 |
| lubridate | 1.9.3 |
| magrittr | 2.0.3 |
| markdown | 1.13 |
| MatrixGenerics | 1.17.0 |
| MatrixModels | 0.5-3 |
| matrixStats | 1.4.1 |
| mclust | 6.1.1 |
| memoise | 2.0.1 |
| microbenchmark | 1.5.0 |
| mime | 0.12 |
| miniUI | 0.1.1.1 |
| minqa | 1.2.8 |
| modelenv | 0.1.1 |
| modelr | 0.1.11 |
| multicool | 1.0.1 |
| munsell | 0.5.1 |
| mvtnorm | 1.3-1 |
| Nebulosa | 1.15.0 |
| nloptr | 2.1.1 |
| numDeriv | 2016.8-1.1 |
| openssl | 2.2.2 |

| | |
|---|---|
| openxlsx | 4.2.7.1 |
| paletteer | 1.6.0 |
| parallelDist | 0.2.6 |
| parallelly | 1.38.0 |
| parsnip | 1.2.1 |
| patchwork | 1.3.0 |
| pbapply | 1.7-2 |
| pbkrtest | 0.5.3 |
| PCAtools | 2.17.0 |
| pillar | 1.9.0 |
| pkgbuild | 1.4.4 |
| pkgconfig | 2.0.3 |
| pkgdown | 2.1.1 |
| pkgload | 1.4.0 |
| plotly | 4.10.4 |
| plyr | 1.8.9 |
| png | 0.1-8 |
| polyclip | 1.10-7 |
| polynom | 1.4-1 |
| pracma | 2.4.4 |
| praise | 1.0.0 |
| presto | 1.0.0 |
| prettyunits | 1.2.0 |
| prismatic | 1.1.2 |
| processx | 3.8.4 |
| prodlim | 2024.06.25 |
| profvis | 0.4.0 |
| progress | 1.2.3 |
| progressr | 0.14.0 |
| promises | 1.3.0 |
| ps | 1.8.0 |
| purrr | 1.0.2 |
| quantreg | 5.98 |
| R.methodsS3 | 1.8.2 |
| R.oo | 1.26.0 |
| R.utils | 2.12.3 |
| R6 | 2.5.1 |
| ragg | 1.3.3 |
| RANN | 2.6.2 |
| rappdirs | 0.3.3 |
| rcmdcheck | 1.4.0 |
| RColorBrewer | 1.1-3 |
| Rcpp | 1.0.13 |
| RcppAnnoy | 0.0.22 |
| RcppArmadillo | 14.0.2-1 |
| RcppEigen | 0.3.4.0.2 |
| RcppGSL | 0.3.13 |
| RcppHNSW | 0.6.0 |

| | |
|---|---|
| RcppParallel | 5.1.9 |
| RcppProgress | 0.4.2 |
| RcppTOML | 0.2.2 |
| RcppZiggurat | 0.1.6 |
| RCurl | 1.98-1.16 |
| readr | 2.1.5 |
| readxl | 1.4.3 |
| recipes | 1.1.0 |
| rematch | 2.0.0 |
| rematch2 | 2.1.2 |
| remotes | 2.5.0 |
| reprex | 2.1.1 |
| reshape2 | 1.4.4 |
| reticulate | 1.39.0 |
| Rfast | 2.1.0 |
| rhdf5 | 2.49.0 |
| rhdf5filters | 1.17.0 |
| Rhdf5lib | 1.27.0 |
| rjson | 0.2.23 |
| rlang | 1.1.4 |
| rmarkdown | 2.28 |
| ROCR | 1.0-11 |
| roxygen2 | 7.3.2 |
| rprojroot | 2.0.4 |
| rsample | 1.2.1 |
| RSpectra | 0.16-2 |
| rstatix | 0.7.2 |
| rstudioapi | 0.16.0 |
| rsvd | 1.0.5 |
| Rtsne | 0.17 |
| rversions | 2.1.2 |
| rvest | 1.0.4 |
| S4Arrays | 1.5.7 |
| S4Vectors | 0.43.2 |
| sass | 0.4.9 |
| ScaledMatrix | 1.13.0 |
| scales | 1.3.0 |
| scattermore | 1.2 |
| scCustomize | 2.1.2 |
| SCpubr | 2.0.2 |
| sctransform | 0.4.1 |
| scuttle | 1.15.4 |
| selectr | 0.4-2 |
| sessioninfo | 1.2.2 |
| Seurat | 5.1.0 |
| SeuratData | 0.2.2.9001 |
| SeuratObject | 5.0.2 |
| SeuratWrappers | 0.3.2 |

| | |
|---|---|
| sfd | 0.1.0 |
| shape | 1.4.6.1 |
| shiny | 1.9.1 |
| SingleCellExperiment | 1.27.2 |
| sitmo | 2.0.2 |
| slider | 0.3.1 |
| snakecase | 0.11.1 |
| snow | 0.4-4 |
| sourcetools | 0.1.7-1 |
| sp | 2.1-4 |
| spam | 2.10-0 |
| SparseArray | 1.5.37 |
| SparseM | 1.84-2 |
| sparseMatrixStats | 1.17.2 |
| spatstat.data | 3.1-2 |
| spatstat.explore | 3.3-2 |
| spatstat.geom | 3.3-3 |
| spatstat.random | 3.3-2 |
| spatstat.sparse | 3.1-0 |
| spatstat.univar | 3.0-1 |
| spatstat.utils | 3.1-0 |
| SQUAREM | 2021.1 |
| statmod | 1.5.0 |
| stringi | 1.8.4 |
| stringr | 1.5.1 |
| SummarizedExperimeı | 1.35.1 |
| sys | 3.4.2 |
| systemfonts | 1.1.0 |
| tensor | 1.5 |
| testthat | 3.2.1.1 |
| textshaping | 0.4.0 |
| tibble | 3.2.1 |
| tidyr | 1.3.1 |
| tidyselect | 1.2.1 |
| tidyverse | 2.0.0 |
| timechange | 0.3.0 |
| timeDate | 4041.11 |
| tinytex | 0.53 |
| tune | 1.2.1 |
| tzdb | 0.4.0 |
| UCSC.utils | 1.1.0 |
| umap | 0.2.10.0 |
| urlchecker | 1.0.1 |
| usethis | 3.0.0 |
| utf8 | 1.2.4 |
| uuid | 1.2-1 |
| uwot | 0.2.2 |
| vctrs | 0.6.5 |

| | |
|---|---|
| VennDiagram | 1.7.3 |
| vipor | 0.4.7 |
| viridis | 0.6.5 |
| viridisLite | 0.4.2 |
| vroom | 1.6.5 |
| waldo | 0.5.3 |
| warp | 0.2.1 |
| whisker | 0.4.1 |
| withr | 3.0.1 |
| workflows | 1.1.4 |
| xfun | 0.47 |
| xml2 | 1.3.6 |
| xopen | 1.0.1 |
| xtable | 1.8-4 |
| XVector | 0.45.0 |
| yaml | 2.3.10 |
| yardstick | 1.3.1 |
| yulab.utils | 0.1.7 |
| zeallot | 0.1.0 |
| zip | 2.3.1 |
| zlibbioc | 1.51.1 |
| zoo | 1.8-12 |
